# ChromDMM: A Dirichlet-Multinomial Mixture Model For Clustering Heterogeneous Epigenetic Data

**DOI:** 10.1101/2022.03.25.485838

**Authors:** Maria Osmala, Gökçen Eraslan, Harri Lähdesmäki

## Abstract

**Motivation:** Research on epigenetic modifications and other chromatin features at genomic regulatory elements elucidates essential biological mechanisms including the regulation of gene expression. Despite the growing number of epigenetic datasets, new tools are still needed to discover novel distinctive patterns of heterogeneous epigenetic signals at regulatory elements.

**Results:** We introduce ChromDMM, a product Dirichlet-multinomial mixture model for clustering genomic regions that are characterised by multiple chromatin features. ChromDMM extends the mixture model framework by profile shifting and flipping that can probabilistically account for inaccuracies in the position and strand-orientation of the genomic regions. Owing to hyper-parameter optimisation, ChromDMM can also regularise the smoothness of the epigenetic profiles across the consecutive genomic regions. With simulated data, we demonstrate that ChromDMM clusters, shifts, and strand-orients the profiles more accurately than previous methods. With ENCODE data, we show that the clustering of enhancer regions in the human genome reveals distinct patterns in several chromatin features. We further validate the enhancer clusters by their enrichment for transcriptional regulatory factor binding sites.

**Availability:** The software is available at https://github.com/MariaOsmala/ChromDMM

## 1 Introduction

Over a decade, the next-generation sequencing technologies have produced massive data amounts to quantify chromatin features, including nucleosomal histone modification locations, transcriptional regulatory factor binding sites and chromatin accessibility(Mardis *et al.*, 2007; Park, 2009; Boyle *et al.*, 2008). The chromatin feature signals are routinely formed as counts of aligned sequencing reads at consecutive non-overlapping genomic windows (or bins) along the entire genome or a short DNA stretch. These coverage signals at regulatory elements, such as enhancers, are often investigated to understand the underlying biological mechanisms in the regulation of gene expression. The signals can be visualised as heatmaps by aligning them within a genomic window centred at the loci (see Fig. 3 as an example). The average *aggregate patterns* of the coverage signals illustrated on top of the heatmap reveal the positional correlations and recurrent patterns in the signals. However, the set of analysed genomic regions can be biologically heterogeneous, in other words, they consist of multiple unknown subclasses, and the aggregate plot derived from all regions falsely displays the superposition of several different chromatin signatures. Consequently, we need a clustering method to reveal the subclasses.

**Figure 1:**
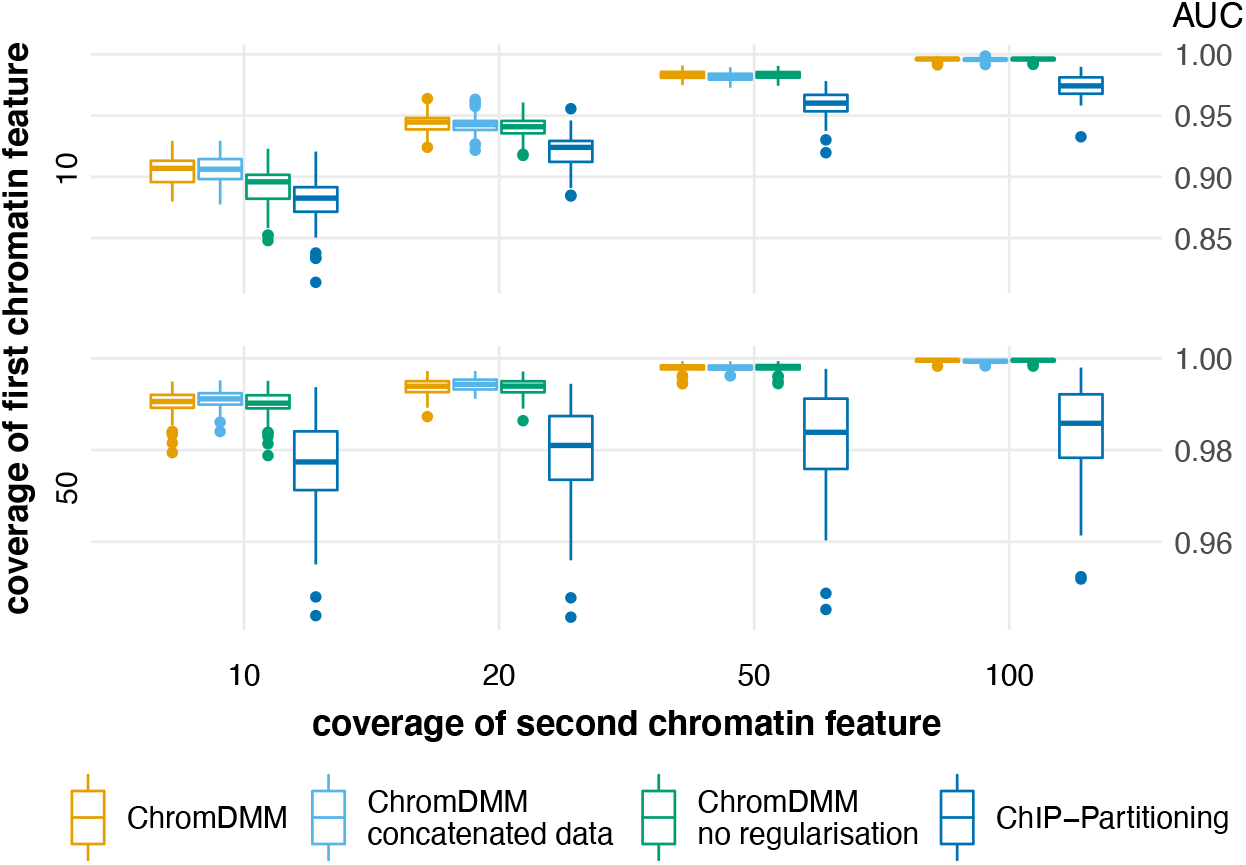
AUC values for clustering simulated data sets that contain two clusters and two chromatin features (H3K4me1 and RNA POL II). The chromatin feature coverages were varied between 10, 20, 50, and 100 for RNA POL II, and between 10 and 50 for H3K4me1 (see Suppl. Figure S8 for more combinations). Boxplots represent results for 100 data sets.

**Figure 2:**
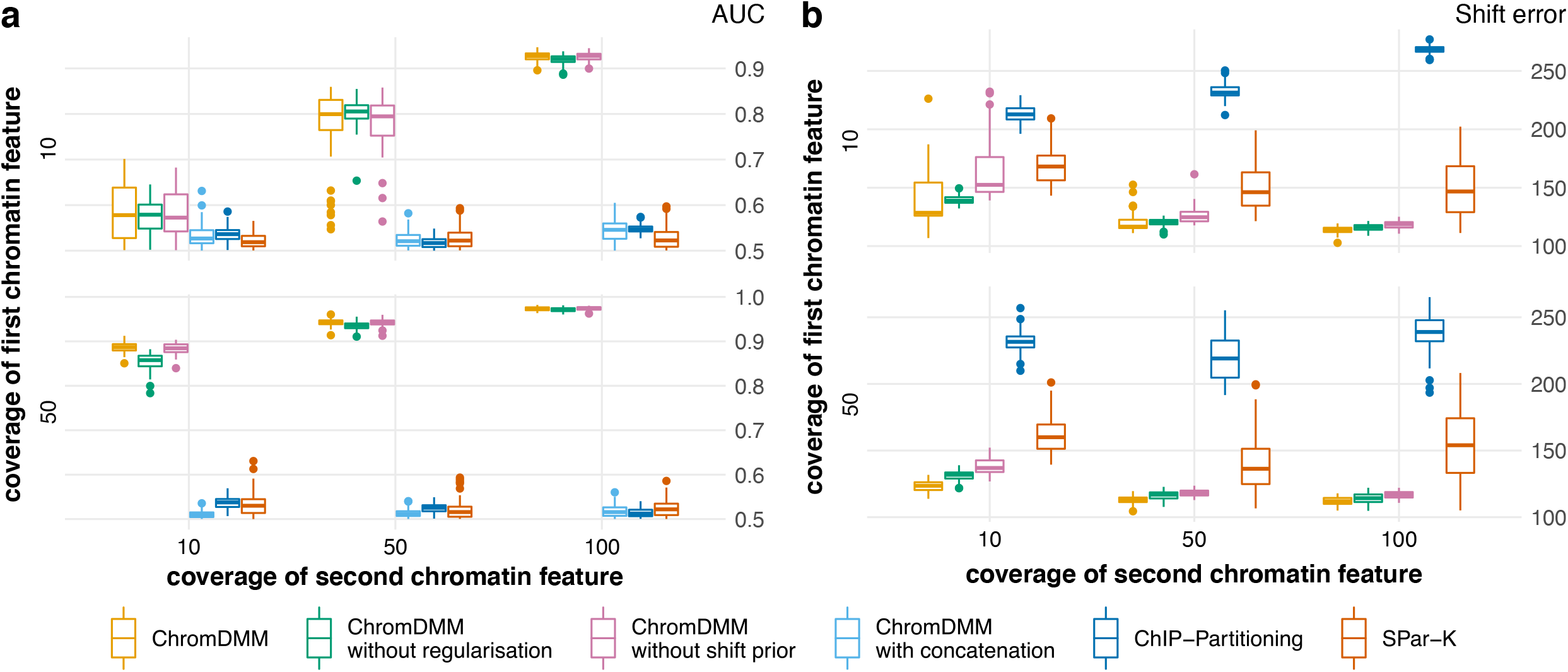
AUC values (a) and average shift errors in bps (b) for clustering simulated data that contain two clusters, two chromatin features and randomly sampled shift and flip states. The chromatin feature coverages varied between 10, 50, and 100. Results are shown for ChromDMM, ChIP-Partitioning, and SPar-K. For comparison, ChromDMM was applied also on concatenated chromatin features. ChromDMM was inferred also with the uniform shift state prior and without the regularisation term. Boxplots represent results for 100 data sets.

**Figure 3:**
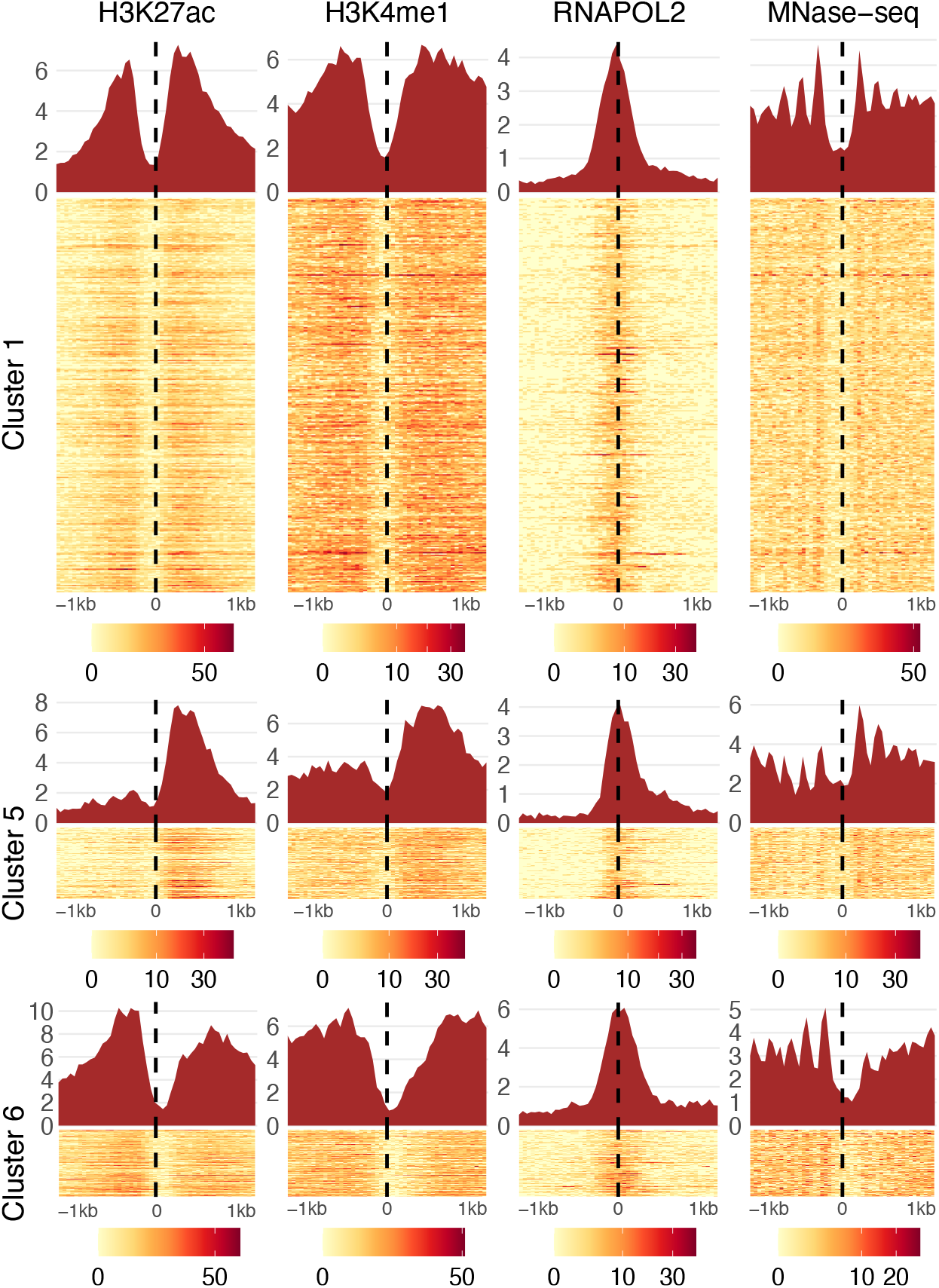
Enhancer clusters revealed by ChromDMM. The coverage signals of individual enhancers assigned to the clusters are visualised as heatmaps. The aggregate patterns are visualised on top of the heatmaps. Four of the ten chromatin features for three of the six clusters are shown. The full set of clusters and chromatin features resulting from enhancer clustering by ChromDMM are presented in Suppl. Figure S16

The clustering method must consider the following properties of the chromatin feature data. First, the data are heterogeneous containing sparse count data as well as varying coverage intensities and patterns. Second, the anchor positions of regulatory elements are typically uncertain; Genomic regions need to change, that is, the coverage signals need alignment with respect to each other to refine the aggregate patterns. Third, chromatin features can be asymmetric concerning the anchor points due to directional biomolecular mechanisms, such as transcription. Therefore, the coverage signals need strand-orientation (flipping).

Several methods have been proposed for the epigenetic data clustering, such as hierarchical clustering (Kundaje *et al.*, 2012) and *k*-means (Heintzman *et al.*, 2007; Ye *et al.*, 2010; Groux and Bucher, 2019). In addition, considering the above-mentioned requirements for the chromatin feature clustering method, Nair *et al.* (2014) introduced a probabilistic mixture model-based clustering method ChIP-Partitioning, which models the statistical variation in the coverage signals using independent Poisson distributions. Nair *et al.* (2014) further demonstrated that ChIP-Partitioning outperforms the hierarchical and k-means clustering methods, particularly when clustering low-coverage count data. However, next-generation sequencing data are typically overdispersed; the data variation is larger than expected by the Poisson distribution, and therefore many overdispersed models have been proposed, for example, for RNA-seq data analysis (Robinson *et al.*, 2010). In addition, previous studies on clustering chromatin features do not provide rigorous probabilistic methods for clustering multiple chromatin features simultaneously. The previous methods also lack a principled method for determining the unknown number of clusters.

We propose a probabilistic clustering method ChromDMM that exploits the discrete, sparse, non-negative and overdispersed nature of the sequencing data. ChromDMM builds on the mixture of Dirichlet-multinomial compound distributions originally proposed for clustering microbial data (Holmes *et al.*, 2012). We extend the model to account for the presence of multiple epigenetic coverage signals at the same genomic locus so that each mixture component exhibits a set of Dirichlet-multinomial compound distributions. We also extend the model with the profile shifting and flipping features that can probabilistically account for the inaccuracies in the positions and strand-orientations of the genomic regions being clustered. In addition, owing to the regularisation of the mixture component parameters, ChromDMM can smooth the chromatin feature patterns at successive bins along the genomic regions. Finally, our probabilistic model can naturally utilise the well-known model selection methods to determine the optimal number of clusters. Section 2 presents the ChromDMM model and its inference in detail. Section 3 analyses the performance of ChromDMM on simulated and real chromatin feature data and compares its performance with ChIP-Partitioning (Nair *et al.*, 2014) and SPar-K (Groux and Bucher, 2019).

## 2 Materials and Methods

The data for a chromatin feature across *N* genomic loci is represented as a *N* × *L* matrix **X** = [**x**_1_,…, **x**_*N*_]^*T*^, where **x**_*i*_ = [*x*_*i*1_,…, *x*_*iL*_]^*T*^ denotes the data for the *i*th genomic window. The length of **x**_*i*_ is defined by the size of the genomic locations *W* and resolution *B* as *L* = *W/B*. For example, data extracted in *W* = 2000 base pair (bp) windows centred at the anchor points with the resolution *B* = 40 bps results in coverage signals of length *L* = 50. The element *x*_*ij*_ denotes the number of sequencing reads whose starting position (5’ end) is aligned to bin *j* of locus *i*. To be exact, *x*_*ij*_ denotes, for example, the histone modification ChIP-seq read counts minus the sequencing-depth normalized control counts (see Suppl. Section S4.2). Collectively, the data for *M* chromatin features are represented as a *N* × *ML* matrix 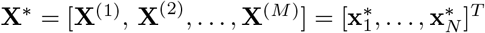, where 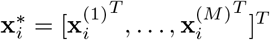 denotes a vector of length *ML* that contains the *M* chromatin feature vectors of a single genomic locus *i*.

### 2.1 Product Dirichlet-multinomial mixture model

The read counts **x** across the *L* bins are naturally modelled by the multinomial distribution with parameters 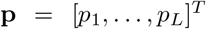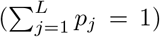. We further assume the multinomial parameters **p** are distributed according to a conjugate Dirichlet distribution with hyperparameters ***α***. Marginalising out the multinomial parameters from the joint distribution of **x** and **p** results in an overdispersed Dirichlet-multinomial compound distribution parameterised by ***α***. The Dirichlet-multinomial distributions can be utilised as the component distributions in a mixture model for the probabilistic clustering.

Compared to the standard Dirichlet-multinomial mixture model (Holmes *et al.*, 2012), we implement two extensions. First, we assume that the likelihood of **x**^*^ is a product multinomial distribution, each multinomial with the chromatin-feature-specific parameters **p**^(*m*)^. This enables modelling several chromatin features simultaneously. Second, we assume the parameters **p**^*^ = (**p**^(1)^,…, **p**^(*M*)^) to have a mixture prior with *K* mixture components; each component *k* is a product of *M* Dirichlet distributions again with the chromatin-feature-specific hyperparameters 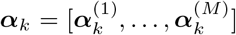. Let the parameters of the product-Dirichlet mixture for all *K* mixture components and *M* chromatin features be represented as a *L* × *KM* matrix 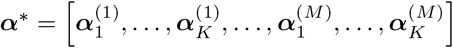. The mixture prior for **p**^*^ is

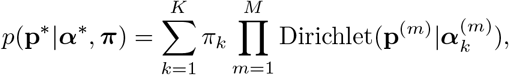

where ***π*** = (*π*_1_,…, *π*_*K*_) denotes the mixture weights. Holmes *et al.* (2012) showed that compounding the multinomial distribution with the Dirichlet mixture prior results in an analytically tractable likelihood. Similarly, in the case of the product-multinomial with the product-Dirichlet mixture prior, the parameters of the product-multinomial can also be marginalised analytically to derive a closed form expression for the likelihood of **X**^*^ as (see Suppl. Section S1.4 for a detailed derivation)

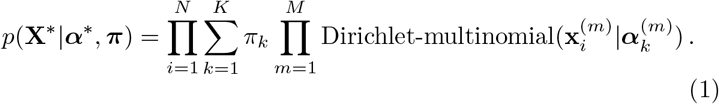

Instead of seeking to obtain the maximum-likelihood estimates for the model parameters, we adopt the Bayesian approach by introducing a prior distribution for the component parameters ***α***^*^.

To account for the correlations between the (expected) read counts at consecutive bins along the chromatin signal, we define a regularised Gamma hyperprior for the mixture component parameters ***α***^*^ as

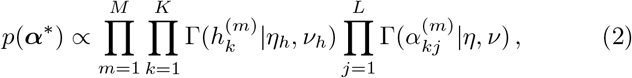

where all 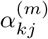 have their own independent Gamma prior with fixed shape *η* and rate *ν* parameters, and the regularisation terms

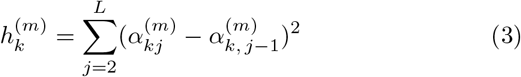

also have their own independent Gamma prior with shape *η*_*h*_ and rate *ν*_*h*_ parameters. Inclusion of the regulatory terms in the prior favours smooth mixture component parameters. A more detailed expression of the proportional distribution of the prior *p*(***α***^*^) is shown in Suppl. Equation S9, and an example of the effect of the regularisation is demonstrated in Suppl. Section S1.6.

Mixture models involve the latent cluster membership variables **z**; each observed 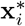 is associated with a corresponding unobserved categorical latent variable **z**_*i*_. The variable **z**_*i*_ = (*z*_*i*1_,…, *z*_*iK*_)^*T*^ is a *K*-dimensional indicator vector: if sample *i* originates from cluster *k*, *z*_*ik*_ = 1; otherwise *z*_*ik*_ = 0. The variables **z**_*i*_ are collected in a *N* × *K* matrix **Z** = [**z**_1_*,…,* **z**_*N*_]^*T*^. The proposed model is parameterised by ***θ*** = (***α***^*^, ***π***) and is presented as a directed acyclic graph in Suppl. Figure S1 together with the distributions of individual components.

### 2.2 The expectation-maximisation algorithm

The posterior log *p*(**X**^*^|***θ***) + log *p*(***θ***) can not be maximised directly. Instead, the MAP estimates for ***θ*** and the probabilistic cluster assignments are obtained by an iterative approach, the expectation-maximisation (EM) algorithm (Bishop, 2006). For the derivation of the EM algorithm assume a distribution for **Z**, *q*(**Z**). Then, the Jensen’s inequality provides a lower bound for the posterior distribution

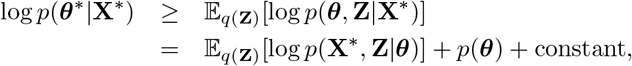

where log *p*(**X**^*^, **Z**|***θ***) is the complete data log-likelihood, and the constant term is independent on ***θ***. Assuming some initial estimates for the parameters ***θ***^old^ and defining *q*(**Z**) = *p*(**Z**|**X**^*^, ***θ***^old^), the lower bound (without the constant term) is

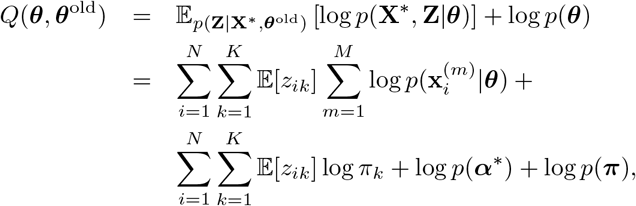

where the expectation is wrt. the posterior probabilities of the cluster assignments conditional on the current parameter estimates ***θ***^old^, i.e., 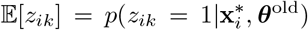. The likelihood term 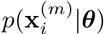 is the Dirichlet-multinomial compound distribution for the *m*th chromatin feature. For a detailed derivation, see Suppl. Section S1.8.

In the EM algorithm, E-steps and an M-steps are repeated, until convergence of the lower bound *Q*(***θ***, ***θ***^old^). In the E-step, the posterior probability that a sample *i* belongs to a cluster *k* given the current parameter estimates ***θ***^old^ is obtained using the standard Bayes rule as

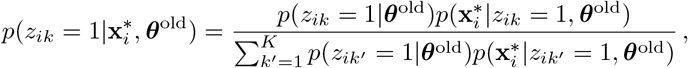

where 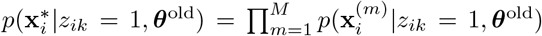 is the likelihood of the sample *i* condiotioned with cluster *k*, i.e., the product Dirichlet-multinomial distribution. The term *p*(*z*_*ik*_ = 1|***θ***^old^) corresponds to the mixture weight *π*_*k*_. In the M-step, as a closed-form solution of ***α***^*^ that maximises *Q*(***θ***, ***θ***^old^) is unattainable, the lower bound is maximised wrt. ***α***^*^ using Broyden-Fletcher-Goldfarb-Shanno (BFGS) method provided in R (Broyden, 1970). In addition, the component parameters 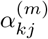 are constrained to be positive by a reparameterisation 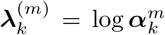 and by re-defining the prior for ***λ***^*^ accordingly using the multivariate change of variables method. For more details on deriving the equations for the model inference, see Suppl. Sections S1.7 – S1.15. The steps of the EM algorithm are summarised in Algorithm **??**. In the initilisation, the cluster membership probabilities 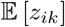 are obtained with the soft *k*-means clustering (MacKay, 2003) on concatenated chromatin features (see details in Suppl. Section S4.1). For a given number of clusters, the EM algorithm is run multiple times each with different random initialisation. Note that it is trivial to parallelise the computation across the multiple runs as well as across varying numbers of clusters.

**Algorithm 1:**
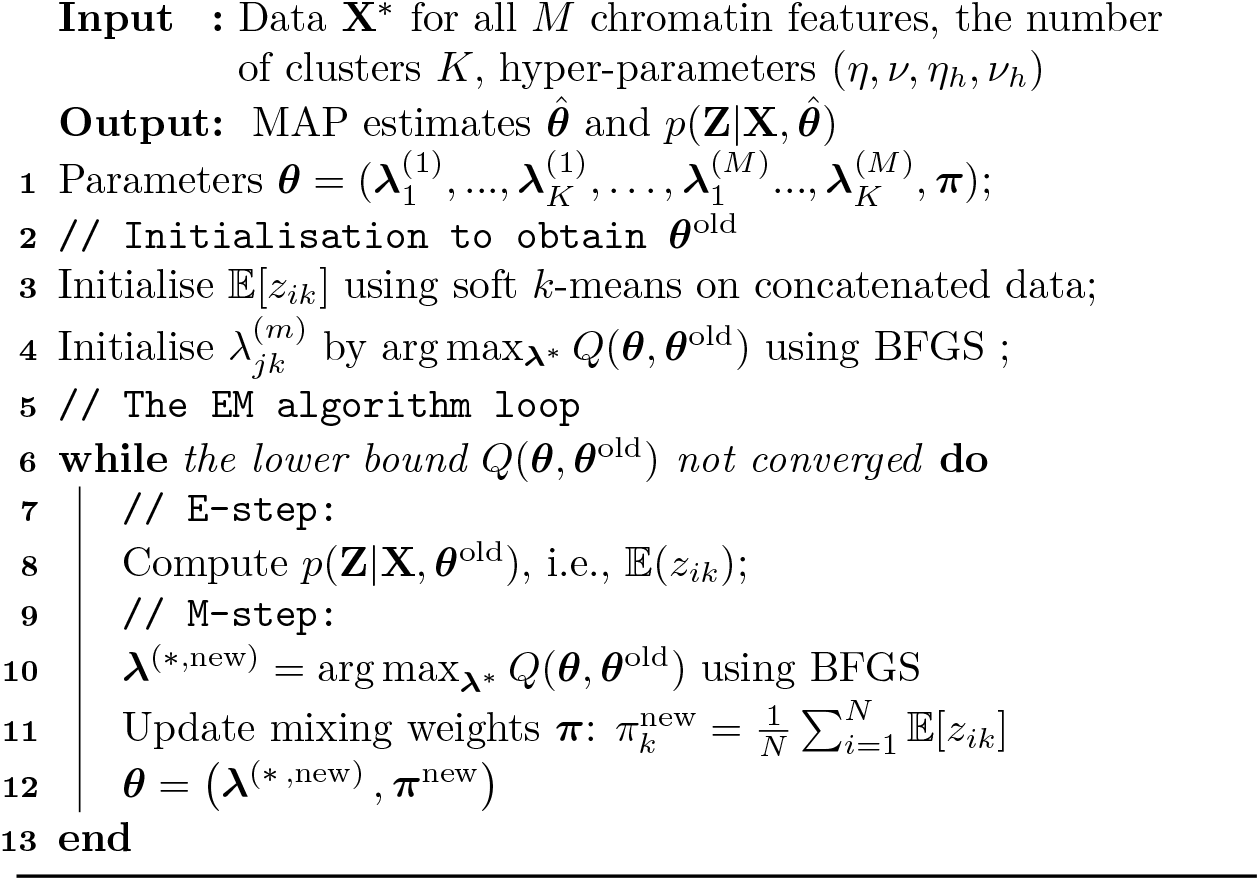
EM algorithm for ChromDMM

### 2.3 Chromatin feature profile shifting and flipping

We extend the product Dirichlet-multinomial mixture model with shifting and flipping features. For profile shifting, we first define the maximum amount of shifting, e.g. 400bp, both upstream and downstream. With a given bin size (e.g. *B* = 40bp), this results in 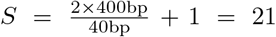 possible shift states, where the shift state 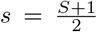 corresponds to no shift. In addition, the length of the Dirichlet parameters 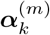 are extended from *L* to *L* + *S* – 1. When evaluating the likelihood model from Equation 1 for a shift state *s*, we use the corresponding *L*-length subset of the extended Dirichlet parameters for each mixture component *k*, denoted as 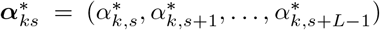. For profile ipping, we either compute the likelihood model definition with a shift state *s* (using again the *L*-length subset of the Dirichlet parameters) if *f* = 1, or reverse the order of the Dirichlet parameters if *f* = 2. Formally, we denote the shifting and flipping-aware likelihood model for a single genomic locus as 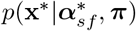.

For each locus, we can define prior probabilities for the shift and flip states. The prior shift state probabilities for the genomic locus *i* are denoted as ***ξ***_*i*_ = (*ξ*_*i*1_,…, *ξ*_*iS*_), where 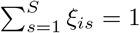. If the genomic loci and their anchor points are defined using ChIP-seq summits, then the prior for shift states can be defined, for example, as a pyramid-shaped prior that has the highest probability at the ChIP-seq peak summit (corresponding to no-shift state) and linearly decreasing the prior to zero beyond the maximum shift state. Similarly, we can define prior flip state probabilities ***ζ***_*i*_ for each locus *i*, where *ζ*_*i*1_ + *ζ*_*i*2_ = 1.

In ChromDMM with the shifting and flipping features, the latent cluster membership variables are re-defined as follows: *z*_*iksf*_ = 1 if the sample *i* originates from the cluster *k*, has shift state *s*, and has strand-orientation *f*; otherwise *z*_*iksf*_ = 0. These latent variables are stored in *N* × *K* × *S* × 2 matrix (or tensor) **Z**. We show in Suppl. Section S2.3 that the EM algorithm can be derived similarly as in Section 2.2, resulting in the following lower bound for the posterior distribution of parameters ***θ***

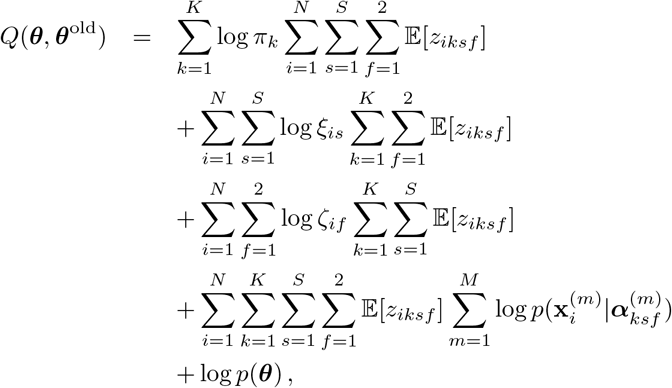

where 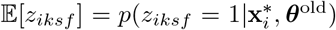. Note that the above mixture model can be applied *(i)* only with shifting, *(ii)* only with flipping, or *(iii)* with both shifting and flipping. In case of *(i)*, we simply drop the index *f* and the corresponding sums, and in case of *(ii)*, we simply drop the index *s* and the corresponding sums. For more details on the derivations of equations needed to infer the shifting and flipping-aware model, see Suppl. Section S2.

After learning the model parameters with the EM algorithm, we infer the final cluster assignment 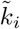 for each sample *i* by marginalizing the shift and flip states. Similarly, we choose the final flip and shift states, 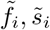, that maximise the posterior given the optimal cluster 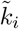 by marginalizing the shift and flip states, respectively:

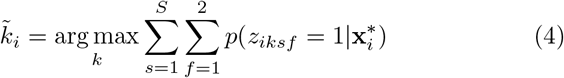

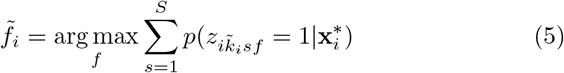

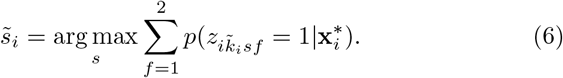

### 2.4 Choosing the number of clusters and identifiability aspects

For probabilistic clustering methods, the Bayesian model selection is commonly used to guide the selection of an appropriate number of clusters *K*. While the exact computation of the marginal likelihood is impractical, we can directly apply the commonly used approximative methods, such as the Bayesian information criterion (BIC) (Schwarz, 1978) or the Akaike information criterion (AIC) (Akaike, 1973).

There are inherent unidentifiability issues in ChromDMM results. Firstly, as in any clustering method, the inferred cluster labels can be switched between two clusters without affecting the clustering accuracy. Secondly, unless informative prior for strand-orientation is provided, the flip state indexes (1 or 2) can always be reversed. For biological interpretation, the aligned and flipped profiles need to be visualised after clustering and compared to the underlying directionality of the genomic region, such as direction of transcription 3’ → 5’ or 5’  → 3’, if known. The learned shift state is also affected by the learned flip state. While evaluating the performance of ChromDMM and other methods on simulated data, we consider these aspects.

## 3 Results

### 3.1 Clustering simulated data

#### 3.1.1 Data simulation and choice for hyperparameters

We used simulated data to investigate the clustering accuracy of ChromDMM, ChIP-Partitioning and SPar-K when the data contain varying number of chromatin features and varying read coverages. Briefly, we first clustered data for four chromatin features (H3K4me1, H3K27ac, RNA polymerase II, and MNase-seq) at 1000 enhancers from the ENCODE project (The ENCODE Project Consortium, 2012) by ChromDMM requiring the inference of both the shift and flip states. For more details, see Suppl. Methods S4.2. From the fitted model, we chose the Dirichlet parameters 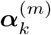 for two clusters. These parameters were used to sample the multinomial parameters 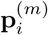 and finally the profiles 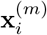 by varying the chromatin-feature-specific coverage between 10, 20, 50, and 100. For each experiment, we simulated 100 data sets. We used the area under receiver operations characteristics curve (AUC) as the performance measure for the clustering accuracy. For more details and the visualisation of a simulated data set with a coverage of 100, see Suppl. Section S4.4.

We performed hyperparameter sweeps on the simulated data to determine robust default values for the ChromDMM model. Briefly, for the Gamma prior for the Dirichlet parameters 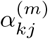 (parameterised by hyperparameters *η* and *ν*), we observed that the results were not sensitive to hyperparameter values and conclude that the choice of *η* = 1.1 and *ν* = 0.1 results in a good performance (Suppl. Figure S5). We also performed the prior predictive checks using the ancestral sampling of the data from prior hyperparameters and demonstrated that the amount of variation generated from the prior is comparable to the variation in the real data (Suppl. Figure S6). Similarly, we chose the hyperparameter values for the regularisation term 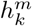. The Gamma prior with mean 1 and variance 0.1 corresponding to hyper-parameters *η*_*h*_ = *ν*_*h*_ = 10 resulted in a robust clustering performance (see Suppl. Figure S7 and Suppl. Section S4.5 for more details). We set the above hyperparameter values as defaults, but a user can e.g. perform prior predictive checks for his/her data and adjust the hyperparameters, if necessary.

We investigated the ability of AIC and BIC to choose the correct number of clusters (two) for the simulated data. We fitted ChromDMM with varying the number of clusters (from one to three). The proportions of cluster numbers selected by AIC and BIC in 100 simulated data sets are presented in Suppl. Figure S14a for 1000 samples and in Suppl. Figure S14b for 6000 samples. We conclude that AIC and BIC detect the correct number of clusters more reliably when the coverage of the chromatin modifications and/or the number of samples increases, although BIC tends to underestimate the number of clusters. The computation times of the three methods (with default parameters) to cluster simulated data containing two clusters and two chromatin features both with coverage 100 were: half an hour (ChromDMM), ca. 10 mins (ChIP-Partitioning), and seconds (SPar-K).

#### 3.1.2 ChromDMM infers accurate clusters

We compared ChrommDMM with ChIP-Partitioning and SPar-K (both applied with the default parameters) in clustering simulated data that contained two clusters and two chromatin features (H3K4me1 and RNA POL II). The clustering performance of ChromDMM exceeded the performance of ChIP-Partitioning (Figure 1 and Suppl. Figure S8). SPar-K performed poorly in these comparisons, particularly when the coverages were low (Suppl. Figure S9). Similar results were obtained on data containing only a single feature (Suppl. Figure S10). For comparison, Figure 1 and Suppl. Figure S8 present also results for an experiment where ChromDMM was fitted either on concatenated chromatin profile data or without the regularisation term. The regularisation improved the clustering performance especially when the coverage for the first chromatin feature (H3K4me1) was low, whereas the use of non-concatenated chromatin profile data resulted in only a marginal improvement in this simulation setting.

#### 3.1.3 Clustering accuracy improves with the number of features

We experimented with the number of chromatin features; beginning from a single feature (H3K4me1), the number of features was increased to four by adding RNA Pol II, H3K27ac and MNase-seq. Suppl. Figure S11a shows how the clustering accuracy increases together with the number of chromatin features for a signal coverage of 10. Figure S11b presents similar results for a varying coverage, where the coverage of the first set of chromatin features was 10 (H3K4m1, H3K27ac) and the coverage of the second set of chromatin features was 50 (RNA Pol II, MNase-seq). We conclude that for ChromDMM and ChIP-Partitioning, the clustering performance increases as a function of the number of chromatin features, whereas for SPar-K the improvement is less consistent. Regardless of the number of chromatin features, ChromDMM obtains the best performance.

#### 3.1.4 ChromDMM infers accurate shift and flip states

The simulated data were also artificially shifted and flipped as described in Suppl. Section S4.4. Briefly, the random shifts were constrained to be multiple of the data resolution (*B* = 40bp) and drawn from the Skellam distribution with mean zero and a variance that included the randomly sampled shifts between 400bp and +400bp. Similarly, the flip states were sampled randomly with equal probability for both strand-orientations. For more details, see Suppl. Algorithm S3.

The clustering accuracy of ChromDMM, ChIP-Partitioning and SPar-K was demonstrated on the randomly shifted and flipped simulated data. We experimented with four versions of ChromDMM: *(i)* ChromDMM with the regularisation term and with the pyramid-shaped shift state prior, *(ii)* same as *(i)* but with concatenated chromatin features, *(iii)* ChromDMM without the regularisation term and with the pyramid-shaped shift prior, and *(iv)* ChromDMM with a uniform prior for the shift states and with the regularisation term. The clustering accuracies of the methods are presented in Figure 2a. Methods considering the concatenated chromatin features, including ChIP-Partitioning and SPar-K, performed poorly in these comparisons, and notably they failed to improve their performance while increasing the coverage values. ChromDMM outperformed other methods, and its clustering performance was further improved by both the informative shift state prior and the regularisation, particularly when the coverage of either chromatin feature was low.

The methods were compared to their accuracy to correctly infer the the shift and the flip states of the genomic regions. ChromDMM and ChIP-Partitioning infer the most probable shift and flip states for each sample *i* from the latent variable probabilities shown in Equations 5 and 6, respectively, whereas SPar-K outputs the inferred shift and flip states separately. The flip state error was defined as the proportion of incorrectly inferred flip states in a given experiment (recall the identifiability aspects from Sec. 2.4). Similarly, the shift error for each sample was computed as the absolute difference between the true shift and the inferred shift in nucleotides. The average shift error over all *N* samples was reported as the final shift error for a single experiment.

The flip errors for the simulated shifted and flipped data containing two chromatin features and two clusters are presented in Suppl. Figure S12. On average, the flip errors decreased as the coverages increased, and they were lower for the ChromDMM methods compared to ChIP-Partitioning and SPar-K. The flip errors were only slightly affected by whether the ChromDMM fit was inferred without the shift prior or without the regularisation. The resulting shift errors for the simulated data are shown in Figure 2b. Again, the average shift errors decreased as the coverages increased, and the shift errors were lowest for the ChromDMM methods. The shift state inference of ChromDMM was further improved by the informative shift prior and the regularisation of the mixture component parameters ***α***^*^. In contrast to the other methods, the shift errors for ChIP-Partitioning remained high even with large coverage values. This likely results from the cluster-assigned patterns drifting from the profile center positions, i.e., ChIP-Partition selects a profile whose unimodal peak or valley between the two-modal peak is shifted far from the profile center, and aligns the rest of the profiles according to this single profile (Suppl. Figure S13d). The cluster patterns inferred by SPar-K also drift (Suppl. Figure S13e), whereas ChromDMM centers the peaks and valleys to the profile centers (Suppl. Figure S13c). This desirable behaviour of ChromDMM stems partly from the robustness of the probabilistic treatment and the shift state prior.

Finally, we investigated the ability of AIC and BIC to choose the correct number of clusters (two) in the simulated shifted and flipped data (Suppl. Figure S15). In contrast to the simpler model studied in Section 3.1, the more complex ChromDMM model with a high number of parameters is heavily penalised by AIC and BIC, and thus require higher coverage and larger number of samples to detect the correct number of clusters.

### 3.2 Clustering enhancers in ENCODE data

#### 3.2.1 ChromDMM reveals distinctive enhancer clusters

We applied ChromDMM, ChIP-Partitioning, and SPar-K with the flip and shift state inference to cluster ENCODE data containing 10 chromatin features extracted at enhancer regions. For the details of the data, preprocessing, and the definition of the enhancers, see Suppl. Section S4.2 and (Osmala and Lähdesmäki, 2020). Based on the ChromDMM fit, the enhancers were assigned to the most probably clusters, and their profiles were re-aligned based on the inferred shift and flip states, e.g. for visualisation. As a result, ChromDMM separated enhancers into six clusters, each with distinctive and refined combinations of chromatin feature patterns. Three of the six clusters are visualised as heatmaps and aggregated patterns in Figure 3 (the full set of clusters and chromatin features are presented in Suppl. Figure S16). In contrast, ChIP-Partitioning and SPar-K failed to identify distinctive patterns and to refine the profile alignment and strand-orientation (Suppl. Figures S18 and S19).

The ChromDMM enhancer clusters possess characteristic combinations of chromatin feature pattern shapes, spacings and signal strengths. The first cluster has symmetric and high enrichment of histone modification and MNase-seq signals with a steep decline of the signals in the middle of the profiles, indicating a nucleosome-free region. In addition, the nucleosome-free region is surrounded by a regular array of well-positioned nucleosomes. In contrast, in the clusters two, three, and four the nucleosome-free region and the well-positioning of the nucleosomes are obscured compared to the other clusters. Thus, the enhancers in these clusters may possess closed chromatin or mobile nucleosomes. The clusters four, five and six have asymmetricity in histone modification enrichment (cluster four and five), in nucleosome positioning (clusters five and six), and in RNA POL II occupancy (clusters four and five). The asymmetricity in the RNA POL II ChIP-seq signal may reflect the direction of transcription. In addition, in the asymmetric clusters, the histone modifications are enriched on either of the two nucleosomes immediately flanking the anchor position (cluster five) or spread widely (clusters four and six).

#### 3.2.2 Biological validation of the inferred clusters

The enhancer clusters revealed by ChromDMM, ChIP-Partitioning, and SPar-K were investigated for the enrichment of the binding sites of transcription factors (TFs) and other regulatory proteins, collectively referred to as transcriptional regulatory factors (TRFs). The ChIP-seq peaks for 220 TRFs were downloaded from ENCODE. For each TRF-cluster pair, a significance test for the enrichment of a given TRF at the cluster was performed by the GAT tool (Heger *et al.*, 2013). A large majority of the enrichments were significant according to the q-value threshold 0.01. To reveal differences in the TRF enrichment between clusters, the fold enrichments were visualised as a heatmap, where the enrichments corresponding to q-value larger than 0.01 were masked out (see Suppl. Figure S17 for ChromDMM clusters). The fold enrichments for TRFs which were significantly enriched in at least one ChromDMM cluster and simultaneously not enriched in at least one another cluster are presented in Figure 4. For more details, see Suppl. Methods S4.3.

**Figure 4:**
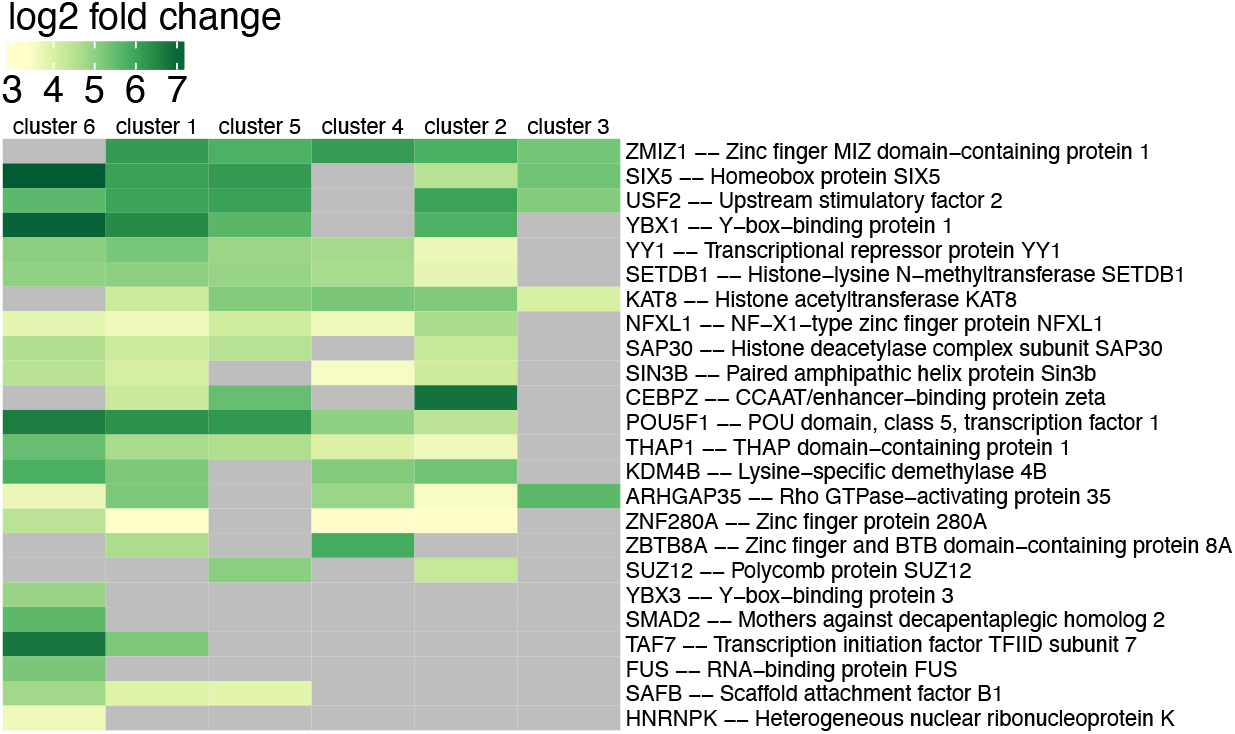
The fold enrichment of TRFs at the enhancer clusters identified by ChromDMM. The fold enrichments correponding to q-value larger than 0.01 were masked out from the heatmap. The enrichment results for all 220 TRFs are presented in Suppl. Figure S17.

The enrichment of TRFs in ChromDMM enhancer clusters reveals the potential biological significance of the distinctive chromatin feature patterns. Clusters three and four with obscured nucleosome-free regions and nucleosome positioning have less enrichment of TRFs than the four other clusters. In contrast, the first cluster with symmetric and strong signals has an enrichment for a large number of TRFs. Similarly to cluster one, cluster five with strong asymmetry in the histone modification and RNA POL II signals has a high TRF enrichment. Asymmetric cluster six with strong average H3K27ac, H3K9ac, and DNase-seq signals differs from the other clusters with unique enrichment for RNA binding and processing related proteins (HNRNPK, FUS) and transcription factors SMAD2 and YBX3. In addition, clusters one and six have enrichment for the largest component and core scaffold of the TFIID basal transcription factor complex (TAF7). Moreover, the clusters one, five and six are enriched for Scaffold attachment factor B1 (SAFB), a protein that binds DNA regions that are bound to the nuclear scaffold. Interestingly, SAFB may be involved in attaching the base of the chromatin loops to the nuclear scaffold and serving as a molecular base to assemble a transcriptosome complex in the vicinity of the actively transcribed genes (Nayler *et al.*, 1998). For comparison, the TRF enrichments at ChIP-Partitioning and SPar-K clusters are visualised in Suppl. Figures S20 and S21.

## 4 Conclusions

Exploring epigenetic data sets provides crucial information on key biological mechanisms such as gene regulation. An example is the clustering of epigenomic signals and other chromatin features at regulatory elements, such as enhancers, to reveal the combinations of chromatin features with varying signal magnitudes and profile shapes. To appropriately account for the sparse, discrete, heterogenous and overdispersed nature of the chromatin feature data, probabilistic clustering methods have been developed.

We proposed ChromDMM, a product Dirichlet-multinomial mixture model that provides a probabilistic method to cluster multiple chromatin feature coverage signals extracted from the same locus. Employing simulated data, we demonstrated that the accuracy of ChromDMM increases with the increasing number of chromatin features, indicating the need for a principled approach that considers the multiple chromatin features simultaneously when clustering regulatory elements. Moreover, we demonstrated that ChromDMM outperforms the previous methods ChIP-Partitioning and SPar-K in clustering accuracy, particularly when the chromatin feature coverages are low. In addition, ChromDMM learns the shift and flip states more accurately compared to ChIP-Partitioning and SPar-K. The accuracy of ChromDMM to infer the clusters and shift states are further improved by mixture component parameter regularisation and an informative shift state prior. Finally, we confirmed that BIC and AIC can detect the correct number of clusters.

We demonstrated that ChromDMM identifies clusters with distinct epigenetic patterns when applied to ENCODE data containing ten chromatin features quantified at enhancers. Moreover, the identified clusters are enriched for different sets of transcriptional regulatory factors, suggesting that the clusters vary in their biological characteristics.

## Funding and Acknowledgements

This work has been supported by the Academy of Finland (314445, 311584), and the Finnish Cultural Foundation. We thank Dr. Romain Groux for providing an implementation of the ChIP-Partitioning method. We acknowledge computer resources provided by the Aalto University School of Science “Science-IT” project.

## Notes

### Competing Interest Statement

The authors have declared no competing interest.

## References

Akaike, H. (1973). Information theory and an extension of the maximum likelihood principle. In B. Petrov and F. Csaki, editors, Second International Symposium on Information Theory, pages 267–281, Budabest, Hungary. Akademiai Kiado.

Bishop, C. M. (2006). Pattern Recognition and Machine Learning (Information Science and Statistics). Springer-Verlag, Berlin, Heidelberg.

Boyle, A. P., Davis, S., Shulha, H. P., Meltzer, P., Margulies, E. H., Weng, Z., Furey, T. S., and Crawford, G. E. (2008). High-resolution mapping and characterization of open chromatin across the genome. Cell, 132(2), 311–322.

Broyden, C. G. (1970). The convergence of a class of double-rank minimization algorithms 1. general considerations. IMA Journal of Applied Mathematics, 6(1), 76–90.

Groux, R. and Bucher, P. (2019). SPar-K: a method to partition NGS signal data. Bioinformatics, 35(21), 4440–4441.

Heger, A., Webber, C., Goodson, M., Ponting, C. P., and Lunter, G. (2013). GAT: a simulation framework for testing the association of genomic intervals. Bioinformatics, 29(16), 2046–2048.

Heintzman, N. D., Stuart, R. K., Hon, G., Fu, Y., Ching, C. W., Hawkins, R. D., Barrera, L. O., Van Calcar, S., Qu, C., Ching, K. A., et al. (2007). Distinct and predictive chromatin signatures of transcriptional promoters and enhancers in the human genome. Nature genetics, 39(3), 311–318.

Holmes, I., Harris, K., and Quince, C. (2012). Dirichlet multinomial mixtures: generative models for microbial metagenomics. PLoS One, 7(2), 1–15.

Kundaje, A., Kyriazopoulou-Panagiotopoulou, S., Libbrecht, M., Smith, C. L., Raha, D., Winters, E. E., Johnson, S. M., Snyder, M., Batzoglou, S., and Sidow, A. (2012). Ubiquitous heterogeneity and asymmetry of the chromatin environment at regulatory elements. Genome research, 22(9), 1735–1747.

MacKay, D. J. (2003). Information theory, inference, and learning algorithms. Cambridge University Press, New York, USA.

Mardis, E. R. et al. (2007). ChIP-seq: welcome to the new frontier. Nature methods, 4(8), 613–613.

Nair, N. U., Kumar, S., Moret, B. M., and Bucher, P. (2014). Probabilistic partitioning methods to find significant patterns in ChIP-Seq data. Bioinformatics, 30(17), 2406–2413.

Nayler, O., Strätling, W., Bourquin, J.-P., Stagljar, I., Lindemann, L., Jasper, H., Hartmann, A. M., Fackelmayer, F. O., Ullrich, A., and Stamm, S. (1998). SAF-B protein couples transcription and pre-mRNA splicing to SAR/MAR elements. Nucleic acids research, 26(15), 3542–3549.

Osmala, M. and Lähdesmäki, H. (2020). Enhancer prediction in the human genome by probabilistic modelling of the chromatin feature patterns. BMC bioinformatics, 21(1), 1–37.

Park, P. J. (2009). ChIP–seq: advantages and challenges of a maturing technology. Nature Reviews Genetics, 10(10), 669–680.

Robinson, M. D., McCarthy, D. J., and Smyth, G. K. (2010). edgeR: a Bioconductor package for differential expression analysis of digital gene expression data. Bioinformatics, 26(1), 139–140.

Schwarz, G. (1978). Estimating the dimension of a model. The annals of statistics, 6(2), 461–464.

The ENCODE Project Consortium (2012). An integrated encyclopedia of DNA elements in the human genome. Nature, 489(7414), 57–74.

Ye, T., Krebs, A. R., Choukrallah, M.-A., Keime, C., Plewniak, F., Davidson, I., and Tora, L. (2010). seqMINER: an integrated ChIP-seq data interpretation platform. Nucleic Acids Research, 39(6), e35.

