## Supplementary Information for "ChromDMM: A Dirichlet-Multinomial Mixture Model For Clustering Heterogeneous Epigenetic Data"

Maria Osmala, Gökçen Eraslan and Harri Lähdesmäki

**S5 Supplementary References 39**

**S6 Supplementary Figures 41**

### S1 Dirichlet-multinomial mixture model

#### S1.1 Components and a directed acyclic graph of the model

The proposed model is presented as a directed acyclic graph in Suppl. Figure S1 together with distributions of individual components. Individual genomic loci or samples are indexed by $i = 1, \ldots, N$, chromatin features are indexed by $m = 1, \ldots, M$, clusters are indexed by $k = 1, \ldots, K$ and the elements (bins) of coverage profiles are indexed by $j = 1, \ldots, L$. The data for $M$ chromatin features of a single locus $i$ is represented as a (length of $LM$) vector $\mathbf{x}_i^{(*)}$. The likelihood of $\mathbf{x}_i^{(*)}$ is presented as Equation 8 in Suppl. Figure S1. The likelihood states that the data at genomic loci $\mathbf{x}_i^{(*)}$, $i = 1, \ldots, N$, are sampled from Dirichlet-multinomial mixtures of size $K$ with component parameters $\boldsymbol{\alpha}_k^*$ and mixture weights $\pi_k$, $k = 1, \ldots, K$. The Dirichlet-multinomial parameters for all $K$ clusters and $M$ chromatin features are represented as a $(L \times KM)$ matrix $\boldsymbol{\alpha}^* = \left[\boldsymbol{\alpha}_1^{(1)}, \ldots, \boldsymbol{\alpha}_K^{(1)}, \ldots, \boldsymbol{\alpha}_1^{(M)}, \ldots, \boldsymbol{\alpha}_K^{(M)}\right]$ (Equation 1). The prior $p(\boldsymbol{\alpha}^*)$ is proportional to independent Gamma distributions (Equation 2), and depends on the regularisation term $h_k^{(m)}$ (Equation 3). The latent binary cluster memberships are presented as the $(N \times K)$ matrix $\mathbf{Z}$ . The categorical prior for the cluster memberships $p(\mathbf{z}_i|\boldsymbol{\pi})$ is presented in Equation 5. The mixing weights $p(\boldsymbol{\pi})$ are assumed uniform a priori (Equation 4).

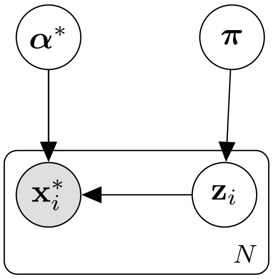

$N$: number of samples (genomic loci)

$M$: number of chromatin features

$K$: number of clusters

$L$: length of $\mathbf{x}_i^{(m)}$ and $\boldsymbol{\alpha}_k^{(m)}$

$$\boldsymbol{\alpha}^* = \left[\boldsymbol{\alpha}_1^{(1)}, \ldots, \boldsymbol{\alpha}_K^{(1)}, \ldots, \boldsymbol{\alpha}_1^{(M)}, \ldots, \boldsymbol{\alpha}_K^{(M)}\right] \tag{1}$$

$$p(\boldsymbol{\alpha}^*) \propto \prod_{m=1}^{M} \prod_{k=1}^{K} \Gamma(h_k^{(m)}|\eta_h, \nu_h) \prod_{j=1}^{L} \Gamma(\alpha_{kj}^{(m)}|\eta, \nu) \tag{2}$$

$$h_k^{(m)} = \sum_{j=2}^{L} (\alpha_{kj}^{(m)} - \alpha_{k,\,j-1}^{(m)})^2 \tag{3}$$

$$p(\pi_k) \propto 1\,, \sum_{k=1}^{K} \pi_k = 1 \tag{4}$$

$$p(\mathbf{z}_i|\boldsymbol{\pi}) = \prod_{k=1}^{K} \pi_k^{z_{ik}} \tag{5}$$

$$p(\mathbf{x}_i^*|\mathbf{z}_i, \boldsymbol{\alpha}^*) = \prod_{k=1}^{K} \prod_{m=1}^{M} \left[\text{Dirichlet-Multinomial}\left(\mathbf{x}_i^{(m)}|\boldsymbol{\alpha}_k^{(m)}\right)\right]^{z_{ik}} \tag{6}$$

$$p(\mathbf{x}_i^*, \mathbf{z}_i|\boldsymbol{\alpha}^*, \boldsymbol{\pi}) = \prod_{k=1}^{K} \prod_{m=1}^{M} \left[\pi_k \text{Dirichlet-Multinomial}\left(\mathbf{x}_i^{(m)}|\boldsymbol{\alpha}_k^{(m)}\right)\right]^{z_{ik}} \tag{7}$$

$$p(\mathbf{x}_i^*|\boldsymbol{\alpha}^*, \boldsymbol{\pi}) = \sum_{k=1}^{K} \pi_k \prod_{m=1}^{M} \text{Dirichlet-Multinomial}\left(\mathbf{x}_i^{(m)}|\boldsymbol{\alpha}_k^{(m)}\right) \tag{8}$$

Figure S1: Model diagram in a directed acyclic graph notation.

#### S1.2 Multinomial distribution and product multinomial distribution

The elements of the chromatin feature profile vector $\mathbf{x}_i^{(m)} = (x_{i1}^{(m)}, \ldots, x_{iL}^{(m)})$ are assumed to follow the multinomial distribution

$$p(\mathbf{x}_i^{(m)}|\mathbf{p}_i^{(m)}) = \Gamma\left(J_i^{(m)} + 1\right) \prod_{j=1}^{L} \frac{\left(p_{ij}^{(m)}\right)^{x_{ij}^{(m)}}}{\Gamma\left(x_{ij}^{(m)} + 1\right)}\,, \text{ where } J_i^{(m)} = \sum_{j=1}^{L} x_{ij}^{(m)}. \tag{S1}$$

We assume that the chromatin features are independent of each other. Hence, the likelihood of $\mathbf{x}_i^{(*)}$ is the product of chromatin-feature-specific multinomial distributions:

$$p(\mathbf{x}_i^{(*)}|\mathbf{p}_i^{(*)}) = \prod_{m=1}^{M} p(\mathbf{x}_i^{(m)}|\mathbf{p}_i^{(m)})\,. \tag{S2}$$

#### S1.3 Product Dirichlet mixture prior

The multinomial parameters $\mathbf{p}_i^{(*)}$ are assumed to be generated a priori by a mixture of product Dirichlet distributions:

$$\begin{aligned} p(\mathbf{p}_i^{(*)}|\boldsymbol{\alpha}^{(*)}, \boldsymbol{\pi}) &= \sum_{k=1}^{K} \pi_k \prod_{m=1}^{M} \text{Dirichlet}(\mathbf{p}_i^{(m)}|\boldsymbol{\alpha}_k^{(m)}) \\ &= \sum_{k=1}^{K} \pi_k \prod_{m=1}^{M} \frac{\Gamma\left(\sum_{j=1}^{L} \alpha_{kj}^{(m)}\right)}{\prod_{j=1}^{L} \Gamma\left(\alpha_{kj}^{(m)}\right)} \prod_{j=1}^{L} \left[p_{ij}^{(m)}\right]^{\alpha_{kj}^{(m)}-1}\,. \end{aligned} \tag{S3}$$

In Equation S3, each mixture component is a product of $M$ Dirichlet distributions, as the multinomial parameters for the $M$ separate chromatin features are assumed independent of each other.

#### S1.4 Product Dirichlet-multinomial likelihood

The product multinomial likelihood for $M$ chromatin features compounded by a mixture of product Dirichlet prior can be written as

$$\begin{aligned} p(\mathbf{x}_i^*|\boldsymbol{\alpha}^*, \boldsymbol{\pi}) &= \int_{\mathbf{p}_i} p(\mathbf{x}_i^*|\mathbf{p}_i^*) p(\mathbf{p}_i^*|\boldsymbol{\alpha}^*, \boldsymbol{\pi})\, \mathrm{d}\mathbf{p}_i^* \\ &= \int_{\mathbf{p}_i^*} \prod_{m=1}^{M} \text{Multinomial}(\mathbf{x}_i^{(m)}|\mathbf{p}_i^{(m)}) \times \sum_{k=1}^{K} \pi_k \prod_{m=1}^{M} \text{Dirichlet}(\mathbf{p}_i^{(m)}|\boldsymbol{\alpha}_k^{(m)})\, \mathrm{d}\mathbf{p}_i^* \\ &= \int_{\mathbf{p}_i^*} \sum_{k=1}^{K} \pi_k \prod_{m=1}^{M} \text{Multinomial}(\mathbf{x}_i^{(m)}|\mathbf{p}_i^{(m)}) \times \text{Dirichlet}(\mathbf{p}_i^{(m)}|\boldsymbol{\alpha}_k^{(m)})\, \mathrm{d}\mathbf{p}_i^* \\ &= \sum_{k=1}^{K} \pi_k \prod_{m=1}^{M} \underbrace{\int_{\mathbf{p}_i^{(m)}} \text{Multinomial}(\mathbf{x}_i^{(m)}|\mathbf{p}_i^{(m)}) \times \text{Dirichlet}(\mathbf{p}_i^{(m)}|\boldsymbol{\alpha}_k^{(m)})\, \mathrm{d}\mathbf{p}_i^{(m)}}_{p(\mathbf{x}_i^{(m)}|\boldsymbol{\alpha}_k^{(m)})}\,. \end{aligned} \tag{S4}$$

We notice that the term $p(\mathbf{x}_i^{(m)}|\boldsymbol{\alpha}_k^{(m)})$ above involves the likelihood and prior for the $m^{\text{th}}$ chromatin feature and corresponds to the Dirichlet-multinomial compound distribution (Mosimann, 1962), which in this application has the following specific expression:

$$p(\mathbf{x}_i^{(m)}|\boldsymbol{\alpha}_k^{(m)}) = \text{Dirichlet-multinomial}(\mathbf{x}_i^{(m)}|\boldsymbol{\alpha}_k^{(m)}) \tag{S5}$$

$$= \frac{\Gamma(J_i^{(m)}+1)\Gamma(\sum_{j=1}^{L}\alpha_{kj}^{(m)})\prod_{j=1}^{L}\Gamma(x_{ij}^{(m)}+\alpha_{kj}^{(m)})}{\Gamma(\sum_{j=1}^{L}x_{ij}^{(m)}+\sum_{j=1}^{L}\alpha_{kj}^{(m)})\prod_{j=1}^{L}\Gamma(x_{ij}^{(m)}+1)\prod_{j=1}^{L}\Gamma(\alpha_{kj}^{(m)})}. \tag{S6}$$

The product Dirichlet-multinomial (compound) likelihood of a single sample $i$ (Equation 8 in Suppl. Figure S1) is thus

$$p(\mathbf{x}_i^*|\boldsymbol{\alpha}^*,\boldsymbol{\pi}) = \sum_{k=1}^{K}\pi_k\prod_{m=1}^{M}\text{Dirichlet-multinomial}(\mathbf{x}_i^{(m)}|\boldsymbol{\alpha}_k^{(m)}). \tag{S7}$$

For all $N$ samples the likelihood is

$$p(\mathbf{X}^*|\boldsymbol{\alpha}^*,\boldsymbol{\pi}) = \prod_{i=1}^{N}\sum_{k=1}^{K}\pi_k\prod_{m=1}^{M}\text{Dirichlet-multinomial}(\mathbf{x}_i^{(m)}|\boldsymbol{\alpha}_k^{(m)}). \tag{S8}$$

#### S1.5 The prior for $\boldsymbol{\alpha}^*$

The prior for $\boldsymbol{\alpha}^*$ can be written as follows

$$p(\boldsymbol{\alpha}_1^{(1)},...,\boldsymbol{\alpha}_K^{(1)},\boldsymbol{\alpha}_1^{(M)}...,\boldsymbol{\alpha}_K^{(M)}) \propto \prod_{m=1}^{M}\prod_{k=1}^{K}\Gamma(h_k^{(m)}|\eta_h,\nu_h)\prod_{j=1}^{L}\Gamma(\alpha_{kj}^{(m)}|\eta,\nu)r \tag{S9}$$

$$\propto \prod_{m=1}^{M}\prod_{k=1}^{K}\frac{\nu_h^{\eta_h}h_k^{(m)\,\eta_h-1}e^{-\nu_h h_k^{(m)}}}{\Gamma(\eta_h)}\prod_{j=1}^{L}\frac{\nu^{\eta}\alpha_{kj}^{(m)\,\eta-1}e^{-\nu\alpha_{kj}^{(m)}}}{\Gamma(\eta)}$$

$$\propto \Gamma(\eta_h)^{-MK}\nu_h^{\eta_h MK}\Gamma(\eta)^{-MKL}\nu^{\eta MKL}$$
$$\times \exp\left\{-\sum_{m=1}^{M}\sum_{k=1}^{K}\left(\nu_h h_k^{(m)}+\sum_{j=1}^{L}\nu\alpha_{kj}^{(m)}\right)\right\}$$
$$\times \prod_{m=1}^{M}\prod_{k=1}^{K}h_k^{(m)\,\eta_h-1}\prod_{j=1}^{L}\alpha_{kj}^{(m)\,\eta-1}.$$

#### S1.6 Effect of the regularisation

The joint prior density of two independent (unregularised) Dirichlet parameters $\alpha_1$ and $\alpha_2$ is plotted in Figure S2a with hyperparameter values $\eta = 3$ and $\nu = 1$, whereas the regularised prior density based on Equation 3 in Suppl. Figure S1 is plotted in Suppl. Figure S2b with hyperparameter values $\eta = 3$, $\nu = 1$, $\eta_h = 1$, and $\nu_h = 4$. The prior for regularisation parameters $h_k^{(m)}$ reduces the differences between, i.e. smoothes the consecutive Dirichlet parameters.

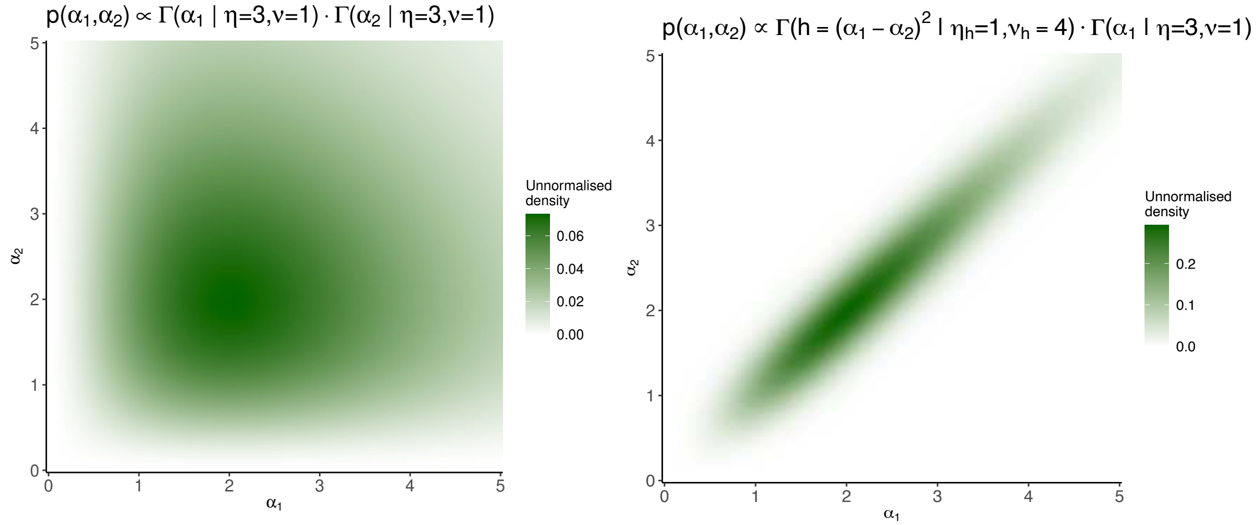

(a) Hyperprior without regularisation

(b) Hyperprior with regularisation

Figure S2: The prior densities of two Dirichlet parameters ($\alpha_1$ and $\alpha_2$) without regularisation (a) and with regularisation (b).

#### S1.7 The prior for $\boldsymbol{\lambda}^*$

The Dirichlet parameters $\alpha_{kj}^{(m)}$ are constrained to be positive by a reparameterisation $\boldsymbol{\lambda}_k^{(m)} = \log \boldsymbol{\alpha}_k^m$. The prior of the transformed variables $\boldsymbol{\lambda}^*$ in terms of the original variables $\boldsymbol{\alpha}^*$ is obtained using the multivariate change of variables method:

$$p(\boldsymbol{\lambda}^*) = p(\boldsymbol{\alpha}^*) |\det \mathbf{J}_{\boldsymbol{\lambda}^* \to \boldsymbol{\alpha}^*}|\,,$$

where $\mathbf{J}_{\boldsymbol{\lambda}^* \to \boldsymbol{\alpha}^*}$ is the Jacobian matrix of the transformation with $M \times K \times L$ rows and columns, and $|\det \mathbf{J}_{\boldsymbol{\lambda}^* \to \boldsymbol{\alpha}^*}|$ is the absolute value of the determinant of the Jacobian matrix. In the Jacobian matrix

$$\mathbf{J}_{\boldsymbol{\lambda}^* \to \boldsymbol{\alpha}^*} = \begin{bmatrix} \frac{\partial \alpha_{11}^{(1)}}{\partial \lambda_{11}^{(1)}} & \frac{\partial \alpha_{11}^{(1)}}{\partial \lambda_{12}^{(1)}} & \cdots & \frac{\partial \alpha_{11}^{(1)}}{\partial \lambda_{KL}^{(M)}} \\ \frac{\partial \alpha_{12}^{(1)}}{\partial \lambda_{11}^{(1)}} & \frac{\partial \alpha_{12}^{(1)}}{\partial \lambda_{12}^{(1)}} & \cdots & \frac{\partial \alpha_{12}^{(1)}}{\partial \lambda_{KL}^{(M)}} \\ \vdots & \vdots & \ddots & \vdots \\ \frac{\partial \alpha_{KL}^{(M)}}{\partial \lambda_{11}^{(1)}} & \frac{\partial \alpha_{KL}^{(M)}}{\partial \lambda_{12}^{(1)}} & \cdots & \frac{\partial \alpha_{KL}^{(M)}}{\partial \lambda_{KL}^{(M)}} \end{bmatrix},$$

only the diagonal terms are nonzero, i.e. $\mathbf{J}_{\boldsymbol{\lambda}^* \to \boldsymbol{\alpha}^*}$ is diagonal. The determinant of the diagonal matrix is just the product of its diagonal elements:

$$\det \mathbf{J}_{\boldsymbol{\lambda}^* \to \boldsymbol{\alpha}^*} = \prod_{m=1}^{M} \prod_{k=1}^{K} \prod_{j=1}^{L} \frac{\partial \alpha_{kj}^{(m)}}{\partial \lambda_{kj}^{(m)}} = \prod_{m=1}^{M} \prod_{k=1}^{K} \prod_{j=1}^{L} \frac{\partial e^{\lambda_{kj}^{(m)}}}{\partial \lambda_{kj}^{(m)}} = \prod_{m=1}^{M} \prod_{k=1}^{K} \prod_{j=1}^{L} e^{\lambda_{kj}^{(m)}} = \prod_{m=1}^{M} \prod_{k=1}^{K} \prod_{j=1}^{L} \alpha_{kj}^{(m)}.$$

Multiplying $p(\boldsymbol{\alpha}^*)$ by the absolute value of the determinant of the Jacobian matrix yields $p(\boldsymbol{\lambda}^*)$:

$$p(\boldsymbol{\lambda}_1^{(1)},...,\boldsymbol{\lambda}_K^{(1)},\boldsymbol{\lambda}_1^{(M)}...,\boldsymbol{\lambda}_K^{(M)}) \propto \prod_{m=1}^{M}\prod_{k=1}^{K}\Gamma(h_k^{(m)};\eta_h,\nu_h)\prod_{j=1}^{L}\Gamma(\alpha_{kj}^{(m)};\eta,\nu)\,|\det \mathbf{J}_{\lambda\to\alpha}| \tag{S10}$$

$$\propto \prod_{m=1}^{M}\prod_{k=1}^{K}\Gamma(h_k^{(m)};\eta_h,\nu_h)\prod_{j=1}^{L}\Gamma(\alpha_{kj}^{(m)};\eta,\nu)\left|\prod_{m=1}^{M}\prod_{k=1}^{K}\prod_{j=1}^{L}\alpha_{kj}^{(m)}\right|$$

$$\propto \Gamma(\eta_h)^{-MK}\nu_h^{\eta_h MK}\Gamma(\eta)^{-MKL}\nu^{\eta MKL} \times \exp\left\{-\sum_{m=1}^{M}\sum_{k=1}^{K}\left(\nu_h h_k^{(m)} + \sum_{j=1}^{L}\nu\alpha_{kj}^{(m)}\right)\right\} \times \prod_{m=1}^{M}\prod_{k=1}^{K} h_k^{(m)\,\eta_h-1}\prod_{j=1}^{L}\alpha_{kj}^{(m)\,\eta-1}\alpha_{kj}^{(m)}$$

$$\propto \Gamma(\eta_h)^{-MK}\nu_h^{\eta_h MK}\Gamma(\eta)^{-MKL}\nu^{\eta MKL} \times \exp\left\{-\sum_{m=1}^{M}\sum_{k=1}^{K}\left(\nu_h h_k^{(m)} + \sum_{j=1}^{L}\nu\alpha_{kj}^{(m)}\right)\right\} \times \prod_{m=1}^{M}\prod_{k=1}^{K} h_k^{(m)\,\eta_h-1}\prod_{j=1}^{L}\alpha_{kj}^{(m)\,\eta}.$$

#### S1.8 The expected log unnormalised posterior (lower bound) $Q(\boldsymbol{\theta}, \boldsymbol{\theta}^{\text{old}})$

Here we derive an expression for the lower bound of the expected log (unnormalised posterior) $Q(\boldsymbol{\theta}, \boldsymbol{\theta}^{\text{old}})$ used in the EM algorithm. The lower bound can be derived as follows

$$
\begin{aligned}
Q(\boldsymbol{\theta}, \boldsymbol{\theta}^{\text{old}}) &= \mathbb{E}_{p(\mathbf{Z}|\mathbf{X}^*, \boldsymbol{\theta}^{\text{old}})} \left[ \log p(\mathbf{X}^*, \mathbf{Z}|\boldsymbol{\theta}) + \log p(\boldsymbol{\theta}) \right] \qquad \text{(S11)} \\
&= \mathbb{E}_{p(\mathbf{Z}|\mathbf{X}^*, \boldsymbol{\theta}^{\text{old}})} \left[ \log p(\mathbf{Z}|\boldsymbol{\theta}) + \log p(\mathbf{X}^*|\mathbf{Z}, \boldsymbol{\theta}) + \log p(\boldsymbol{\theta}) \right] \\
&= \mathbb{E}_{p(\mathbf{Z}|\mathbf{X}^*, \boldsymbol{\theta}^{\text{old}})} \left[ \left[ \log \prod_{i=1}^{N} p(\mathbf{z}_i|\boldsymbol{\pi}) \right] + \left[ \log \prod_{i=1}^{N} \prod_{m=1}^{M} p(\mathbf{x}_i^{(m)}|\mathbf{z}_i, \boldsymbol{\theta}) \right] + \log p(\boldsymbol{\alpha}^*) + \log p(\boldsymbol{\pi}) \right] \\
&= \mathbb{E}_{p(\mathbf{Z}|\mathbf{X}^*, \boldsymbol{\theta}^{\text{old}})} \left[ \left[ \log \prod_{i=1}^{N} \prod_{k=1}^{K} \pi_k^{z_{ik}} \right] + \left[ \log \prod_{i=1}^{N} \prod_{m=1}^{M} \prod_{k=1}^{K} \left[ p(\mathbf{x}_i^{(m)}|\boldsymbol{\theta}) \right]^{z_{ik}} \right] + \log p(\boldsymbol{\alpha}^*) + \log p(\boldsymbol{\pi}) \right] \\
&= \mathbb{E}_{p(\mathbf{Z}|\mathbf{X}^*, \boldsymbol{\theta}^{\text{old}})} \left[ \left[ \sum_{i=1}^{N} \sum_{k=1}^{K} z_{ik} \log \pi_k + \sum_{i=1}^{N} \sum_{k=1}^{K} z_{ik} \sum_{m=1}^{M} \log p(\mathbf{x}_i^{(m)}|\boldsymbol{\theta}) \right] + \log p(\boldsymbol{\alpha}^*) + \log p(\boldsymbol{\pi}) \right] \\
&= \sum_{i=1}^{N} \sum_{k=1}^{K} \mathbb{E}[z_{ik}] \log \pi_k + \sum_{i=1}^{N} \sum_{k=1}^{K} \mathbb{E}[z_{ik}] \sum_{m=1}^{M} \log p(\mathbf{x}_i^{(m)}|\boldsymbol{\theta}) + \log p(\boldsymbol{\alpha}^*) + \log p(\boldsymbol{\pi}),
\end{aligned}
$$

where $\mathbb{E}[z_{ik}] = \mathbb{E}_{p(\mathbf{Z}|\mathbf{X}^*, \boldsymbol{\theta}^{\text{old}})}[z_{ik}] = p(z_{ik} = 1|\mathbf{x}_i, \boldsymbol{\theta}^{\text{old}})$. The likelihood term $p(\mathbf{x}_i^{(m)}|\boldsymbol{\theta})$ in Equation S11 is the Dirichlet-multinomial compound distribution derived in Suppl. Section S1.4 (Equation S6).

#### S1.9 The lower bound $Q(\boldsymbol{\theta}, \boldsymbol{\theta}^{\textbf{old}})$ – terms depending on $\boldsymbol{\alpha}^*$

Let us divide the lower bound into two parts

$$
Q(\boldsymbol{\theta}, \boldsymbol{\theta}^{\text{old}}) = \mathbb{E}_{p(\mathbf{Z}|\mathbf{X}^*, \boldsymbol{\theta}^{\text{old}})} \left[ \log p(\mathbf{X}^*, \mathbf{Z}|\boldsymbol{\theta}) \right] + \mathbb{E}_{p(\mathbf{Z}|\mathbf{X}^*, \boldsymbol{\theta}^{\text{old}})} \left[ \log p(\boldsymbol{\theta}) \right]
$$

and concentrate first on the expected complete data likelihood:

$$
\mathbb{E}_{p(\mathbf{Z}|\mathbf{X}^*, \boldsymbol{\theta}^{\text{old}})} \left[ \log p(\mathbf{X}^*, \mathbf{Z}|\boldsymbol{\theta}) \right] = \sum_{i=1}^{N} \sum_{k=1}^{K} \mathbb{E}[z_{ik}] \log \pi_k + \sum_{i=1}^{N} \sum_{k=1}^{K} \mathbb{E}[z_{ik}] \sum_{m=1}^{M} \log p(\mathbf{x}_i^{(m)}|\boldsymbol{\theta}). \qquad \text{(S12)}
$$

Substituting the Dirichlet-multinomial compound distribution (Equation S6) in place of $p(\mathbf{x}_i^{(m)}|\boldsymbol{\theta})$ in Equation S12 and considering only terms depending on $\boldsymbol{\alpha}^*$, we get

$$
\begin{aligned}
&\sum_{i=1}^{N} \sum_{k=1}^{K} \mathbb{E}[z_{ik}] \sum_{m=1}^{M} \left\{ \sum_{j=1}^{L} \log \Gamma \left( x_{ij}^{(m)} + \alpha_{kj}^{(m)} \right) - \log \Gamma \left( \sum_{j=1}^{L} x_{ij}^{(m)} + \sum_{j=1}^{L} \alpha_{kj}^{(m)} \right) \right. \\
&\qquad \left. - \left[ \sum_{j=1}^{L} \log \Gamma \left( \alpha_{kj}^{(m)} \right) - \log \Gamma \left( \sum_{j=1}^{L} \alpha_{kj}^{(m)} \right) \right] \right\} \\
&= \sum_{m=1}^{M} \sum_{k=1}^{K} \sum_{i=1}^{N} \mathbb{E}[z_{ik}] \left\{ \sum_{j=1}^{L} \log \Gamma \left( x_{ij}^{(m)} + \alpha_{kj}^{(m)} \right) - \log \Gamma \left( \sum_{j=1}^{L} x_{ij}^{(m)} + \sum_{j=1}^{L} \alpha_{kj}^{(m)} \right) \right. \\
&\qquad \left. - \left[ \sum_{j=1}^{L} \log \Gamma \left( \alpha_{kj}^{(m)} \right) - \log \Gamma \left( \sum_{j=1}^{L} \alpha_{kj}^{(m)} \right) \right] \right\}. \qquad \text{(S13)}
\end{aligned}
$$

The second term in $Q(\boldsymbol{\theta}, \boldsymbol{\theta}^{\text{old}})$, the expected logarithm of the prior, does not depend on $z_{ik}$:

$$\mathbb{E}_{p(\mathbf{Z}|\mathbf{X}^*, \boldsymbol{\theta}^{\text{old}})}\left[\log p(\boldsymbol{\theta})\right] = \log p(\boldsymbol{\alpha}_1^{(1)}, ..., \boldsymbol{\alpha}_K^{(1)}, \boldsymbol{\alpha}_1^{(M)} ..., \boldsymbol{\alpha}_K^{(M)}) + \log p(\boldsymbol{\pi})\,.$$

Using the definition of the prior in Equation S9 and considering again only the terms depending on $\boldsymbol{\alpha}^*$, we get

$$\begin{aligned}\log p(\boldsymbol{\alpha}^*) = &-\sum_{m=1}^{M}\sum_{k=1}^{K}\left[\nu_h h_k^{(m)} + \sum_{j=1}^{L}\nu\alpha_{kj}^{(m)}\right] \\ &+\sum_{m=1}^{M}\sum_{k=1}^{K}(\eta_h - 1)\log h_k^{(m)} + \sum_{m=1}^{M}\sum_{k=1}^{K}\sum_{j=1}^{L}(\eta - 1)\log\alpha_{kj}^{(m)} \\ =&\sum_{m=1}^{M}\sum_{k=1}^{K}\left\{(\eta_h - 1)\log h_k^{(m)} - \nu_h h_k^{(m)} + \sum_{j=1}^{L}\left[(\eta - 1)\log\alpha_{kj}^{(m)} - \nu\alpha_{kj}^{(m)}\right]\right\}\,.\end{aligned} \tag{S14}$$

The whole lower bound considering only the terms depending on $\boldsymbol{\alpha}^*$ is thus

$$\begin{aligned}Q(\boldsymbol{\theta}, \boldsymbol{\theta}^{\text{old}}) = \sum_{m=1}^{M}\sum_{k=1}^{K}\Bigg\{&(\eta_h - 1)\log h_k^{(m)} - \nu_h h_k^{(m)} + \sum_{j=1}^{L}\left[(\eta - 1)\log\alpha_{kj}^{(m)} - \nu\alpha_{kj}^{(m)}\right] \\ &+\sum_{i=1}^{N}\mathbb{E}[z_{ik}]\Bigg[\sum_{j=1}^{L}\log\Gamma\left(x_{ij}^{(m)} + \alpha_{kj}^{(m)}\right) - \log\Gamma\left(\sum_{j=1}^{L}x_{ij}^{(m)} + \sum_{j=1}^{L}\alpha_{kj}^{(m)}\right) \\ &-\left(\sum_{j=1}^{L}\log\Gamma\left(\alpha_{kj}^{(m)}\right) - \log\Gamma\left(\sum_{j=1}^{L}\alpha_{kj}^{(m)}\right)\right)\Bigg]\Bigg\}\,.\end{aligned} \tag{S15}$$

#### S1.10 The lower bound $Q(\boldsymbol{\theta}, \boldsymbol{\theta}^{\text{old}})$ – terms depending on $\boldsymbol{\alpha}_k^{(m)}$

To update the estimate for $\boldsymbol{\alpha}^*$ in the M-step of the Expectation-Maximisation algorithm, the lower bound is maximised wrt. $\boldsymbol{\alpha}_k^{(m)}$ for each $k$ and $m$ separately employing the BFGS algorithm. The terms in Equation S15 that depend only on $\boldsymbol{\alpha}_k^{(m)}$ are

$$\begin{aligned}&(\eta_h - 1)\log h_k^{(m)} - \nu_h h_k^{(m)} + (\eta - 1)\sum_{j=1}^{L}\log\alpha_{kj}^{(m)} - \nu\sum_{j=1}^{L}\alpha_{kj}^{(m)} \\ &\sum_{i=1}^{N}\sum_{j=1}^{L}\mathbb{E}[z_{ik}]\log\Gamma\left(x_{ij}^{(m)} + \alpha_{kj}^{(m)}\right) - \sum_{i=1}^{N}\mathbb{E}[z_{ik}]\log\Gamma\left(\sum_{j=1}^{L}x_{ij}^{(m)} + \sum_{j=1}^{L}\alpha_{kj}^{(m)}\right) \\ &\qquad - \sum_{i=1}^{N}\mathbb{E}[z_{ik}]\left[\sum_{j=1}^{L}\log\Gamma\left(\alpha_{kj}^{(m)}\right) + \log\Gamma\left(\sum_{j=1}^{L}\alpha_{kj}^{(m)}\right)\right].\end{aligned} \tag{S16}$$

#### S1.11 The derivative of the lower bound wrt. $\alpha_{kj}^{(m)}$

The BFGS algorithm requires the derivative of the function to be optimised. The derivative of lower bound w.r.t. $\alpha_{kj}^{(m)}$ is derived considering the terms of the lower bound which depend only on $\alpha_{kj}^{(m)}$

$$
\begin{aligned}
&(\eta_h - 1)\log h_k^{(m)} - \nu_h h_k^{(m)} + (\eta - 1)\log \alpha_{kj}^{(m)} - \nu \alpha_{kj}^{(m)} \\
&+ \sum_{i=1}^{N} \mathbb{E}[z_{ik}] \Bigg[ \log\Gamma\left(x_{ij}^{(m)} + \alpha_{kj}^{(m)}\right) - \log\Gamma\left(\sum_{j=1}^{L} x_{ij}^{(m)} + \sum_{j=1}^{L} \alpha_{kj}^{(m)}\right) \\
&\qquad - \left( \log\Gamma\left(\alpha_{kj}^{(m)}\right) - \log\Gamma\left(\sum_{j=1}^{L} \alpha_{kj}^{(m)}\right)\right)\Bigg].
\end{aligned}
\tag{S17}
$$

The derivative can be written using digamma function $\psi(x) = \frac{d(\log\Gamma(x))}{dx} = \frac{\Gamma'(x)}{\Gamma(x)}$ as follows:

$$
\begin{aligned}
\frac{\partial Q(\boldsymbol{\theta}, \boldsymbol{\theta}^{\text{old}})}{\partial \alpha_{kj}^{(m)}} =& (\eta_h - 1)\frac{\frac{\partial h_k^{(m)}}{\partial \alpha_{kj}^{(m)}}}{h_k^{(m)}} - \nu_h \frac{\partial h_k^{(m)}}{\partial \alpha_{kj}^{(m)}} + \frac{(\eta - 1)}{\alpha_{kj}^{(m)}} - \nu \\
&+ \sum_{i=1}^{N} \mathbb{E}[z_{ik}] \Bigg[ \psi\left(x_{ij}^{(m)} + \alpha_{kj}^{(m)}\right) - \psi\left(\sum_{j=1}^{L} x_{ij}^{(m)} + \sum_{j=1}^{L} \alpha_{kj}^{(m)}\right) \\
&\qquad - \left( \psi(\alpha_{kj}^{(m)}) - \psi(\sum_{j=1}^{L} \alpha_{kj}^{(m)})\right)\Bigg].
\end{aligned}
\tag{S18}
$$

The term $\frac{\partial h_k^{(m)}}{\partial \alpha_{kj}^{(m)}}$ in Equation S18 can be written as

$$
g_{jk}^{(m)} = \frac{\partial h_k^{(m)}}{\partial \alpha_{kj}^{(m)}} = \frac{\partial \sum_{j=2}^{L} (\alpha_{kj}^{(m)} - \alpha_{k,j-1}^{(m)})^2}{\partial \alpha_{kj}^{(m)}} = \begin{cases} 2(\alpha_{kj}^{(m)} - \alpha_{k,j+1}^{(m)}) & \text{for } j = 1 \\ 2(\alpha_{kj}^{(m)} - \alpha_{k,j-1}^{(m)}) & \text{for } j = L \\ 2(2\alpha_{kj}^{(m)} - \alpha_{k,j+1}^{(m)} - \alpha_{k,j-1}^{(m)}) & \text{otherwise.} \end{cases}
\tag{S19}
$$

#### S1.12 The derivative of the lower bound wrt. $\lambda_{kj}^{(m)}$

The derivative of the lower bound in Equation S11 wrt. $\lambda_{kj}^{(m)}$ according to the chain rule is

$$
\begin{aligned}
\frac{\partial Q(\boldsymbol{\theta}, \boldsymbol{\theta}^{\text{old}})}{\partial \lambda_{kj}^{(m)}} &= \frac{\partial Q(\boldsymbol{\theta}, \boldsymbol{\theta}^{\text{old}})}{\partial \alpha_{kj}^{(m)}} \frac{d\alpha_{kj}^{(m)}}{d\lambda_{kj}^{(m)}} \\
&= \frac{\partial Q(\boldsymbol{\theta}, \boldsymbol{\theta}^{\text{old}})}{\partial \alpha_{kj}^{(m)}} \frac{1}{\frac{d\lambda_{kj}^{(m)}}{d\alpha_{kj}^{(m)}}} \\
&= \frac{\partial Q(\boldsymbol{\theta}, \boldsymbol{\theta}^{\text{old}})}{\partial \alpha_{kj}^{(m)}} \frac{1}{\frac{1}{\alpha_{kj}^{(m)}}} \\
&= \frac{\partial Q(\boldsymbol{\theta}, \boldsymbol{\theta}^{\text{old}})}{\partial \alpha_{kj}^{(m)}} \alpha_{kj}^{(m)} \\
&= \alpha_{kj}^{(m)} \left( (\eta_h - 1) \frac{g_{jk}^{(m)}}{h_k^{(m)}} - \nu_h g_{jk}^{(m)} + \frac{\eta - 1}{\alpha_{kj}^{(m)}} - \nu \right) \\
&\quad + \alpha_{kj}^{(m)} \sum_{i=1}^{N} \mathbb{E}[z_{ik}] \Bigg[ \psi \left( x_{ij}^{(m)} + \alpha_{kj}^{(m)} \right) - \psi \left( \sum_{j=1}^{L} x_{ij}^{(m)} + \sum_{j=1}^{L} \alpha_{kj}^{(m)} \right) \\
&\quad - \left( \psi \left( \alpha_{kj}^{(m)} \right) - \psi \left( \sum_{j=1}^{L} \alpha_{kj}^{(m)} \right) \right) \Bigg].
\end{aligned}
\tag{S20}
$$

The terms computed according to Equation S20 for each $j$ are the elements of the gradient required by BFGS.

#### S1.13 The EM updates for cluster assignments $\mathbb{E}[z_{ik}]$

The cluster assignments $\mathbb{E}[z_{ik}] = p(z_{ik} = 1 | \mathbf{x}_i^*, \boldsymbol{\theta}^{\text{old}})$ are computed as

$$
\begin{aligned}
p(z_{ik} = 1 | \mathbf{x}_i^*, \boldsymbol{\theta}^{\text{old}}) &= \frac{p(z_{ik} = 1 | \boldsymbol{\theta}^{\text{old}}) \prod_{m=1}^{M} p(\mathbf{x}_i^{(m)} | z_{ik} = 1, \boldsymbol{\theta}^{\text{old}})}{\sum_{k'=1}^{K} p(z_{ik'} = 1 | \boldsymbol{\theta}^{\text{old}}) \prod_{m=1}^{M} p(\mathbf{x}_i^{(m)} | z_{ik'} = 1, \boldsymbol{\theta}^{\text{old}})} \\
&= \frac{\pi_k^{\text{old}} \prod_{m=1}^{M} \text{Dirichlet-multinomial}(\mathbf{x}_i^{(m)} | \boldsymbol{\alpha}_k^{(m,\,\text{old})})}{\sum_{k'=1}^{K} \pi_{k'}^{\text{old}} \prod_{m=1}^{M} \text{Dirichlet-multinomial}(\mathbf{x}_i^{(m)} | \boldsymbol{\alpha}_{k'}^{(m,\,\text{old})})}.
\end{aligned}
\tag{S21}
$$

#### S1.14 The EM updates for mixture weights $\boldsymbol{\pi}$

The mixing weights are updated as

$$\pi_k = \frac{\sum_{i=1}^{N} \mathbb{E}[z_{ik}]}{\sum_{i=1}^{N} \sum_{k'=1}^{K} \mathbb{E}[z_{ik'}]} \,.$$

#### S1.15 Checking the EM convergence

Computing the negative log unnormalised posterior given the current parameter estimates is utilised to check the convergence of the EM algorithm. EM algorithm is guaranteed to maximise the lower bound (the expected log unnormalised posterior) and, therefore to maximise the log unnormalised posterior. The negative log unnormalised posterior is

$$\begin{aligned}
-\log p(\boldsymbol{\alpha}^*, \boldsymbol{\pi}) &\propto -\log p(\mathbf{X}|\boldsymbol{\alpha}^*, \boldsymbol{\pi}) - \log p(\boldsymbol{\alpha}^*) \\
&= -\sum_{i=1}^{N} \log \left\{ \sum_{k=1}^{K} \pi_k \exp \left[ \sum_{m=1}^{M} \log \text{Dirichlet-multinomial}(\mathbf{x}_i^{(m)}|\boldsymbol{\alpha}_k^{(m)}) \right] \right\} \\
&\quad - \sum_{m=1}^{M} \sum_{k=1}^{K} \log \Gamma(h_k^{(m)}|\eta_h, \nu_h) \\
&\quad - \sum_{m=1}^{M} \sum_{k=1}^{K} \sum_{j=1}^{L} \log \Gamma(\alpha_{kj}^{(m)}|\eta, \nu) \\
&= -\sum_{i=1}^{N} \log \Bigg\{ \sum_{k=1}^{K} \pi_k \exp \Bigg[ \sum_{m=1}^{M} \Bigg( \log \Gamma(J_i^{(m)} + 1) + \log \Gamma(\sum_{j=1}^{L} \alpha_{kj}^{(m)}) + \sum_{j=1}^{L} \log \Gamma(x_{ij}^{(m)} + \alpha_{kj}^{(m)}) \\
&\qquad - \log \Gamma(\sum_{j=1}^{L} x_{ij}^{(m)} + \sum_{j=1}^{L} \alpha_{kj}^{(m)}) - \sum_{j=1}^{L} \log \Gamma(x_{ij}^{(m)} + 1) - \sum_{j=1}^{L} \log \Gamma(\alpha_{kj}^{(m)}) \Bigg) \Bigg] \Bigg\} \\
&\quad - MK\eta_h \log \nu_h - (\eta_h - 1) \sum_{m=1}^{M} \sum_{k=1}^{K} \log h_k^{(m)} + \nu_h \sum_{m=1}^{M} \sum_{k=1}^{K} h_k^{(m)} + MK \log \Gamma(\eta_h) \\
&\quad - MKL\eta \log \nu - (\eta - 1) \sum_{m=1}^{M} \sum_{k=1}^{K} \sum_{j=1}^{L} \log \alpha_{kj}^{(m)} + \nu \sum_{m=1}^{M} \sum_{k=1}^{K} \sum_{j=1}^{L} \alpha_{kj}^{(m)} + MKL \log \Gamma(\eta).
\end{aligned}$$

### S2 Product Dirichlet-multinomial mixture model with shifting and flipping

#### S2.1 Components and a directed acyclic graph of the model

This section describes the Dirichlet-multinomial mixture model that enables both the shifting and flipping of the chromatin feature profiles. The proposed model is presented as a directed acyclic graph in Suppl. Figure S3 together with the distributions of individual components. Note that in Suppl. Figure S3 and in the following sections, we denote the length of chromatin feature profile $\mathbf{x}^{(m)}$ as $L_{\mathbf{x}}$ to separate it from the length $L_{\boldsymbol{\alpha}}$ of the Dirichlet parameters $\boldsymbol{\alpha}_k^{(m)}$.

We introduce shift and flip states for each genomic loci. For the number of possible shift states, we set an odd number $S$; the middle shift state $s = \frac{S+1}{2}$ corresponds to no-shift-state. The maximum allowed shift either upstream or downstream is $\frac{S-1}{2} \times B$ nucleotides, where $B$ is the data resolution. To implement the shifting procedure in ChromDMM, we extended the length of the Dirichlet parameter $\boldsymbol{\alpha}_k^m$ from $L_{\mathbf{x}}$ to $L_{\boldsymbol{\alpha}} = L_{\mathbf{x}} + S - 1$. The number of possible flip states is 2; either the original profile is considered (flip state 1) or the profile is reversed (flip state 2).

When evaluating the likelihood for a shift state $s$ and flip state $f$, we use the corresponding $L$-length subset of the extended Dirichlet parameters for each mixture component $k$, denoted as $\boldsymbol{\alpha}^*_{ksf}$ either in it's original form ($f = 1$, Equation 8 in Suppl. Figure S3) or in its reversed form ($f = 2$, Equation 9 in Suppl. Figure S3). In addition, for each locus, we define prior probabilities for the shift and flip states. The prior shift state probabilities for the genomic locus $i$ are denoted as $\boldsymbol{\xi}_i = (\xi_{i1}, \ldots, \xi_{iS})$, where $\sum_{s=1}^{S} \xi_{is} = 1$. If the anchor points of the genomic loci being clustered are defined according to ChIP-seq peak summits, then the shift state prior can be defined, e.g. as a pyramid-shaped prior that has the highest probability at the ChIP-seq peak summit (corresponding to no-shift-state $s = \frac{S+1}{2}$) and linearly decreasing the prior to zero beyond the maximum shift states. The prior flip state probabilities for the genomic locus $i$ are denoted as $\boldsymbol{\zeta}_i = (\zeta_{i1}, \zeta_{i2})$, where $\sum_{f=1}^{2} \zeta_{if} = 1$.

The latent cluster membership variables are re-defined as follows: $z_{iksf} = 1$ if the sample $i$ originates from the cluster $k$, has shift state $s$, and has strand-orientation $f$; otherwise $z_{iksf} = 0$. These latent variables are stored in $N \times K \times S \times 2$ matrix (or tensor) $\mathbf{Z}$.

The Dirichlet-multinomial mixture model can be applied *(i)* only with shifting, *(ii)* only with flipping, or *(iii)* with both shifting and flipping. In case of *(i)*, we simply drop the index $f$, the corresponding sums and products, and the flip state prior terms $\zeta_{if}$ in Equations presented in Suppl. Figure S3. In addition, we only need to consider the definition of $\alpha_{kjs}^{(m)} = \alpha_{k,j+s-1}^{(m)}$( Equation 8 in Suppl. Figure S3). In case of *(ii)*, we simply drop the index $s$, the corresponding sums and products, and the shift state prior terms $\xi_{is}$ in Equations presented in Suppl.

Figure S3. In the flipping-only model, both the chromatin feature profiles and the Dirichlet parameters are of the same length $L_{\mathbf{x}}$. In addition, $\alpha_{kj,f=1}^{(m)} = \alpha_{kj}^{(m)}$, and $\alpha_{kj,f=2}^{(m)} = \alpha_{k,L_{\mathbf{x}}-j+1}^{(m)}$. In both the shifting-only model *(i)* and the flipping-only model *(ii)*, the dimensionality of the latent variables $\mathbf{z}_i$ are 2 ($K \times S$ and $K \times 2$, respectively). The equations needed for the implementation of these models are derived similarly as presented in this Section for the full model *(iii)*.

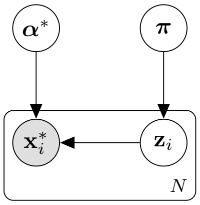

$N$: number of samples (genomic loci)

$M$: number of chromatin features

$K$: number of clusters

$S$: number of possible shift states (odd)

$L_{\boldsymbol{\alpha}}$: length of $\boldsymbol{\alpha}_k^m$

$L_{\mathbf{x}}$: length of $\boldsymbol{x}_i^m$

$L_{\boldsymbol{\alpha}} = L_{\mathbf{x}} + S - 1$

$\mathbf{z}_{ik}$: the shift and flip state state of $\boldsymbol{\alpha}_k^*$ corresponding to sample $\mathbf{x}_i^*$ (1)

binary indicator variable $z_{iksf} \in \{0, 1\}$,

If for sample $i$ the shift state is $s$, flip state is $f \in \{1, 2\}$ and cluster assignment is $k$,

$z_{iksf} = 1$, and other elements of matrix $\mathbf{z}_{ik}$ are zero

$$\boldsymbol{\alpha}^* = \left[\boldsymbol{\alpha}_1^{(1)}, \ldots, \boldsymbol{\alpha}_K^{(1)}, \ldots, \boldsymbol{\alpha}_1^{(M)}, \ldots, \boldsymbol{\alpha}_K^{(M)}\right] \tag{2}$$

$$p(\boldsymbol{\alpha}^*) \propto \prod_{m=1}^{M} \prod_{k=1}^{K} \Gamma(h_k^{(m)} | \eta_h, \nu_h) \prod_{j=1}^{L_{\boldsymbol{\alpha}}} \Gamma(\alpha_{kj}^{(m)} | \eta, \nu) \tag{3}$$

$$h_k^{(m)} = \sum_{j=2}^{L_{\boldsymbol{\alpha}}} (\alpha_{kj}^{(m)} - \alpha_{k,\, j-1}^{(m)})^2 \tag{4}$$

$$p(\pi_k) \propto 1\,, \sum_{k=1}^{K} \pi_k = 1 \tag{5}$$

$\boldsymbol{\xi}_i$: Prior shift state probabilities for sample $i$ with elements $\xi_{is}$, $\sum_{s=1}^{S} \xi_s = 1$.

For example, highest probability with shift state $s = (S+1)/2$ (no shift)

and the probabilities decreases the further we move from "no shift" state.

To have global prior, set $\xi_{is}$ the same for all $i$.

$\boldsymbol{\zeta}_i$: Prior flip state probabilities for sample $i$ with elements $\zeta_{if}$.

To have uniform global prior, set $\zeta_{if} = 0.5$ for all $i$ and $f$.

$$p(\mathbf{z}_i | \boldsymbol{\pi}, \boldsymbol{\xi}_i, \boldsymbol{\zeta}_i) = \prod_{k=1}^{K} \prod_{s=1}^{S} \prod_{f=1}^{2} (\pi_k \xi_{is} \zeta_{if})^{z_{iksf}} \tag{6}$$

$$\alpha_{kj}^{(m)}(z_{iksf} = 1) = \alpha_{kjsf}^{(m)} \tag{7}$$

$$f = 1 : \alpha_{kj}^{(m)}(z_{iks1} = 1) = \alpha_{kjs1}^{(m)} = \alpha_{k,\, j+s-1}^{(m)} \tag{8}$$

$$f = 2 : \alpha_{kj}^{(m)}(z_{iks2} = 1) = \alpha_{kjs2}^{(m)} = \alpha_{k,\, L_{\boldsymbol{\alpha}}-S+s-j-1}^{(m)} \tag{9}$$

$$p(\mathbf{x}_i^* | \mathbf{z}_i, \boldsymbol{\alpha}^*) = \prod_{k=1}^{K} \prod_{s=1}^{S} \prod_{f=1}^{2} \prod_{m=1}^{M} \left[\text{Dirichlet-Multinomial}\left(\mathbf{x}_i^{(m)} | \boldsymbol{\alpha}_k^{(m)}(z_{iksf})\right)\right]^{z_{iksf}} \tag{10}$$

$$p(\mathbf{x}_i^*, \mathbf{z}_i | \boldsymbol{\alpha}^*, \boldsymbol{\pi}) = \prod_{k=1}^{K} \prod_{s=1}^{S} \prod_{f=1}^{2} \prod_{m=1}^{M} \left[\pi_k \xi_{is} \zeta_{if} \text{Dirichlet-Multinomial}\left(\mathbf{x}_i^{(m)} | \boldsymbol{\alpha}_k^{(m)}(z_{iksf})\right)\right]^{z_{iksf}} \tag{11}$$

$$p(\mathbf{x}_i^* | \boldsymbol{\alpha}^*, \boldsymbol{\pi}) = \sum_{k=1}^{K} \pi_k \sum_{s=1}^{S} \xi_{is} \sum_{f=1}^{2} \zeta_{if} \prod_{m=1}^{M} \text{Dirichlet-Multinomial}\left(\mathbf{x}_i^{(m)} | \boldsymbol{\alpha}_{ksf}^{(m)}\right) \tag{12}$$

Figure S3: Model diagram in a directed acyclic graph notation. The model includes both the shifting and the flipping features.

##### S2.2 Dirichlet-multinomial distribution

The Dirichlet-multinomial compound distribution with a shift state $s$ and a flip state $f$ is

$$
\begin{aligned}
p(\mathbf{x}_i^{(m)}|\boldsymbol{\alpha}_{ksf}^{(m)}) &= \text{Dirichlet-multinomial}(\mathbf{x}_i^{(m)}|\boldsymbol{\alpha}_{ksf}^{(m)}) \\
&= \frac{\Gamma(J_i^{(m)}+1)\Gamma(\sum_{j=1}^{L_{\mathbf{x}}}\alpha_{kjsf}^{(m)})}{\prod_{j=1}^{L_{\mathbf{x}}}\Gamma(x_{ij}^{(m)}+1)\prod_{j=1}^{L_{\mathbf{x}}}\Gamma(\alpha_{kjsf}^{(m)})} \times \frac{\prod_{j=1}^{L_{\mathbf{x}}}\Gamma(x_{ij}^{(m)}+\alpha_{kjsf}^{(m)})}{\Gamma(\sum_{j=1}^{L_{\mathbf{x}}}x_{ij}^{(m)}+\sum_{j=1}^{L_{\mathbf{x}}}\alpha_{kjsf}^{(m)})} \\
&= \frac{\Gamma(J_i^{(m)}+1)\Gamma(\sum_{j=1}^{L_{\mathbf{x}}}\alpha_{kjsf}^{(m)})\prod_{j=1}^{L_{\mathbf{x}}}\Gamma(x_{ij}^{(m)}+\alpha_{kjsf}^{(m)})}{\Gamma(\sum_{j=1}^{L_{\mathbf{x}}}x_{ij}^{(m)}+\sum_{j=1}^{L_{\mathbf{x}}}\alpha_{kjsf}^{(m)})\prod_{j=1}^{L_{\mathbf{x}}}\Gamma(x_{ij}^{(m)}+1)\prod_{j=1}^{L_{\mathbf{x}}}\Gamma(\alpha_{kjsf}^{(m)})},
\end{aligned}
\tag{S22}
$$

where $J_i^{(m)} = \sum_{j=1}^{L_{\mathbf{x}}} x_{ij}^{(m)}$.

#### S2.3 The expected log unnormalised posterior (lower bound) $Q(\boldsymbol{\theta}, \boldsymbol{\theta}^{\text{old}})$

The lower bound of the expected log (unnormalised) posterior $Q(\boldsymbol{\theta}, \boldsymbol{\theta}^{\text{old}})$ for the model that includes the shifting and flipping states can be written as

$$
\begin{aligned}
Q(\boldsymbol{\theta}, \boldsymbol{\theta}^{\text{old}}) &= \mathbb{E}_{p(\mathbf{Z}|\mathbf{X}^*, \boldsymbol{\theta}^{\text{old}})}\left[\log p(\mathbf{X}^*, \mathbf{Z}|\boldsymbol{\theta}) + \log p(\boldsymbol{\theta})\right] \qquad \text{(S23)} \\
&= \mathbb{E}_{p(\mathbf{Z}|\mathbf{X}^*, \boldsymbol{\theta}^{\text{old}})}\left[\log p(\mathbf{Z}|\boldsymbol{\theta}) + \log p(\mathbf{X}^*|\mathbf{Z}, \boldsymbol{\theta}) + \log p(\boldsymbol{\theta})\right] \\
&= \mathbb{E}_{p(\mathbf{Z}|\mathbf{X}^*, \boldsymbol{\theta}^{\text{old}})}\left[\log \prod_{i=1}^{N} p(\mathbf{z}_i|\boldsymbol{\pi}, \boldsymbol{\xi}_i, \boldsymbol{\zeta}_i) + \log \prod_{i=1}^{N} p(\mathbf{x}_i^*|\mathbf{z}_i, \boldsymbol{\theta}) + \log p(\boldsymbol{\theta})\right] \\
&= \mathbb{E}_{p(\mathbf{Z}|\mathbf{X}^*, \boldsymbol{\theta}^{\text{old}})}\Bigg[\log \prod_{i=1}^{N}\prod_{k=1}^{K}\prod_{s=1}^{S}\prod_{f=1}^{2} (\pi_k \xi_{is} \zeta_{if})^{z_{iksf}} + \log \prod_{i=1}^{N}\prod_{k=1}^{K}\prod_{s=1}^{S}\prod_{f=1}^{2}\left[\prod_{m=1}^{M} p(\mathbf{x}_i^{(m)}|\boldsymbol{\alpha}_{ksf})\right]^{z_{iksf}} \\
&\qquad + \log p(\boldsymbol{\theta})\Bigg] \\
&= \mathbb{E}_{p(\mathbf{Z}|\mathbf{X}^*, \boldsymbol{\theta}^{\text{old}})}\Bigg[\sum_{i=1}^{N}\sum_{k=1}^{K}\sum_{s=1}^{S}\sum_{f=1}^{2} z_{iksf}\left[\log \pi_k + \log \xi_{is} + \log \zeta_{if}\right] \\
&\qquad + \sum_{i=1}^{N}\sum_{k=1}^{K}\sum_{s=1}^{S}\sum_{f=1}^{2} z_{iksf} \sum_{m=1}^{M} \log p(\mathbf{x}_i^{(m)}|\boldsymbol{\alpha}_{ksf}^{(m)}) + \log p(\boldsymbol{\theta})\Bigg] \\
&= \sum_{i=1}^{N}\sum_{k=1}^{K}\sum_{s=1}^{S}\sum_{f=1}^{2} \mathbb{E}[z_{iksf}] \log \pi_k + \sum_{i=1}^{N}\sum_{k=1}^{K}\sum_{s=1}^{S}\sum_{f=1}^{2} \mathbb{E}[z_{iksf}] \log \xi_{is} \\
&\quad + \sum_{i=1}^{N}\sum_{k=1}^{K}\sum_{s=1}^{S}\sum_{f=1}^{2} \mathbb{E}[z_{iksf}] \log \zeta_{if} + \sum_{i=1}^{N}\sum_{k=1}^{K}\sum_{s=1}^{S}\sum_{f=1}^{2} \mathbb{E}[z_{iksf}] \sum_{m=1}^{M} \log p(\mathbf{x}_i^{(m)}|\boldsymbol{\alpha}_{ksf}^{(m)}) + \log p(\boldsymbol{\theta}) \\
&= \sum_{k=1}^{K} \log \pi_k \sum_{i=1}^{N}\sum_{s=1}^{S}\sum_{f=1}^{2} \mathbb{E}[z_{iksf}] + \sum_{i=1}^{N}\sum_{s=1}^{S} \log \xi_{is} \sum_{k=1}^{K}\sum_{f=1}^{2} \mathbb{E}[z_{iksf}] \\
&\quad + \sum_{i=1}^{N}\sum_{f=1}^{2} \log \zeta_{if} \sum_{k=1}^{K}\sum_{s=1}^{S} \mathbb{E}[z_{iksf}] + \sum_{i=1}^{N}\sum_{k=1}^{K}\sum_{s=1}^{S}\sum_{f=1}^{2} \mathbb{E}[z_{iksf}] \sum_{m=1}^{M} \log p(\mathbf{x}_i^{(m)}|\boldsymbol{\alpha}_{ksf}^{(m)}) + \log p(\boldsymbol{\theta}),
\end{aligned}
$$

where $\mathbb{E}[z_{iksf}] = p(z_{iksf} = 1|\mathbf{x}_i^*, \boldsymbol{\theta}^{\text{old}})$.

#### S2.4 Lower bound $Q(\boldsymbol{\theta}, \boldsymbol{\theta}^{\text{old}})$ – terms depending on $\boldsymbol{\alpha}_k^{(m)}$

The terms of the lower bound depending only on $\boldsymbol{\alpha}_k^{(m)}$ are presented in Equation S24. Note the two indexing $j' = 1, \dots, L_{\mathbf{x}}$ and $j = 1, \dots, L_{\boldsymbol{\alpha}}$.

$$
\begin{aligned}
&\sum_{i=1}^{N}\sum_{s=1}^{S}\mathbb{E}[z_{iks,\,f=1}]\left[\sum_{j'=1}^{L_{\mathbf{x}}}\log\Gamma\left(x_{ij'}^{(m)}+\alpha_{k,\,j'+s-1}^{(m)}\right)-\log\Gamma\left(\sum_{j'=1}^{L_{\mathbf{x}}}x_{ij'}^{(m)}+\sum_{j'=1}^{L_{\mathbf{x}}}\alpha_{k,\,j'+s-1}^{(m)}\right)\right. \\
&\qquad\left.-\left[\sum_{j'=1}^{L_{\mathbf{x}}}\log\Gamma\left(\alpha_{k,\,j'+s-1}^{(m)}\right)-\log\Gamma\left(\sum_{j'=1}^{L_{\mathbf{x}}}\alpha_{k,\,j'+s-1}^{(m)}\right)\right]\right] \\
&+\sum_{i=1}^{N}\sum_{s=1}^{S}\mathbb{E}[z_{iks,\,f=2}]\left[\sum_{j'=1}^{L_{\mathbf{x}}}\log\Gamma\left(x_{ij'}^{(m)}+\alpha_{k,\,L_{\boldsymbol{\alpha}}-S+s-j'+1}^{(m)}\right)-\log\Gamma\left(\sum_{j'=1}^{L_{\mathbf{x}}}x_{ij'}^{(m)}+\sum_{j'=1}^{L_{\mathbf{x}}}\alpha_{k,\,L_{\boldsymbol{\alpha}}-S+s-j'+1}^{(m)}\right)\right. \\
&\qquad\left.-\left[\sum_{j'=1}^{L_{\mathbf{x}}}\log\Gamma\left(\alpha_{k,\,L_{\boldsymbol{\alpha}}-S+s-j'+1}^{(m)}\right)-\log\Gamma\left(\sum_{j'=1}^{L_{\mathbf{x}}}\alpha_{k,\,L_{\boldsymbol{\alpha}}-S+s-j'+1}^{(m)}\right)\right]\right] \\
&+(\eta_h-1)\log h_k^{(m)}-\nu_h h_k^{(m)}+\sum_{j=1}^{L_{\boldsymbol{\alpha}}}\left[(\eta-1)\log\alpha_{kj}^{(m)}-\nu\alpha_{kj}^{(m)}\right],
\end{aligned}
\tag{S24}
$$

where $h_k^{(m)} = \sum_{j=2}^{L_{\boldsymbol{\alpha}}}(\alpha_{kj}^{(m)} - \alpha_{k,\,j-1}^{(m)})^2$ .

The sum of $\alpha$ terms does not depend on the order of the terms in the sum, so Equation S24 simplifies to

$$
\begin{aligned}
&\sum_{i=1}^{N}\sum_{s=1}^{S}\mathbb{E}[z_{iks,\,f=1}]\sum_{j'=1}^{L_{\mathbf{x}}}\log\Gamma\left(x_{ij'}^{(m)}+\alpha_{k,\,j'+s-1}^{(m)}\right) \\
&+\sum_{i=1}^{N}\sum_{s=1}^{S}\mathbb{E}[z_{iks,\,f=2}]\sum_{j'=1}^{L_{\mathbf{x}}}\log\Gamma\left(x_{ij'}^{(m)}+\alpha_{k,\,L_{\boldsymbol{\alpha}}-S+s-j'+1}^{(m)}\right) \\
&-\sum_{i=1}^{N}\sum_{s=1}^{S}\sum_{f=1}^{2}\mathbb{E}[z_{iksf}]\log\Gamma\left(\sum_{j'=1}^{L_{\mathbf{x}}}x_{ij'}^{(m)}+\sum_{j'=1}^{L_{\mathbf{x}}}\alpha_{k,\,j'+s-1}^{(m)}\right) \\
&-\Bigg[\sum_{i=1}^{N}\sum_{s=1}^{S}\sum_{f=1}^{2}\mathbb{E}[z_{iksf}]\sum_{j'=1}^{L_{\mathbf{x}}}\log\Gamma\left(\alpha_{k,\,j'+s-1}^{(m)}\right) \\
&\qquad\qquad-\sum_{i=1}^{N}\sum_{s=1}^{S}\sum_{f=1}^{2}\mathbb{E}[z_{iksf}]\log\Gamma\left(\sum_{j'=1}^{L_{\mathbf{x}}}\alpha_{k,\,j'+s-1}^{(m)}\right)\Bigg] \\
&+(\eta_h-1)\log h_k^{(m)}-\nu_h h_k^{(m)}+\sum_{j=1}^{L_{\boldsymbol{\alpha}}}\left[(\eta-1)\log\alpha_{kj}^{(m)}-\nu\alpha_{kj}^{(m)}\right], \\
&\text{where } h_k^{(m)}=\sum_{j=2}^{L_{\boldsymbol{\alpha}}}(\alpha_{kj}^{(m)}-\alpha_{k,\,j-1}^{(m)})^2\,.
\end{aligned}
\tag{S25}
$$

#### S2.5 The derivative of the lower bound wrt. $\alpha_{kj}^{(m)}$

To derive the derivative of the expected log unnormalised posterior wrt. $\alpha_{kj}^{(m)}$, consider the terms of Equation S25 depending on $\alpha_{kj}^{(m)}$. For each element $j$, we need to consider the shift states we need to sum over. These shift states depend on $j$, the length of the profiles $L_{\mathbf{x}}$, and the number of shift states. In addition, we introduce $L_{\mathbf{x}}-1$ extra "pseudo" shift states in both ends of the shift state vector resulting in total of $2\times(L_{\mathbf{x}}-1)$ "pseudo" shift states. The "pseudo" shift states simplify the equations.

Consider a small example where $L_{\mathbf{x}}=3$, $L_{\boldsymbol{\alpha}}=5$ and $S=3$, and omit the prior terms and the indexes $m$ and $k$ for clarity. In this case, the terms of Equation S25 are

$$\sum_{i=1}^{N} \mathbb{E}[z_{i1,\,f=1}] \Big[\log\Gamma\left(x_{i1}+\alpha_1\right)+\log\Gamma\left(x_{i2}+\alpha_2\right)+\log\Gamma\left(x_{i3}+\alpha_3\right)\Big]$$
$$+\sum_{i=1}^{N} \mathbb{E}[z_{i2,\,f=1}] \Big[\log\Gamma\left(x_{i1}+\alpha_2\right)+\log\Gamma\left(x_{i2}+\alpha_3\right)+\log\Gamma\left(x_{i3}+\alpha_4\right)\Big]$$
$$+\sum_{i=1}^{N} \mathbb{E}[z_{i3,\,f=1}] \Big[\log\Gamma\left(x_{i1}+\alpha_3\right)+\log\Gamma\left(x_{i2}+\alpha_4\right)+\log\Gamma\left(x_{i3}+\alpha_5\right)\Big]$$
$$+\sum_{i=1}^{N} \mathbb{E}[z_{i1,\,f=2}] \Big[\log\Gamma\left(x_{i1}+\alpha_3\right)+\log\Gamma\left(x_{i2}+\alpha_2\right)+\log\Gamma\left(x_{i3}+\alpha_1\right)\Big]$$
$$+\sum_{i=1}^{N} \mathbb{E}[z_{i2,\,f=2}] \Big[\log\Gamma\left(x_{i1}+\alpha_4\right)+\log\Gamma\left(x_{i2}+\alpha_3\right)+\log\Gamma\left(x_{i3}+\alpha_2\right)\Big]$$
$$+\sum_{i=1}^{N} \mathbb{E}[z_{i3,\,f=2}] \Big[\log\Gamma\left(x_{i1}+\alpha_5\right)+\log\Gamma\left(x_{i2}+\alpha_4\right)+\log\Gamma\left(x_{i3}+\alpha_3\right)\Big]$$
$$-\sum_{i=1}^{N}\sum_{f=1}^{2} \mathbb{E}[z_{i1f}] \log\Gamma\left(\sum_{j'=1}^{L_{\mathbf{x}}} x_{ij'}+\alpha_1+\alpha_2+\alpha_3\right)$$
$$-\sum_{i=1}^{N}\sum_{f=1}^{2} \mathbb{E}[z_{i2f}] \log\Gamma\left(\sum_{j'=1}^{L_{\mathbf{x}}} x_{ij'}+\alpha_2+\alpha_3+\alpha_4\right)$$
$$-\sum_{i=1}^{N}\sum_{f=1}^{2} \mathbb{E}[z_{i3f}] \log\Gamma\left(\sum_{j'=1}^{L_{\mathbf{x}}} x_{ij'}+\alpha_3+\alpha_4+\alpha_5\right)$$
$$-\sum_{i=1}^{N}\sum_{f=1}^{2} \mathbb{E}[z_{i1f}] \Big[\log\Gamma\left(\alpha_1\right)+\log\Gamma\left(\alpha_2\right)+\log\Gamma\left(\alpha_3\right)\Big]$$
$$-\sum_{i=1}^{N}\sum_{f=1}^{2} \mathbb{E}[z_{i2f}] \Big[\log\Gamma\left(\alpha_2\right)+\log\Gamma\left(\alpha_3\right)+\log\Gamma\left(\alpha_4\right)\Big]$$
$$-\sum_{i=1}^{N}\sum_{f=1}^{2} \mathbb{E}[z_{i3f}] \Big[\log\Gamma\left(\alpha_3\right)+\log\Gamma\left(\alpha_4\right)+\log\Gamma\left(\alpha_5\right)\Big]$$
$$+\sum_{i=1}^{N}\sum_{f=1}^{2} \mathbb{E}[z_{i1f}] \log\Gamma\left(\alpha_1+\alpha_2+\alpha_3\right)$$
$$+\sum_{i=1}^{N}\sum_{f=1}^{2} \mathbb{E}[z_{i2f}] \log\Gamma\left(\alpha_2+\alpha_3+\alpha_4\right)$$
$$+\sum_{i=1}^{N}\sum_{f=1}^{2} \mathbb{E}[z_{i3f}] \log\Gamma\left(\alpha_3+\alpha_4+\alpha_5\right)\ .$$

Considering the terms depending on $\alpha_1$ ($j = 1$), we need to sum over only one shift state $s = 1$:

$$
\begin{aligned}
&\sum_{i=1}^{N} \mathbb{E}[z_{i1,\,f=1}] \log\Gamma\left(x_{i1} + \alpha_1\right) + \sum_{i=1}^{N} \mathbb{E}[z_{i1,\,f=2}] \log\Gamma\left(x_{i3} + \alpha_1\right) \\
&- \sum_{i=1}^{N}\sum_{f=1}^{2} \mathbb{E}[z_{i1f}] \log\Gamma\left(\sum_{j'=1}^{L_{\mathbf{x}}} x_{ij'} + \alpha_1 + \alpha_2 + \alpha_3\right) \\
&- \sum_{i=1}^{N}\sum_{f=1}^{2} \mathbb{E}[z_{i1f}] \log\Gamma\left(\alpha_1\right) + \sum_{i=1}^{N}\sum_{f=1}^{2} \mathbb{E}[z_{i1f}] \log\Gamma\left(\alpha_1 + \alpha_2 + \alpha_3\right) \\
&= \sum_{i=1}^{N} \sum_{s=j-L_{\mathbf{x}}+1}^{j} \mathbb{E}[z_{is,\,f=1}] \log\Gamma\left(x_{i,\,j-s+1} + \alpha_j\right) \\
&\quad + \sum_{i=1}^{N} \sum_{s=j-L_{\mathbf{x}}+1}^{j} \mathbb{E}[z_{is,\,f=2}] \log\Gamma\left(x_{i,\,L_{\mathbf{x}}-j+s} + \alpha_j\right) \\
&\quad + \sum_{i=1}^{N}\sum_{f=1}^{2} \sum_{s=j-L_{\mathbf{x}}+1}^{j} \mathbb{E}[z_{isf}] \Bigg[ -\log\Gamma\left(\sum_{j'=1}^{L_{\mathbf{x}}} x_{ij'} + \sum_{j'=1}^{L_{\mathbf{x}}} \alpha_{j'+s-1}\right) \\
&\qquad\qquad - \left[\log\Gamma\left(\alpha_j\right) - \log\Gamma\left(\sum_{j'=1}^{L_{\mathbf{x}}} \alpha_{j'+s-1}\right)\right]\Bigg].
\end{aligned}
$$

The last rows of the above Equation present how the terms depending on $\alpha_j$ would generalise. When $j = 1$, $j - L_{\mathbf{x}} + 1 = 1 - 3 + 1 = -1$, we sum over shift states $s \in \{-1, 0, 1\}$. Here we have introduced two extra "pseudo" shift states $-1$ and $0$ and we set $\mathbb{E}[z_{isf}] = 0$ when $s \in \{-1, 0\}$.

Considering the terms depending on $\alpha_2$ ($j = 2$), we need to sum over two shift states $s \in \{1, 2\}$:

$$
\begin{aligned}
&\sum_{i=1}^{N} \mathbb{E}[z_{i1,\,f=1}] \log\Gamma\left(x_{i2} + \alpha_2\right) + \sum_{i=1}^{N} \mathbb{E}[z_{i2,\,f=1}] \log\Gamma\left(x_{i1} + \alpha_2\right) \\
&+ \sum_{i=1}^{N} \mathbb{E}[z_{i1,\,f=2}] \log\Gamma\left(x_{i2} + \alpha_2\right) + \sum_{i=1}^{N} \mathbb{E}[z_{i2,\,f=2}] \log\Gamma\left(x_{i3} + \alpha_2\right) \\
&- \sum_{i=1}^{N} \sum_{f=1}^{2} \mathbb{E}[z_{i1f}] \log\Gamma\left(\sum_{j'=1}^{L_{\mathbf{x}}} x_{ij'} + \alpha_1 + \alpha_2 + \alpha_3\right) \\
&- \sum_{i=1}^{N} \sum_{f=1}^{2} \mathbb{E}[z_{i2f}] \log\Gamma\left(\sum_{j'=1}^{L_{\mathbf{x}}} x_{ij'} + \alpha_2 + \alpha_3 + \alpha_4\right) \\
&- \sum_{i=1}^{N} \sum_{f=1}^{2} \mathbb{E}[z_{i1f}] \log\Gamma\left(\alpha_2\right) \\
&- \sum_{i=1}^{N} \sum_{f=1}^{2} \mathbb{E}[z_{i2f}] \log\Gamma\left(\alpha_2\right) \\
&+ \sum_{i=1}^{N} \sum_{f=1}^{2} \mathbb{E}[z_{i1f}] \log\Gamma\left(\alpha_1 + \alpha_2 + \alpha_3\right) \\
&+ \sum_{i=1}^{N} \sum_{f=1}^{2} \mathbb{E}[z_{i2f}] \log\Gamma\left(\alpha_2 + \alpha_3 + \alpha_4\right) \\
&= \sum_{i=1}^{N} \sum_{s=j-L_{\mathbf{x}}+1}^{j} \mathbb{E}[z_{is,\,f=1}] \log\Gamma\left(x_{i,\,j-s+1} + \alpha_j\right) \\
&\quad + \sum_{i=1}^{N} \sum_{s=j-L_{\mathbf{x}}+1}^{j} \mathbb{E}[z_{is,\,f=2}] \log\Gamma\left(x_{i,\,L_{\mathbf{x}}-j+s} + \alpha_j\right) \\
&\quad + \sum_{i=1}^{N} \sum_{f=1}^{2} \sum_{s=j-L_{\mathbf{x}}+1}^{j} \mathbb{E}[z_{isf}] \Bigg[ -\log\Gamma\left(\sum_{j'=1}^{L_{\mathbf{x}}} x_{ij'} + \sum_{j'=1}^{L_{\mathbf{x}}} \alpha_{j'+s-1}\right) \\
&\qquad\qquad - \left[\log\Gamma\left(\alpha_j\right) - \log\Gamma\left(\sum_{j'=1}^{L_{\mathbf{x}}} \alpha_{j'+s-1}\right)\right]\Bigg].
\end{aligned}
$$

The last rows of the above Equation present how the terms depending on $\alpha_j$ would generalise. As $j = 2$, $j - L_{\mathbf{x}} + 1 = 2 - 3 + 1 = 0$, we sum over shift states $s \in \{0, 1, 2\}$. Here we have introduced one extra "pseudo" shift state 0 and set $\mathbb{E}[z_{isf}] = 0$ when $s \in \{0\}$.

Considering terms depending on $\alpha_3$ ($j = 3$), we need to sum over shift states $s \in \{1, 2, 3\}$:

$$\sum_{i=1}^{N} \mathbb{E}[z_{i1,\,f=1}]) + \log\Gamma\left(x_{i3} + \alpha_3\right) + \sum_{i=1}^{N} \mathbb{E}[z_{i2,\,f=1}] \log\Gamma\left(x_{i2} + \alpha_3\right) + \sum_{i=1}^{N} \mathbb{E}[z_{i3,\,f=1}] \log\Gamma\left(x_{i1} + \alpha_3\right)$$
$$+ \sum_{i=1}^{N} \mathbb{E}[z_{i1,\,f=2}] \log\Gamma\left(x_{i1} + \alpha_3\right) + \sum_{i=1}^{N} \mathbb{E}[z_{i2,\,f=2}] \log\Gamma\left(x_{i2} + \alpha_3\right) + \sum_{i=1}^{N} \mathbb{E}[z_{i3,\,f=2}] \log\Gamma\left(x_{i3} + \alpha_3\right)$$
$$- \sum_{i=1}^{N} \sum_{f=1}^{2} \mathbb{E}[z_{i1f}] \log\Gamma\left(\sum_{j'=1}^{L_{\mathbf{x}}} x_{ij'} + \alpha_1 + \alpha_2 + \alpha_3\right)$$
$$- \sum_{i=1}^{N} \sum_{f=1}^{2} \mathbb{E}[z_{i2f}] \log\Gamma\left(\sum_{j'=1}^{L_{\mathbf{x}}} x_{ij'} + \alpha_2 + \alpha_3 + \alpha_4\right)$$
$$- \sum_{i=1}^{N} \sum_{f=1}^{2} \mathbb{E}[z_{i3f}] \log\Gamma\left(\sum_{j'=1}^{L_{\mathbf{x}}} x_{ij'} + \alpha_3 + \alpha_4 + \alpha_5\right)$$
$$- \sum_{i=1}^{N} \sum_{f=1}^{2} \mathbb{E}[z_{i1f}] \log\Gamma\left(\alpha_3\right) - \sum_{i=1}^{N} \sum_{f=1}^{2} \mathbb{E}[z_{i2f}] \log\Gamma\left(\alpha_3\right) - \sum_{i=1}^{N} \sum_{f=1}^{2} \mathbb{E}[z_{i3f}] \log\Gamma\left(\alpha_3\right)$$
$$+ \sum_{i=1}^{N} \sum_{f=1}^{2} \mathbb{E}[z_{i1f}] \log\Gamma\left(\alpha_1 + \alpha_2 + \alpha_3\right)$$
$$+ \sum_{i=1}^{N} \sum_{f=1}^{2} \mathbb{E}[z_{i2f}] \log\Gamma\left(\alpha_2 + \alpha_3 + \alpha_4\right)$$
$$+ \sum_{i=1}^{N} \sum_{f=1}^{2} \mathbb{E}[z_{i3f}] \log\Gamma\left(\alpha_3 + \alpha_4 + \alpha_5\right)$$
$$= \sum_{i=1}^{N} \sum_{s=j-L_{\mathbf{x}}+1}^{j} \mathbb{E}[z_{is,\,f=1}] \log\Gamma\left(x_{i,\,j-s+1} + \alpha_j\right)$$
$$+ \sum_{i=1}^{N} \sum_{s=j-L_{\mathbf{x}}+1}^{j} \mathbb{E}[z_{is,\,f=2}] \log\Gamma\left(x_{i,\,L_{\mathbf{x}}-j+s} + \alpha_j\right)$$
$$+ \sum_{i=1}^{N} \sum_{f=1}^{2} \sum_{s=j-L_{\mathbf{x}}+1}^{j} \mathbb{E}[z_{isf}] \Bigg[ -\log\Gamma\left(\sum_{j'=1}^{L_{\mathbf{x}}} x_{ij'} + \sum_{j'=1}^{L_{\mathbf{x}}} \alpha_{j'+s-1}\right) - \left[\log\Gamma\left(\alpha_j\right) - \log\Gamma\left(\sum_{j'=1}^{L_{\mathbf{x}}} \alpha_{j'+s-1}\right)\right]\Bigg].$$

The last rows of the above Equation present how the terms depending on $\alpha_j$ would generalise. As $j = 3$, $j - L_{\mathbf{x}} + 1 = 3 - 3 + 1 = 1$, we sum over shift states $s \in \{1, 2, 3\}$. Here we do not have to add extra "pseudo" shift states.

Considering terms depending on $\alpha_4$ ($j = 4$), we need to sum over shift states $s \in \{2, 3\}$:

$$
\begin{aligned}
&\sum_{i=1}^{N} \mathbb{E}[z_{i2,\,f=1}] \log\Gamma\left(x_{i3} + \alpha_4\right) + \sum_{i=1}^{N} \mathbb{E}[z_{i3,\,f=1}] \log\Gamma\left(x_{i2} + \alpha_4\right) \\
&+ \sum_{i=1}^{N} \mathbb{E}[z_{i2,\,f=2}] \log\Gamma\left(x_{i1} + \alpha_4\right) + \sum_{i=1}^{N} \mathbb{E}[z_{i3,\,f=2}] \log\Gamma\left(x_{i2} + \alpha_4\right)] \\
&- \sum_{i=1}^{N}\sum_{f=1}^{2} \mathbb{E}[z_{i2f}] \log\Gamma\left(\sum_{j'=1}^{L_{\mathbf{x}}} x_{ij'} + \alpha_2 + \alpha_3 + \alpha_4\right) \\
&- \sum_{i=1}^{N}\sum_{f=1}^{2} \mathbb{E}[z_{i3f}] \log\Gamma\left(\sum_{j'=1}^{L_{\mathbf{x}}} x_{ij'} + \alpha_3 + \alpha_4 + \alpha_5\right) \\
&- \sum_{i=1}^{N}\sum_{f=1}^{2} \mathbb{E}[z_{i2f}] \log\Gamma\left(\alpha_4\right) - \sum_{i=1}^{N}\sum_{f=1}^{2} \mathbb{E}[z_{i3f}] \log\Gamma\left(\alpha_4\right) \\
&+ \sum_{i=1}^{N}\sum_{f=1}^{2} \mathbb{E}[z_{i2f}] \log\Gamma\left(\alpha_2 + \alpha_3 + \alpha_4\right) \\
&+ \sum_{i=1}^{N}\sum_{f=1}^{2} \mathbb{E}[z_{i3f}] \log\Gamma\left(\alpha_3 + \alpha_4 + \alpha_5\right) \\
&= \sum_{i=1}^{N} \sum_{s=j-L_{\mathbf{x}}+1}^{j} \mathbb{E}[z_{is,\,f=1}] \log\Gamma\left(x_{i,\,j-s+1} + \alpha_j\right) \\
&\quad + \sum_{i=1}^{N} \sum_{s=j-L_{\mathbf{x}}+1}^{j} \mathbb{E}[z_{is,\,f=2}] \log\Gamma\left(x_{i,\,L_{\mathbf{x}}-j+s} + \alpha_j\right) \\
&\quad + \sum_{i=1}^{N}\sum_{f=1}^{2} \sum_{s=j-L_{\mathbf{x}}+1}^{j} \mathbb{E}[z_{isf}] \Bigg[ -\log\Gamma\left(\sum_{j'=1}^{L_{\mathbf{x}}} x_{ij'} + \sum_{j'=1}^{L_{\mathbf{x}}} \alpha_{j'+s-1}\right) \\
&\qquad\qquad - \left[\log\Gamma\left(\alpha_j\right) - \log\Gamma\left(\sum_{j'=1}^{L_{\mathbf{x}}} \alpha_{j'+s-1}\right)\right]\Bigg].
\end{aligned}
$$

The last rows of the above Equation present how the terms depending on $\alpha_j$ would generalise. As $j = 4$, $j - L_{\mathbf{x}} + 1 = 4 - 3 + 1 = 2$, we sum over shift states $s \in \{2, 3, 4\}$. Here we have introduced one extra "pseudo" shift state 4; $\mathbb{E}[z_{isf}] = 0$ when $s \in \{4\}$.

Considering terms depending on $\alpha_5$ ($j = 5$), we sum over the shift state $s = 3$

$$
\begin{aligned}
&\sum_{i=1}^{N} \mathbb{E}[z_{i3,\,f=1}] \log\Gamma\left(x_{i3} + \alpha_5\right) + \sum_{i=1}^{N} \mathbb{E}[z_{i3,\,f=2}] \log\Gamma\left(x_{i1} + \alpha_5\right) \\
&- \sum_{i=1}^{N}\sum_{f=1}^{2} \mathbb{E}[z_{i3f}] \log\Gamma\left(\sum_{j'=1}^{L_{\mathbf{x}}} x_{ij'} + \alpha_3 + \alpha_4 + \alpha_5\right) \\
&- \sum_{i=1}^{N}\sum_{f=1}^{2} \mathbb{E}[z_{i3f}] \log\Gamma\left(\alpha_5\right) \\
&+ \sum_{i=1}^{N}\sum_{f=1}^{2} \mathbb{E}[z_{i3f}] \log\Gamma\left(\alpha_3 + \alpha_4 + \alpha_5\right) \\
&= \sum_{i=1}^{N} \sum_{s=j-L_{\mathbf{x}}+1}^{j} \mathbb{E}[z_{is,\,f=1}] \log\Gamma\left(x_{i,\,j-s+1} + \alpha_j\right) \\
&\quad + \sum_{i=1}^{N} \sum_{s=j-L_{\mathbf{x}}+1}^{j} \mathbb{E}[z_{is,\,f=2}] \log\Gamma\left(x_{i,\,L_{\mathbf{x}}-j+s} + \alpha_j\right) \\
&\quad + \sum_{i=1}^{N}\sum_{f=1}^{2} \sum_{s=j-L_{\mathbf{x}}+1}^{j} \mathbb{E}[z_{isf}] \Bigg[ -\log\Gamma\left(\sum_{j'=1}^{L_{\mathbf{x}}} x_{ij'} + \sum_{j'=1}^{L_{\mathbf{x}}} \alpha_{j'+s-1}\right) \\
&\qquad\qquad - \left[\log\Gamma\left(\alpha_j\right) - \log\Gamma\left(\sum_{j'=1}^{L_{\mathbf{x}}} \alpha_{j'+s-1}\right)\right]\Bigg].
\end{aligned}
$$

The last rows of the above Equation present how the terms depending on $\alpha_j$ would generalise. As $j = 5$, $j - L_{\mathbf{x}} + 1 = 5 - 3 + 1 = 3$, we sum over shift states $s \in \{3, 4, 5\}$. Here we have introduced two extra "pseudo" shift states 4 and 5; $\mathbb{E}[z_{isf}] = 0$ when $s \in \{4, 5\}$.

We conclude that the lower bound terms (omitting the prior terms) that depend on $\alpha_j$ generalise for all $j \in \{1, \ldots, L_{\boldsymbol{\alpha}}\}$, $s \in \{1, \ldots, S\}$ and $L_{\mathbf{x}}$ as follows:

$$
\begin{aligned}
&\sum_{i=1}^{N} \sum_{s=j-L_{\mathbf{x}}+1}^{j} \mathbb{E}[z_{is,\,f=1}] \log\Gamma\left(x_{i,\,j-s+1} + \alpha_j\right) \\
&+ \sum_{i=1}^{N} \sum_{s=j-L_{\mathbf{x}}+1}^{j} \mathbb{E}[z_{is,\,f=2}] \log\Gamma\left(x_{i,\,L_{\mathbf{x}}-j+s} + \alpha_j\right) \\
&+ \sum_{i=1}^{N}\sum_{f=1}^{2} \sum_{s=j-L_{\mathbf{x}}+1}^{j} \mathbb{E}[z_{isf}] \Bigg[ -\log\Gamma\left(\sum_{j'=1}^{L_{\mathbf{x}}} x_{ij'} + \sum_{j'=1}^{L_{\mathbf{x}}} \alpha_{j'+s-1}\right) \\
&\qquad\qquad - \left[\log\Gamma\left(\alpha_j\right) - \log\Gamma\left(\sum_{j'=1}^{L_{\mathbf{x}}} \alpha_{j'+s-1}\right)\right]\Bigg],
\end{aligned}
\tag{S26}
$$

where $\mathbb{E}[z_{iksf}] = 0$, when $s \in \{j - L_{\mathbf{x}} + 1, \ldots, 0, S + 1, \ldots L_{\boldsymbol{\alpha}}\}$. Here we have introduced $L_{\mathbf{x}} - 1$ extra "pseudo" shift states in both ends of the shift state vector, resulting in a total of $2 \times (L_{\mathbf{x}} - 1)$ "pseudo" shift states.

Now the terms of the lower bound in Equation S25 that depend on $\alpha_{kj}^{(m)}$ are

$$
\begin{aligned}
&(\eta_h - 1)\log h_k^{(m)} - \nu_h h_k^{(m)} + (\eta - 1)\log \alpha_{kj}^{(m)} - \nu \alpha_{kj}^{(m)} \\
&+ \sum_{i=1}^{N} \sum_{s=j-L_{\mathbf{x}}+1}^{j} \mathbb{E}[z_{iks,\,f=1}] \log\Gamma\left(x_{i,\,j-s+1}^{(m)} + \alpha_{kj}^{(m)}\right) \\
&+ \sum_{i=1}^{N} \sum_{s=j-L_{\mathbf{x}}+1}^{j} \mathbb{E}[z_{iks,\,f=2}] \log\Gamma\left(x_{i,\,L_{\mathbf{x}}-j+s}^{(m)} + \alpha_{kj}^{(m)}\right) \\
&+ \sum_{i=1}^{N} \sum_{f=1}^{2} \sum_{s=j-L_{\mathbf{x}}+1}^{j} \mathbb{E}[z_{iksf}] \Bigg[ -\log\Gamma\left( \sum_{j'=1}^{L_{\mathbf{x}}} x_{ij'}^{(m)} + \sum_{j'=1}^{L_{\mathbf{x}}} \alpha_{k,\,j'+s-1}^{(m)} \right) \\
&\qquad\qquad - \left[ \log\Gamma\left(\alpha_{kj}^{(m)}\right) - \log\Gamma\left( \sum_{j'=1}^{L_{\mathbf{x}}} \alpha_{k,\,j'+s-1}^{(m)} \right) \right] \Bigg],
\end{aligned}
$$

where $\mathbb{E}[z_{iksf}] = 0$, when $s \in \{j - L_{\mathbf{x}} + 1, \ldots, 0, S + 1, \ldots, L_{\boldsymbol{\alpha}}\}$.

Now it is straightforward to derive the derivative of the lower bound wrt. $\alpha_{kj}^{(m)}$ as

$$
\begin{aligned}
\frac{\partial Q(\boldsymbol{\theta}, \boldsymbol{\theta}^{\text{old}})}{\partial \alpha_{kj}^{(m)}} =&(\eta_h - 1)\frac{g_{kj}^{(m)}}{h_k^{(m)}} - \nu_h g_{kj}^{(m)} + \frac{(\eta - 1)}{\alpha_{kj}^{(m)}} - \nu \\
&+ \sum_{i=1}^{N} \sum_{s=j-L_{\mathbf{x}}+1}^{j} \mathbb{E}[z_{iks,\,f=1}] \psi\left(x_{i,\,j-s+1}^{(m)} + \alpha_{kj}^{(m)}\right) \\
&+ \sum_{i=1}^{N} \sum_{s=j-L_{\mathbf{x}}+1}^{j} \mathbb{E}[z_{iks,\,f=2}] \psi\left(x_{i,\,L_{\mathbf{x}}-j+s}^{(m)} + \alpha_{kj}^{(m)}\right) \\
&+ \sum_{i=1}^{N} \sum_{f=1}^{2} \sum_{s=j-L_{\mathbf{x}}+1}^{j} \mathbb{E}[z_{iksf}] \Bigg[ -\psi\left( \sum_{j'=1}^{L_{\mathbf{x}}} x_{ij'}^{(m)} + \sum_{j'=1}^{L_{\mathbf{x}}} \alpha_{k,\,j'+s-1}^{(m)} \right) \\
&\qquad\qquad - \left[ \psi\left(\alpha_{kj}^{(m)}\right) - \psi\left( \sum_{j'=1}^{L_{\mathbf{x}}} \alpha_{k,\,j'+s-1}^{(m)} \right) \right] \Bigg]
\end{aligned}
$$

$$
g_{kj}^{(m)} = \begin{cases} 2(\alpha_{kj}^{(m)} - \alpha_{k,\,j+1}^{(m)}) & \text{for } j = 1 \\ 2(\alpha_{kj}^{(m)} - \alpha_{k,\,j-1}^{(m)}) & \text{for } j = L_{\boldsymbol{\alpha}} \\ 2(2\alpha_{kj}^{(m)} - \alpha_{k,\,j+1}^{(m)} - \alpha_{k,\,j-1}^{(m)}) & \text{otherwise}\,, \end{cases}
$$

where $\mathbb{E}[z_{iksf}] = 0$, when $s \in \{j - L_{\mathbf{x}} + 1, \ldots, 0, S+1, \ldots, L_{\boldsymbol{\alpha}}\}$.

##### S2.6 The derivative of the lower bound wrt. $\lambda_{kj}^{(m)}$

The derivative of lower bound in Equation S25 wrt. $\lambda_{kj}^{(m)}$ is

$$\begin{aligned}
\frac{\partial Q(\boldsymbol{\theta}, \boldsymbol{\theta}^{\text{old}})}{\partial \lambda_{kj}^{(m)}} =& \alpha_{kj}^{(m)} \left( (\eta_h - 1) \frac{g_{kj}^{(m)}}{h_k^{(m)}} - \nu_h g_{kj}^{(m)} + \frac{(\eta - 1)}{\alpha_{kj}^{(m)}} - \nu \right) \\
&+ \alpha_{kj}^{(m)} \sum_{i=1}^{N} \sum_{s=j-L_{\mathbf{x}}+1}^{j} \mathbb{E}[z_{iks,\, f=1}] \psi \left( x_{i,\, j-s+1}^{(m)} + \alpha_{kj}^{(m)} \right) \\
&+ \alpha_{kj}^{(m)} \sum_{i=1}^{N} \sum_{s=j-L_{\mathbf{x}}+1}^{j} \mathbb{E}[z_{iks,\, f=2}] \psi \left( x_{i,\, L_{\mathbf{x}}-j+s}^{(m)} + \alpha_{kj}^{(m)} \right) \\
&+ \alpha_{kj}^{(m)} \sum_{i=1}^{N} \sum_{f=1}^{2} \sum_{s=j-L_{\mathbf{x}}+1}^{j} \mathbb{E}[z_{iksf}] \Bigg[ -\psi \left( \sum_{j'=1}^{L_{\mathbf{x}}} x_{ij'}^{(m)} + \sum_{j'=1}^{L_{\mathbf{x}}} \alpha_{k,\, j'+s-1}^{(m)} \right) \\
&\qquad - \left[ \psi \left( \alpha_{kj}^{(m)} \right) - \psi \left( \sum_{j'=1}^{L_{\mathbf{x}}} \alpha_{k,\, j'+s-1}^{(m)} \right) \right] \Bigg]
\end{aligned} \tag{S27}$$

$$g_{kj}^{(m)} = \begin{cases} 2(\alpha_{kj}^{(m)} - \alpha_{k,\, j+1}^{(m)}) & \text{for } j = 1 \\ 2(\alpha_{kj}^{(m)} - \alpha_{k,\, j-1}^{(m)}) & \text{for } j = L_{\boldsymbol{\alpha}} \\ 2(2\alpha_{kj}^{(m)} - \alpha_{k,\, j+1}^{(m)} - \alpha_{k,\, j-1}^{(m)}) & \text{otherwise}\,, \end{cases}$$

where $\mathbb{E}[z_{iks,\, f=1}] = 0$, when $s \in \{j - L_{\mathbf{x}} + 1, \ldots, 0, S+1, \ldots, L_{\boldsymbol{\alpha}}\}$.

##### S2.7 The EM updates for cluster assignments $\mathbb{E}[z_{iksf}]$

The cluster assignments $\mathbb{E}[z_{iksf}] = p(z_{iksf} = 1 | \mathbf{x}_i^*, \boldsymbol{\theta}^{\text{old}})$ are computed as

$$\begin{aligned}
p(z_{iksf} = 1 | \mathbf{x}_i^*, \boldsymbol{\theta}^{\text{old}}) &= \frac{p(z_{iksf} = 1 | \boldsymbol{\theta}^{\text{old}}) p(\mathbf{x}_i^* | z_{iksf} = 1, \boldsymbol{\theta}^{\text{old}})}{\sum_{k'=1}^{K} \sum_{s'=1}^{S} \sum_{f'=1}^{2} p(z_{ik's'f'} = 1 | \boldsymbol{\theta}^{\text{old}}) p(\mathbf{x}_i^* | z_{ik's'f'} = 1, \boldsymbol{\theta}^{\text{old}})} \\
&= \frac{\pi_k^{\text{old}} \xi_{is} \zeta_{if} \prod_{m=1}^{M} \text{Dirichlet-multinomial}(\mathbf{x}_i^{(m)} | \alpha_{ksf}^{(m,\, \text{old})})}{\sum_{k'=1}^{K} \sum_{s'=1}^{S} \sum_{f'=1}^{2} \pi_{k'}^{\text{old}} \xi_{is'} \zeta_{if'} \prod_{m=1}^{M} \text{Dirichlet-multinomial}(\mathbf{x}_i^{(m)} | \alpha_{k's'f'}^{(m,\, \text{old})})}\,.
\end{aligned}$$

##### S2.8 The EM updates for mixture weights $\boldsymbol{\pi}$

The mixing weights are updated as

$$\pi_k = \frac{\sum_{i=1}^{N}\sum_{s=1}^{S}\sum_{f=1}^{2}\mathbb{E}[z_{iksf}]}{\sum_{i=1}^{N}\sum_{k'=1}^{K}\sum_{s=1}^{S}\sum_{f=1}^{2}\mathbb{E}[z_{ik'sf}]}\,.$$

##### S2.9 Checking the EM convergence

Computing the negative log unnormalised posterior given the current parameter estimates is utilised to check the convergence of the EM algorithm:

$$\begin{aligned}
-\log p(\boldsymbol{\alpha}^*,\boldsymbol{\pi}) \propto& -\log p(\mathbf{X}|\boldsymbol{\alpha}^*,\boldsymbol{\pi}) - \log p(\boldsymbol{\alpha}^*)\\
=& -\sum_{i=1}^{N}\log\left\{\sum_{k=1}^{K}\pi_k\sum_{s=1}^{S}\xi_{is}\sum_{f=1}^{2}\zeta_{if}\exp\left[\sum_{m=1}^{M}\log\text{Dirichlet-multinomial}(\mathbf{x}_i^{(m)}|\boldsymbol{\alpha}_{ksf}^{(m)})\right]\right\}\\
&-\sum_{m=1}^{M}\sum_{k=1}^{K}\log\Gamma(h_k^{(m)}|\eta_h,\nu_h)\\
&-\sum_{m=1}^{M}\sum_{k=1}^{K}\sum_{j=1}^{L_{\boldsymbol{\alpha}}}\log\Gamma(\alpha_{kj}^{(m)}|\eta,\nu)\\
=& -\sum_{i=1}^{N}\log\Bigg\{\sum_{k=1}^{K}\pi_k\sum_{s=1}^{S}\xi_{is}\sum_{f=1}^{2}\zeta_{if}\exp\Bigg[\sum_{m=1}^{M}\Bigg(\log\Gamma(J_i^{(m)}+1)+\log\Gamma(\sum_{j=1}^{L_{\mathbf{x}}}\alpha_{kjsf}^{(m)})\\
&+\sum_{j=1}^{L_{\mathbf{x}}}\log\Gamma(x_{ij}^{(m)}+\alpha_{kjsf}^{(m)})\\
&-\log\Gamma(\sum_{j=1}^{L_{\mathbf{x}}}x_{ij}^{(m)}+\sum_{j=1}^{L_{\mathbf{x}}}\alpha_{kjsf}^{(m)})-\sum_{j=1}^{L_{\mathbf{x}}}\log\Gamma(x_{ij}^{(m)}+1)-\sum_{j=1}^{L_{\mathbf{x}}}\log\Gamma(\alpha_{kjsf}^{(m)})\Bigg)\Bigg]\Bigg\}\\
&-MK\eta_h\log\nu_h-(\eta_h-1)\sum_{m=1}^{M}\sum_{k=1}^{K}\log h_k^{(m)}+\nu_h\sum_{m=1}^{M}\sum_{k=1}^{K}h_k^{(m)}+MK\log\Gamma(\eta_h)\\
&-MKL_{\boldsymbol{\alpha}}\eta\log\nu-(\eta-1)\sum_{m=1}^{M}\sum_{k=1}^{K}\sum_{j=1}^{L_{\boldsymbol{\alpha}}}\log\alpha_{kj}^{(m)}+\nu\sum_{m=1}^{M}\sum_{k=1}^{K}\sum_{j=1}^{L_{\boldsymbol{\alpha}}}\alpha_{kj}^{(m)}+MKL_{\boldsymbol{\alpha}}\log\Gamma(\eta)\,.
\end{aligned}$$

### S3 Model selection: choosing the number of clusters

#### S3.1 Calculation of BIC

Bayesian information criterion (BIC) is computed as

$$\mathrm{BIC} = -2 \log \hat{\mathcal{L}} + k \log(n)\,,$$

where $\hat{\mathcal{L}}$, $k$ and $n$ represent the likelihood, the number of parameters in the model, and the number of observations, respectively. Since Bayesian approach was adopted in this study, $\hat{\mathcal{L}}$ represents the posterior rather than the likelihood. Here, $k \log(n)$ acts as a penalty term depending on the number of model parameters so that more complex models are penalised more strongly.

The number of parameters $k$ in the model depends on whether the model infers the shift states or not. Given the number of chromatin features $M$, the number of clusters $K$, and the length $L_{\mathbf{x}}$ of the profile $\mathbf{x}^{(m)}$, for the model that do not infer the shift states, the number of parameters is

$$k = L_{\mathbf{x}} \times K \times M + (K - 1)\,,$$

where $L_{\mathbf{x}} \times K \times M$ denotes the number of Dirichlet parameters and $K - 1$ represents $K$ mixture weights ($\boldsymbol{\pi}$) that sum up to one.

For the models that infers the shift states, the length of Dirichlet parameters is $L_{\boldsymbol{\alpha}}$ (see Suppl. Section S2), and thus the number of model parameters is

$$k = L_{\boldsymbol{\alpha}} \times K \times M + (K - 1)\,.$$

#### S3.2 Calculation of AIC

Akaike information criterion (AIC) is computed as

$$\mathrm{AIC} = -\log \hat{\mathcal{L}} + k\,,$$

where $\hat{\mathcal{L}}$, $k$ are the same as described in Section S3.1. The number of clusters is selected by computing AIC and BIC on different models inferred by varying the number of clusters, and selecting the cluster number that result to the smallest AIC or BIC value.

### S4 Supplementary Methods

#### S4.1 Soft k-means clustering algorithm

Soft k-means clustering with kmeans++ initialisation is utilised to obtain starting values for the ChromDMM Expectation-Maximisation algorithm. K-means results in initial estimates of the ChromDMM component parameters and the cluster assignments. In kmeans++ initialisation, the idea is to weight the data points according to their squared distance from the closest cluster center already chosen. Instead of picking centers randomly, after picking one, the probability of picking a more distant point to the original center is larger. kmeans++ has been demonstrated to improve both the speed and the accuracy of k-means (MacKay, 2003).

For the kmeans++ clustering, the $N \times L_{\mathbf{x}}$ ($L_{\mathbf{x}}$ is the length of $\mathbf{x}^{(m)}$) count matrices of a given chromatin feature were normalised row-wise. The sum of each row of the matrix was computed, and if not zero, the elements of the row were divided by the sum. This was performed separately for each chromatin feature matrix. Then the individual chromatin features were concatenated and clustered by kmeans++. The resulting cluster centers were utilised as initial estimates for the $\boldsymbol{\alpha}_k^{(m)}$ parameters.

**Algorithm S1:** Soft k-means clustering algorithm.

1. Cluster centers, denoted as $m^{(k)}$, were initialised using kmeans++ algorithm.

2. Cluster membership probabilities of the data points were calculated as:

$$r_k^{(n)} = \frac{\exp(-\beta\, d(m^{(k)}, x^{(n)}))}{\sum_{k'} \exp(-\beta\, d(m^{(k')}, x^{(n)}))}$$

   where $r_k^{(n)}$ indicated the probability that data point $x^{(n)}$ was a member of cluster $k$. The parameter $\beta$ defined the stiffness; as $\beta$ approaches infinity, the algorithm is identical to hard k-means clustering. The function $d(\cdot)$ defined the distance. As a distance function, the Euclidean distance was utilised.

3. The cluster centers were computed as:

$$m^{(k)} = \frac{\sum_n r_k^{(n)} x^{(n)}}{\sum_n r_k^{(n)}}$$

4. If the sum of differences between the previous cluster centers and last cluster centers were negligible stop, otherwise go to step 2.

#### S4.2 Chromatin feature data from ENCODE, enhancer definition and data preprocessing

This section describes the definition of enhancer elements in the K562 cell line and the formation of chromatin feature signals in the enhanser regions. To define the genomic coordinates of the enhancer elements, the p300 ChIP-seq peaks and DNase I HS peaks for the K562 cell line were downloaded from the ENCODE project (The ENCODE Project Consortium, 2012;

Davis *et al.*, 2018). To prevent including promoters in the set of enhancers, the TSS coordinates for protein-coding genes were obtained from GENCODE. The enhancers were defined as the TSS-distal p300 ChIP-seq peak summits overlapping a DNase I hypersensitivity peak quantified by DNase-seq. The enhancers that overlapped with ENCODE blacklist regions were excluded. The 1000 most significant enhancers (based on q-values of the p300 ChIP-seq peaks) were selected for clustering. In addition, we downloaded data for ten chromatin features (H2AZ, H3K27ac H3K4me1, H3K4me2, H3K4me3, H3K79me2, H3K9ac, RNAPOL2, DNase-seq, MNase-seq) for the K562 cell line from ENCODE. For every enhancer, defined as a 2kb region centered at the anchor point, the 5' ends of chromatin feature reads aligning to the region were counted in bins of 40 bp, resulting in profiles of length $L = 50$. The ENCODE accessions for the chromatin feature data sets and peak files are provided in Suppl. Table S1. The GENCODE annotation and ENCODE blacklist regions accessions are provided in Suppl. Table S2.

The chromatin feature ChIP-seq, DNase-seq and MNase-seq data were pre-processed with the following steps, similarly as in (Osmala and Lähdesmäki, 2020):

1. Raw reads were aligned to the human genome version hg19 with Bowtie 2 (Langmead and Salzberg, 2012) (bowtie2-2.3.3.1) adopting the default options.

2. Polymerase Chain Reaction (PCR) duplicates were removed (Marx, 2017).

3. Possible isogenic replicates were pooled.

4. The fragment lengths for ChIP-sed reads were estimated from the cross-correlation profiles obtained with phantompeakqualtools (spp version 1.14) (Landt *et al.*, 2012; Kharchenko *et al.*, 2008).

5. The ChIP-seq reads were shifted by the half of the estimated fragment length with the combination of bedtools2 and samtools, In addition, the MNase-seq reads were shifted by 149/2, the half of the length of DNA wrapping around a nucleosome ($\sim$ 149 bps). In contrast, the DNase-seq and control reads were not shifted.

6. The genome-wide coverage signals were generated. When creating the coverage signal, the control coverage was normalised wrt. the ChIP coverage to equalise the library sizes, and the control coverage was subtracted from the ChIP coverage. The normalisation was performed as previously described (Li *et al.*, 2011; Landt *et al.*, 2012; Le Martelot *et al.*, 2012).

7. The control signal was not subtracted from the DNase-seq and MNase-seq signals.

8. The coverage was computed in 40 bp bins, the coverage values were rounded to the nearest integer, and the negative values were converted to zero.

#### S4.3 GAT analysis for the enrichment of transcriptional regulatory factor (TRF) ChIP-seq peaks at enhancer clusters

The enhancer clusters revealed by ChromDMM, ChIP-Partitioning, and SPar-K were investigated for the enrichment of the binding sites of transcription factors (TFs) and other regulatory proteins, collectively referred to as transcriptional regulatory factors (TRFs). To this end, we downloaded the optimal IDR thresholded TRF ChIP-seq peaks in the K562 cell line produced by the ENCODE Processing Pipeline (The ENCODE Project Consortium, 2012; Davis *et al.*, 2018). The peak files with status archived were excluded from the analysis, and the most recent file for each TRF was selected. The peak files of RNA polymerases, p300 and CTCF were excluded from the peak list resulting in a large collection of 220 unique TRF peak sets. The ENCODE TRF ChIP-seq peak file accessions are provided in Suppl. Table S1.

The TRF enrichment analysis for the enhancer clusters was performed using GAT tool. The genomic coordinates of the enhancers were realigned according to the inferred shift states, and genomic windows of size 50 bp centred at the enhancer middle coordinates were defined. First, GAT computes the overlap between the TRF intervals (of varying size) and the enhancer intervals (of size 50bp) in nucleotides. Then GAT simulates randomly placed intervals in the genome with the same size distribution as the TRF peaks within the mappable genome. The mappable genome was defined utilising the sequence alignability file with ENCODE identifier ENCF000EHJ (see Suppl. Table S2), and the regions with mappability score less than 1 were excluded. The overlap between the enhancer cluster and the simulated set was recorded. The simulation was repeated multiple times, and the average overlap over all simulations was computed as the expected overlap. The ratio of the observed and the expected overlap denotes the fold enrichment. The fold enrichments in log2 scale were collected in a $220 \times 6$ matrix, which was further clustered by rows and columns, and visualised as a heatmap utilising the `ComplexHeatmap R` package (Gu *et al.*, 2016). In addition, an empirical p-value for the observed overlap was computed from the simulated null distribution. Multiple testing correction for the p-values was performed utilising Benjamini–Hochberg False discovery rate (FDR) method to obtain q-values (Benjamini and Hochberg, 1995). In addition to the TRF abbreviations, more descriptive names of TRFs were queried from Panther (Mi *et al.*, 2020) and listed in Figure 4 and Suppl. Figures and S17, S20, and S21. The biological functions of selected TRFs were further investigated employing Factorbook (Wang *et al.*, 2012).

#### S4.4 Simulated data generation

Data for four chromatin features (H3K4me1, H3K27ac, RNA polymerase II, and MNase-seq, $M = 4$) at 1000 enhancers were obtained from the ENCODE project as described in Suppl. Section S4.2. The data were clustered with ChromDMM by requiring inference of both the shift and flip states. The length of a profile $\mathbf{x}^{(m)}$ was $L_{\mathbf{x}} = 50$ and the number of shift states was $S = 21$. From the fitted model, we chose the Dirichlet parameters $\boldsymbol{\alpha}_k^{(m)}$ for two clusters ($K = 2$). The Dirichlet parameters $\boldsymbol{\alpha}_k^{(m)}$ of length $L_{\boldsymbol{\alpha}} = L_{\mathbf{x}} + S - 1 = 70$ ($k \in 1, \ldots, K$ and $m \in 1, \ldots, M$) were employed to generatively sample data as presented in Suppl. Algorithm S2.

The Dirichlet-parameters were further extended to length $L_{\boldsymbol{\alpha}} + S - 1 = 90$ enabling to create shifted data profiles of length 70 and 50. The shifted profiles of length 50 were clustered with ChromDMM requiring the inference of the shift states, and the shifted profiles of length 70 were realigned according to the inferred shift states. To extend the Dirichlet parameters to length 90, the extensions on both ends received the minimum value from the Dirichlet parameter values upstream or downstream the enhancer center position.

**Algorithm S2:** Simulated data generation

1 **for** *each chromatin feature* $m \leftarrow 1$ **to** $M$ **do**
2     $n_m \in \{10, 20, 50, 100\}$ `// Chromatin-feature-specific coverage`
3     **for** *each cluster* $k \leftarrow 1$ **to** $K$ **do**
4         **for** *each sample* $i \leftarrow 1$ **to** $N_k$ **do**
5             `// Sample multinomial parameter from the Dirichlet distribution`
6             $\mathbf{p}_{ik}^{(m)} \sim \text{Dirichlet}(\boldsymbol{\alpha}_k^{(m)})$
7             `// Sample profile` $\mathbf{x}_i^{(m)}$
8             $\mathbf{x}_{ik}^{(m)} \sim \text{Multinomial}(\mathbf{p}_{ik}^{(m)}, n_m)$
9         **end**
10     **end**
11 **end**
12 **return** $\mathbf{X}^*$

The data simulation proceeds as follows: Given the Dirichlet parameters $\boldsymbol{\alpha}_k^{(m)}$, the Multinomial parameters $\mathbf{p}_{ik}^{(m)}$ for each genomic locus $i$ were sampled from the Dirichlet distribution (row 6 in Suppl. Algorithm S2). The profiles $\mathbf{x}_i^{(m)}$ were further generated from the Multinomial distribution given the chromatin-feature-specific coverage $n_m$ (row 8 in Suppl. Algorithm S2). The length of the profiles $\mathbf{x}_i^{(m)}$ was 90. In ChromDMM experiments without the shift state inference, the middle 50 elements of the profiles were extracted. In contrast, in ChromDMM experiments with the shift state inference, the middle 70 elements of the profiles were extracted. In the simulated data experiments, the number of clusters was $K = 2$, and the number of samples belonging to cluster $k$, $N_k$, was 500 for both clusters. In addition, the number of chromatin features $M$ varied between one, two, three and four. The data generation presented in Suppl. Algorithm S2 was repeated 100 times with unique random seed for each repetition. Visualisation of one instance of the simulated data containing all four chromatin features is presented in Suppl. Figure S4. In the example, the coverage for all four chromatin features is 100.

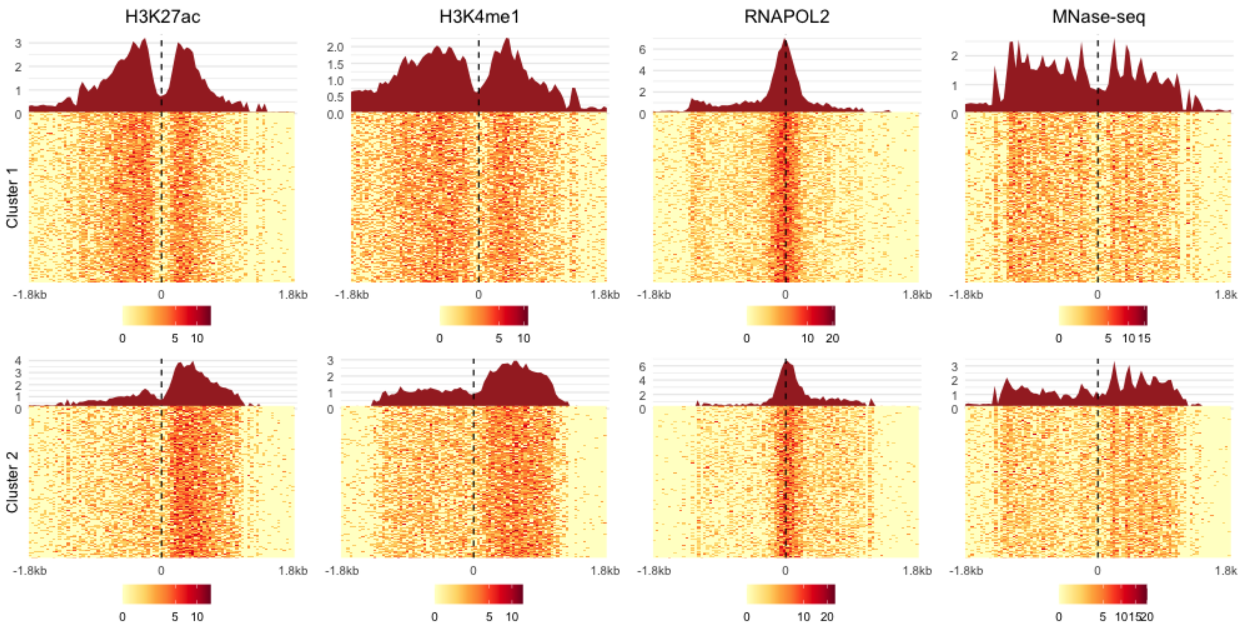

Figure S4: Simulated chromatin feature data containing two clusters and four chromatin features. All four chromatin features had coverage of 100. Heatmaps visualise the coverage signals for the first and the second cluster, and each cluster contains 500 genomic loci. The genomic loci are defined as 3.6kb regions centered at the anchor points (dashed line). On top of the heatmaps, the average *aggregate patterns* of the coverage signals are illustrated for both clusters and the four chromatin features.

The simulation of the shift and flip states is presented in Suppl. Algorithm S3. The flip states for each profile were sampled randomly with equal probabilities for the two flip states 1 and 2 (Suppl. Algorithm S3, row 3). The number of shifts states $S$ depends on the maximum absolute shift $s_{max}$ and the resolution of data $B$. The maximum shift in nucleotides was 400, resulting in $\frac{400}{40} + 1 = 21$ shift states. The simulated shifts were obtained by sampling two Poisson distributed values with parameter $\lambda$. The difference between the two Poisson distributed values follows Skellam distribution with mean 0 and variance $2 \times \lambda$. The Skellam distributed difference was used as the simulated shift $s_i$, rounded to the nearest multiplication of the bin size $B = 40$ (Suppl. Algorithm S3, rows 14 and 15) . The $\lambda$ parameter was chosen iteratively, by increasing it until the absolute value of any sampled shift exceeded the

**Algorithm S3:** Simulation of the shift and flip states

```
// Sample flip states f
for each sample i ← 1 to N do
    f_i ~Bernoulli(p = 0.5)
end
// Sample shifts s in nucleotides
s_i ← 0, i = 1, ..., N // Initialise shifts to 0
B = 40 // Resolution or bin size in bp
s_max = 400 // Absolute maximum shift in nucleotides
λ = 1000 // Poisson parameter
while abs(range(s)) < s_max do
    for each sample i ← 1 to N do
        s_1 ~Poisson(λ)
        s_2 ~Poisson(λ)
        s_i = s_1 − s_2
        s_i =round(s_i / B) × B // Round s_i to the nearest multiplication of B
        λ = λ + 100
    end
end
// Ensure that absolute shifts not exceed s_max
s[which(s ≥ s_max)] = s_max
s[which(s ≤ −s_max)] = −s_max
return Shifts s and flips f
```

allowed maximum shift $s_{max} = 400$. The simulated shift and flip states were the same in all 100 data generation instances, i.e. the same random seed was utilised to simulate the shift and flip states.

The generation of simulated data with shifted and flipped profiles is presented in Suppl. Algorithm S4. The profiles in the data to be shifted and flipped are originally of length 90. The shifts in nucleotides are converted to indexes which are used to extract the shifted data from the profiles (Suppl. Algorithm S4, row 8), and for each sample $i$, the shifted profiles of length $L_x = 50$ were extracted (Suppl. Algorithm S4, row 10). The shifted profile was flipped if the sampled flip state for the particular sample was 2.

**Algorithm S4:** Shifted and flipped simulated data generation

// The length of profiles $\mathbf{x}_i^{(m)}$ are $\frac{W_{orig}}{B} = L_{orig}$. We have used values $W_{orig} = 3.6$kb, $B = 40$bp, hence the profile lenghts were originally $L_{orig} = 90$. The final data to be clustered corresponds $W = 2$kb genomic windows, i.e. the profiles are of length $L_x = 50$.

// Shift the profiles $\mathbf{x}_i^*$

$S = \frac{2 \times s_{max}}{B} + 1$ // Number of shift states $S$, here 21

$start = \frac{W_{orig}}{2 \times B} - \frac{W}{2 \times B} + 1$ // 21

$end = \frac{W_{orig}}{2 \times B} + \frac{W}{2 \times B}$ // 70

**for** *each sample* $i \leftarrow 1$ **to** $N$ **do**

  // Shift state indexes, in range $\left[-\frac{s_{max}}{B}, \frac{s_{max}}{B}\right]$, $[-10, 10]$

  $s_i^{\text{ind}} = \frac{s_i}{B}$

  **for** *each chromatin features* $m \leftarrow 1$ **to** $M$ **do**

    $\mathbf{x}_{i,\,shifted}^{(m)} = \mathbf{x}_i^{(m)} \left[(start + s_i^{\text{ind}}) : (end + s_i^{\text{ind}})\right]$

    // Flip the profile if $f_i = 2$

    **if** $f_i = 2$ **then**

      $\mathbf{x}_{i,\,final}^{(m)} = \text{rev}(\mathbf{x}_{i,\,shifted}^{(m)})$ // Reverse the profile elements

    **else**

      $\mathbf{x}_{i,\,final}^{(m)} = \mathbf{x}_{i,\,shifted}^{(m)}$

  **end**

**end**

**return** $\mathbf{X}_{final}^*$

#### S4.5 Choosing the hyperparameter values

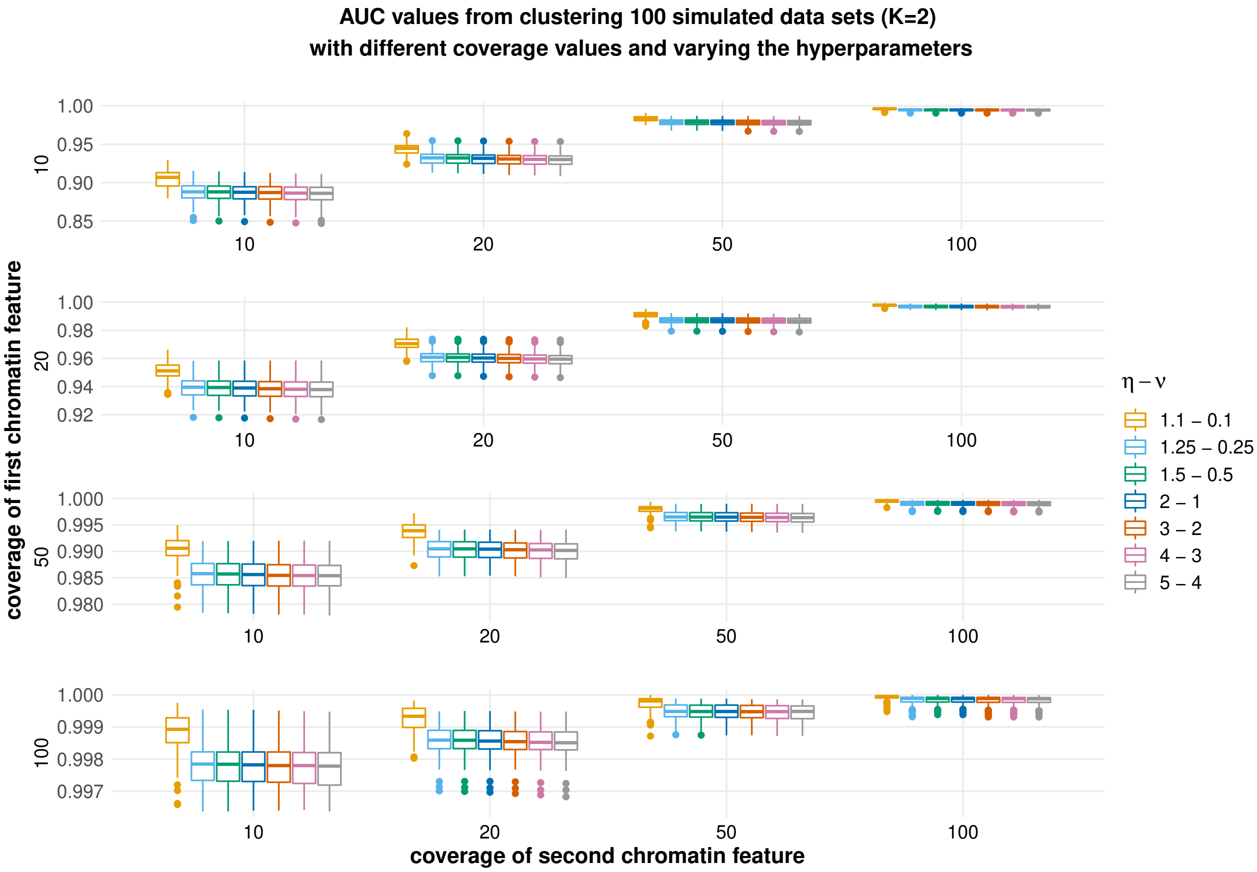

Figure S5: AUC values for clustering simulated data that contain two clusters and two chromatin features. The chromatin feature coverages varied between 10, 20, 50, and 100 and different choices for prior gamma distribution were tested. The hyperparameter values $\eta = 1.1$ and $\nu = 0.1$ resulted in the highest AUC scores. Boxplots represent results for 100 data sets.

To choose suitable prior Gamma distribution for the Dirichlet parameters $\alpha_{kj}^{(m)}$ parameterised by hyperparameters $\eta$ and $\nu$, we noticed that setting $\alpha_j = 1$ for all $j$ leads to symmetric and uniform Dirichlet distribution. Therefore, the mode of the Gamma prior was set to one. In contrast, setting $\alpha < 1$ or $\alpha > 1$ would result in a higher prior probability for sparse or dense multinomial parameters, respectively. If one lacks strong expectation for either sparse or dense multinomial distribution, the Gamma distribution with high variance would provide the model some added flexibility to choose between them. We experimented with different Gamma distributions with varying variance and concluded that the prior had little effect on the results, and the choice $\eta = 1.1$ and $\nu = 0.1$ resulting in a variance of 110 indicated the best performance in terms of AUC when clustering data containing two clusters of two chromatin features (see Suppl. Figure S5). We also demonstrated with the prior predictive check, i.e. the ancestral sampling of the data from prior hyperparameters, that the noise level generated from the prior was comparable to the noise in real data (Suppl. Figure S6). The user of the ChromDMM can verify that the prior is meaningful by running the prior predictive checks for his/her data.

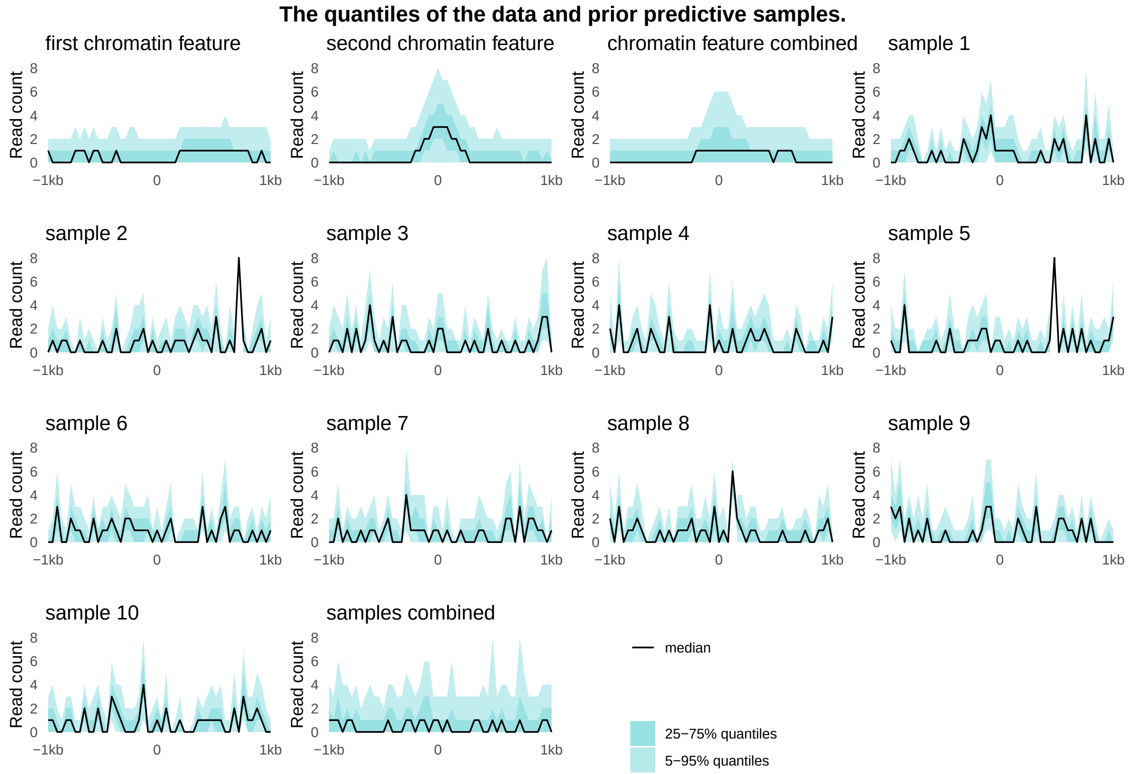

Figure S6: Prior predictive check for simulated data. The quantiles of the first and second chromatin features for 1000 genomic loci are shown together with the quantiles of combined chromatin features containing 2000 genomic loci. Both chromatin features had coverage of 50. Ancestral sampling was used to sample 10 prior predictive data sets. First, 10 $\boldsymbol{\alpha}$ vectors from the Gamma prior with hyperparameters $\eta = 1.1$ and $\nu = 0.1$ were generated. The elements of the $\boldsymbol{\alpha}$ vectors were sampled independently. Each of the 10 Dirichlet parameters were used to generate 1000 samples of multinomial parameters $\mathbf{p}_i$ from the Dirichlet distribution, and the profiles $\mathbf{x}_i$ were sampled from the multinomial distribution with coverage 50. The quantiles of these 10 prior predictive data sets together with the combined data (10000 samples) are presented in the figure.

To choose the hyperparameters $\eta_h$ and $\nu_h$ for the regularisation term $h_k^m$, prior predictive checks are unfeasible. Therefore, the simulated data were clustered by setting $\eta = 1.1$ and $\nu = 0.1$ and varying the mean of the regularisation term prior distribution between 0.1, 1, and 10 and the variance between 0.1, 1, 10, and 100. In addition, the clustering was performed without regularisation. The Gamma prior with mean 1 and variance 0.1 corresponding to hyperparameters $\eta_h = \nu_h = 10$ resulted in the best clustering performance in terms of AUC values, particularly when the coverage of both chromatin features was low (see Suppl. Figure S7). Based on these experiments, we adopted the following hyperparameter values: $\eta = 1.1$, $\nu = 0.1$, $\eta_h = 10$, and $\nu_h = 10$.

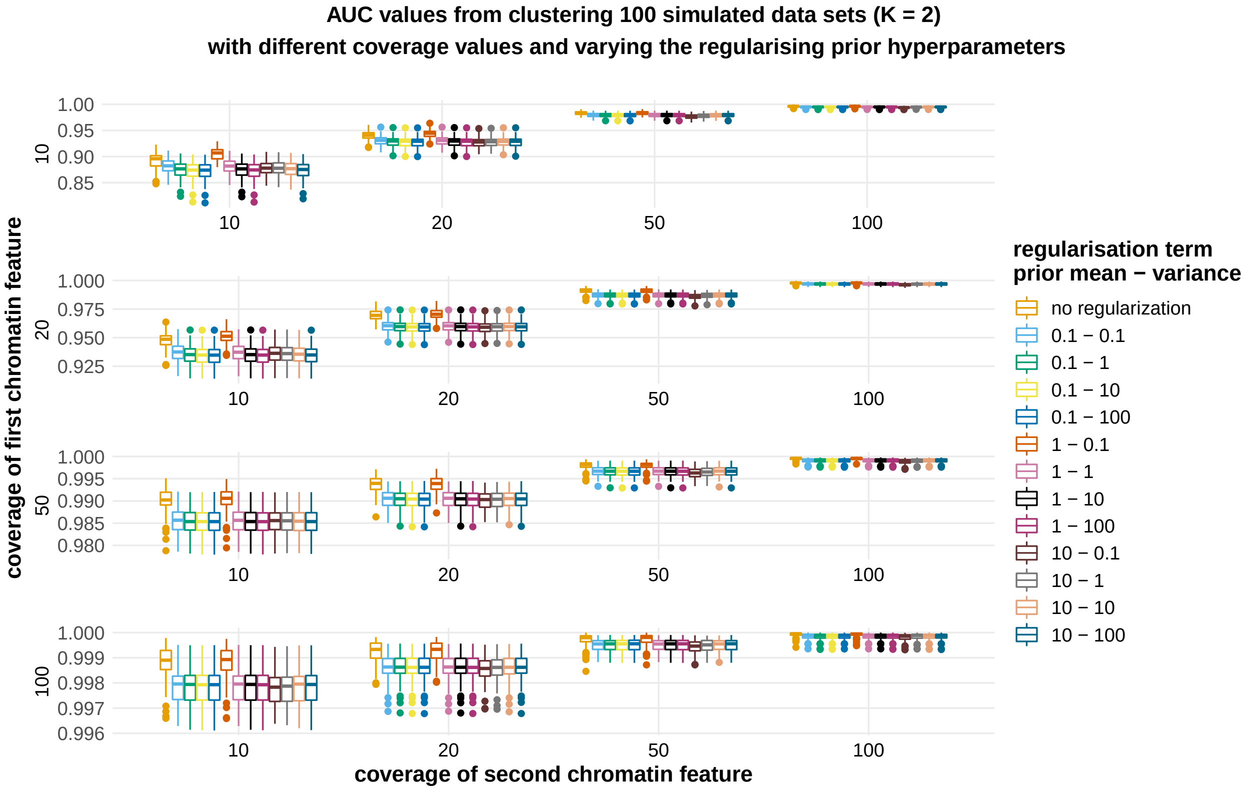

Figure S7: AUC values for clustering simulated data sets containing two clusters and two chromatin features. The chromatin feature coverages varied between 10, 20, 50, and 100 and different choices for the regularisation term gamma prior distribution were tested. The mean of the regularising prior varied between 0.1, 1, and 10 and the variance varied between 0.1, 1, 10, and 100. The regularisation term hyperparameter values $\eta_h = 10$ and $\nu_h = 10$ corresponding to mean 1 and variance of 0.1 resulted in highest AUC scores especially when coverage for both chromatin features were low. The values for the other hyperparameters were $\eta = 1.1$ and $\nu = 0.1$. Boxplots present results for 100 data sets. .

In addition to the choice of hyperparameters for ChromDMM, the performance of ChromDMM, ChIP-Partitioning and SPar-K depend on the initialisation strategy and numerical parameters including the maximum number of EM iterations in ChromDMM and ChIP-Partitioning, the maximum number of iterations in ChromDMM BFGS step, the number of k-means iterations in SPar-K, and the numerical tolerances when checking the converge of the iterations. The experimentation with these options for ChIP-Partitioning and SPar-K is out of the scope of our work, so we resorted to default options.

Mixture models are known for their convergence to a local mode. A straightforward strategy to explore the search space is to repeat the model inference with random initial starting points. In the simulated data experiments, ChromDMM was fitted using 10 random initialisations and the best fit according to the lower bound was selected as the final model. In the real enhancer clustering, the number of repetitions was 20. The default options for ChIP-Partitioning and SPar-K do not include such repetitions.

### S5 Supplementary References

Benjamini, Y. and Hochberg, Y. (1995). Controlling the false discovery rate: a practical and powerful approach to multiple testing. *Journal of the Royal statistical society: series B (Methodological)*, **57**(1), 289–300.

Davis, C. A., Hitz, B. C., Sloan, C. A., Chan, E. T., Davidson, J. M., Gabdank, I., Hilton, J. A., Jain, K., Baymuradov, U. K., Narayanan, A. K., *et al.* (2018). The encyclopedia of dna elements (encode): data portal update. *Nucleic acids research*, **46**(D1), D794–D801.

Gu, Z., Eils, R., and Schlesner, M. (2016). Complex heatmaps reveal patterns and correlations in multidimensional genomic data. *Bioinformatics*, **32**(18), 2847–2849.

Kharchenko, P. V., Tolstorukov, M. Y., and Park, P. J. (2008). Design and analysis of ChIP-seq experiments for DNA-binding proteins. *Nature Biotechnology*, **26**(12), 1351–1359.

Landt, S. G., Marinov, G. K., Kundaje, A., Kheradpour, P., Pauli, F., Batzoglou, S., Bernstein, B. E., Bickel, P., Brown, J. B., Cayting, P., Chen, Y., DeSalvo, G., Epstein, C., Fisher-Aylor, K. I., Euskirchen, G., Gerstein, M., Gertz, J., Hartemink, A. J., Hoffman, M. M., Iyer, V. R., Jung, Y. L., Karmakar, S., Kellis, M., Kharchenko, P. V., Li, Q., Liu, T., Liu, X. S., Ma, L., Milosavljevic, A., Myers, R. M., Park, P. J., Pazin, M. J., Perry, M. D., Raha, D., Reddy, T. E., Rozowsky, J., Shoresh, N., Sidow, A., Slattery, M., Stamatoyannopoulos, J. A., Tolstorukov, M. Y., White, K. P., Xi, S., Farnham, P. J., Lieb, J. D., Wold, B. J., and Snyder, M. (2012). ChIP-seq guidelines and practices of the ENCODE and modENCODE consortia. *Genome Research*, **22**(9), 1813–1831.

Langmead, B. and Salzberg, S. L. (2012). Fast gapped-read alignment with Bowtie 2. *Nature Methods*, **9**(4), 357–359.

Le Martelot, G., Canella, D., Symul, L., Migliavacca, E., Gilardi, F., Liechti, R., Martin, O., Harshman, K., Delorenzi, M., Desvergne, B., Herr, W., Deplancke, B., Schibler, U., Rougemont, J., Guex, N., Hernandez, N., and Naef, F. (2012). Genome-Wide RNA Polymerase II Profiles and RNA Accumulation Reveal Kinetics of Transcription and Associated Epigenetic Changes During Diurnal Cycles. *PLoS Biology*, **10**(11), 1–16.

Li, Q., Brown, J. B., Huang, H., and Bickel, P. J. (2011). Measuring reproducibility of high-throughput experiments. *Annals of Applied Statistics*, **5**(3), 1752–1779.

MacKay, D. J. (2003). *Information theory, inference, and learning algorithms*. Cambridge University Press, New York, USA.

Marx, V. (2017). How to deduplicate PCR. *Nature Methods*, **14**(5), 473–476.

Mi, H., Ebert, D., Muruganujan, A., Mills, C., Albou, L.-P., Mushayamaha, T., and Thomas, P. D. (2020). PANTHER version 16: a revised family classification, tree-based classification tool, enhancer regions and extensive API. *Nucleic Acids Research*, **49**(D1), D394–D403.

Mosimann, J. E. (1962). On the compound multinomial distribution, the multivariate $\beta$-distribution, and correlations among proportions. *Biometrika*, **49**(1/2), 65–82.

Osmala, M. and Lähdesmäki, H. (2020). Enhancer prediction in the human genome by probabilistic modelling of the chromatin feature patterns. *BMC bioinformatics*, **21**(1), 1–37.

The ENCODE Project Consortium (2012). An integrated encyclopedia of DNA elements in the human genome. *Nature*, **489**(7414), 57–74.

Wang, J., Zhuang, J., Iyer, S., Lin, X., Whitfield, T. W., Greven, M. C., Pierce, B. G., Dong, X., Kundaje, A., Cheng, Y., *et al.* (2012). Sequence features and chromatin structure around the genomic regions bound by 119 human transcription factors. *Genome research*, **22**(9), 1798–1812.

#### S6 Supplementary Figures

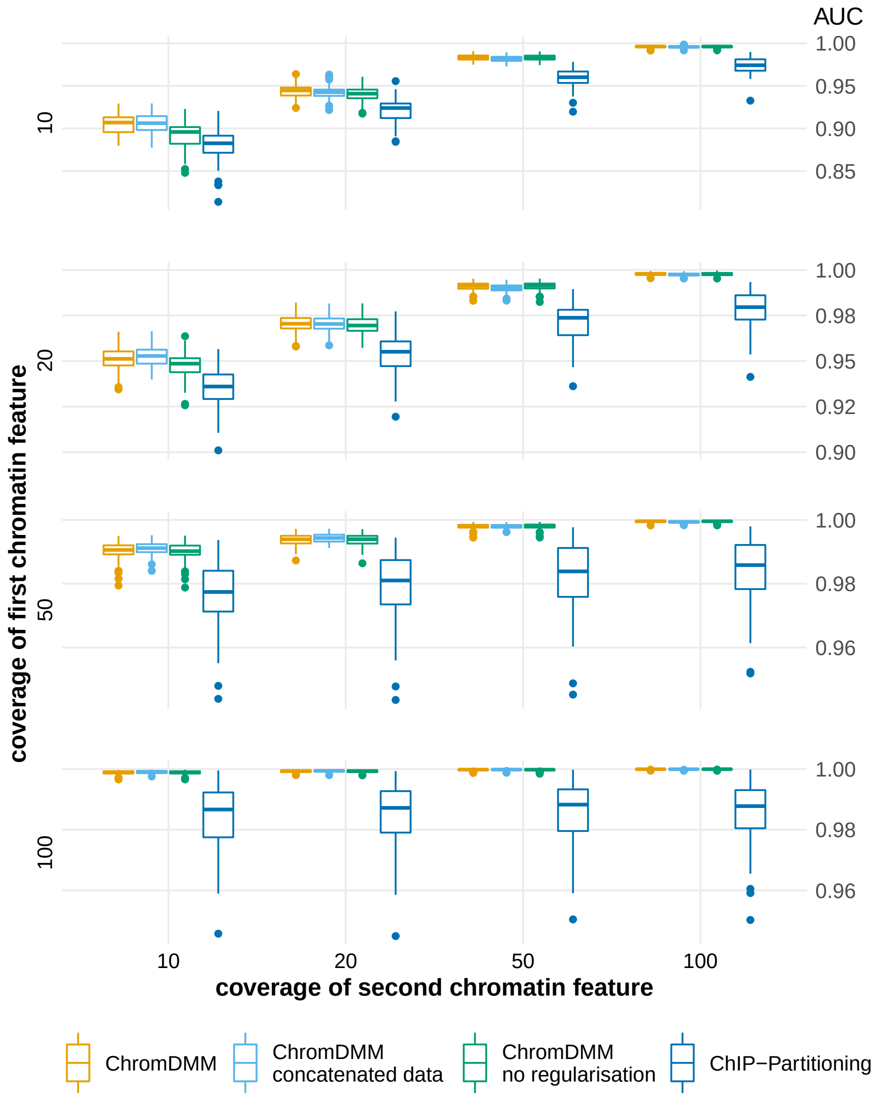

Figure S8: AUC values for clustering simulated data sets that contain two clusters and two chromatin features (H3K4me1 and RNA POL II). The chromatin feature coverages were varied between 10, 20, 50, and 100. Boxplots represent results for 100 data sets.

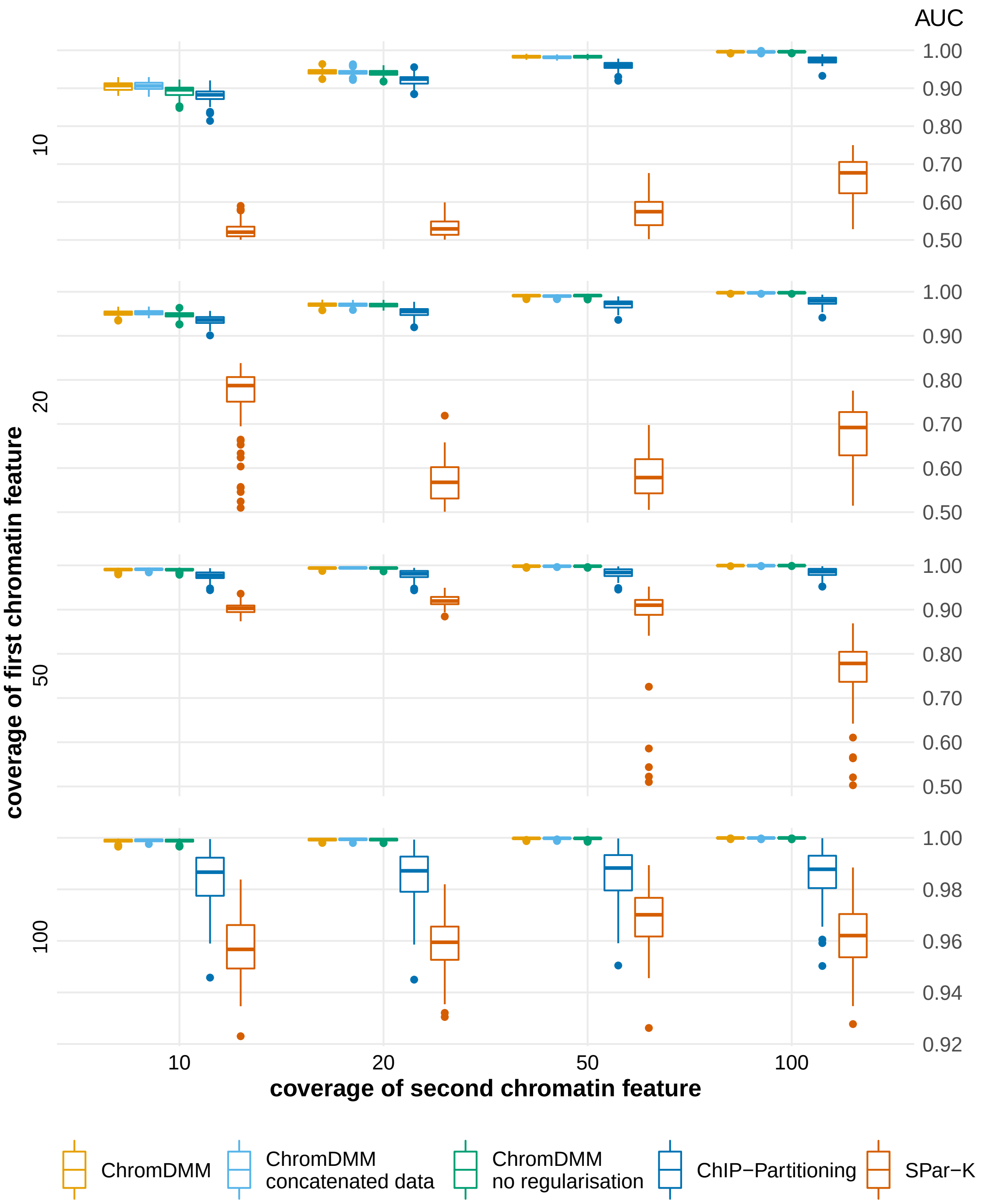

Figure S9: AUC values for clustering simulated data sets that contain two clusters and two chromatin features (H3K4me1 and RNA POL II). The results for ChromDMM and ChIP-Partitioning are the same as in Suppl. Figure S8, but the Figure presents results also for SPar-K. The chromatin feature coverages were varied between 10, 20, 50, and 100. Boxplots represent results for 100 data sets.

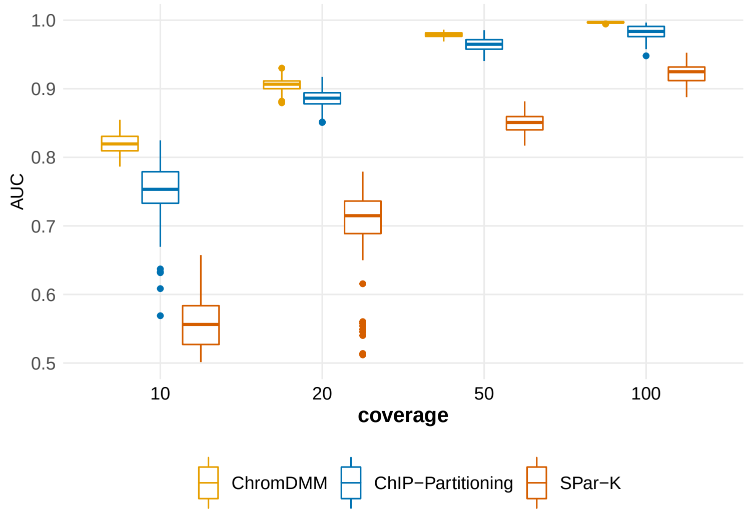

Figure S10: AUC values for clustering simulated data sets that contain two clusters and one chromatin feature. The chromatin feature coverage varied between 10, 20, 50, and 100. Boxplots represent results for 100 data sets. ChromDMM perfoms best compared to ChIP-Partitioning and SPar-K.

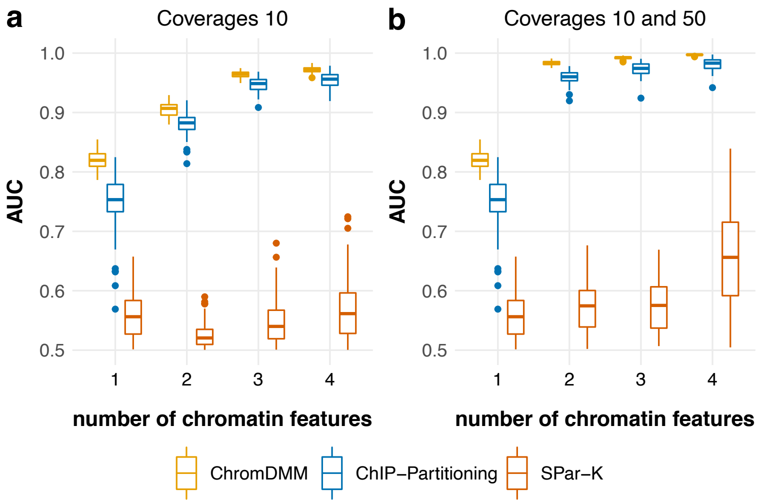

Figure S11: AUC values for clustering simulated data sets that contain two clusters and varying the number of chromatin features between 1 and 4. The data set 1 consisted of chromatin feature H3K4me1, the set 2 consisted of H3K4me1 and RNA POL II, the set 3 consisted of H3K27ac, H3K4me1, and RNA POL II, and the set 4 consisted of H3K27ac, H3K4me1, RNA POL II, and MNase-seq. In Figure a), the coverage of all chromatin features were 10. In Figure b), the coverage of the first two chromatin features (H3K4me1 and H3K27ac) were 10 and the coverage of the last two chromatin features (RNA POL II and MNase-seq) were 50. Boxplots represent results for 100 data sets.

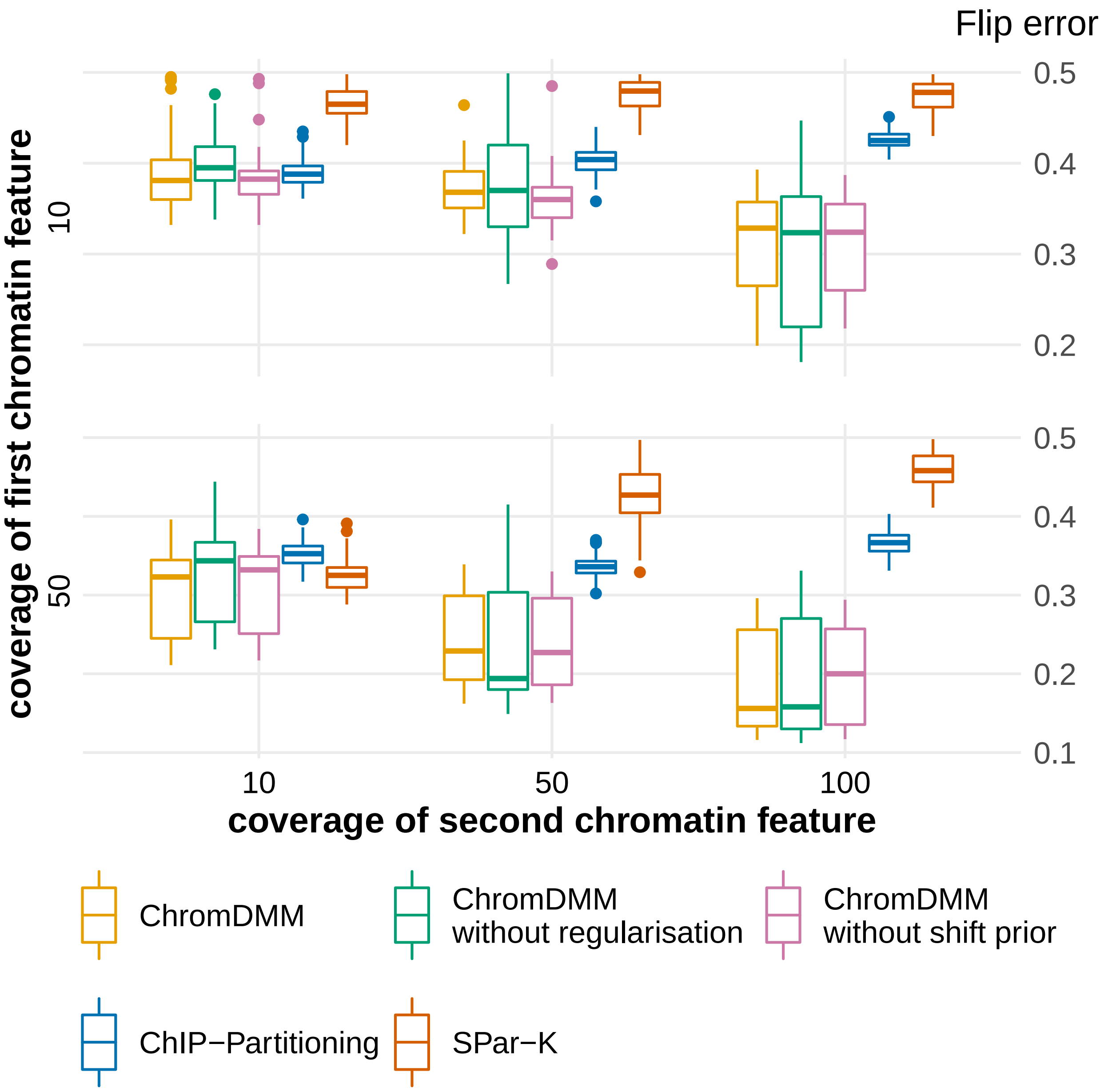

Figure S12: Proportions of incorrect flip states for clustering simulated data that contain two clusters, two chromatin features and randomly sampled shift and flip states. The chromatin feature coverages varied between 10, 50, and 100. Results are shown for ChromDMM, ChIP-Partitioning, and SPar-K. ChromDMM was inferred also with the uniform shift state prior and without the regularisation term. Boxplots represent results for 100 data sets.

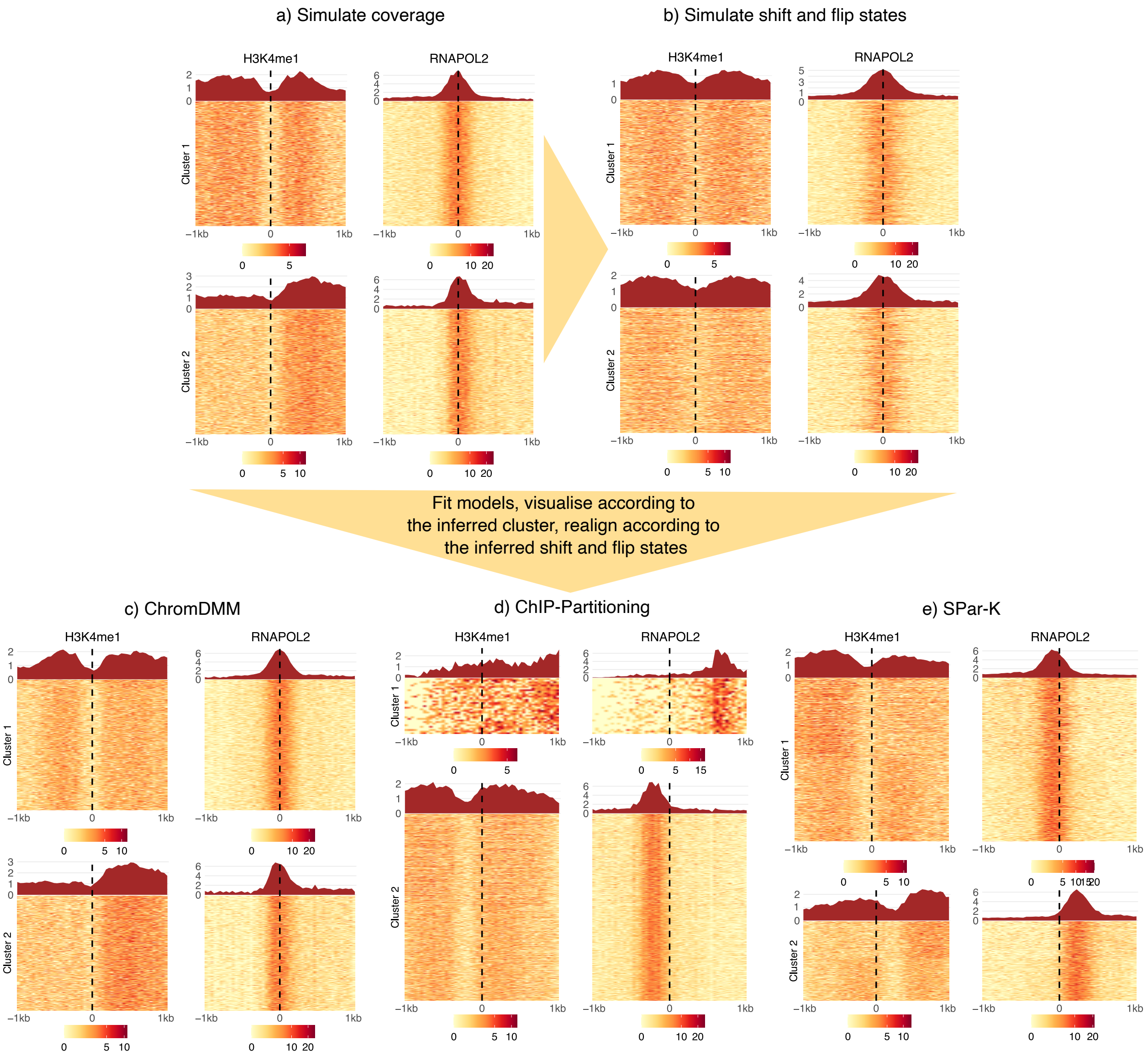

Figure S13: Figure a) presents an example of simulated data containing two clusters and two chromatin features. The coverage for both chromatin features is 100. Figure b) presents shifted and flipped simulated data to illustrate that the asymmetry of the profiles and fine structure in the patterns disappear. The methods were fitted to the data in shown in Figure b) to infer the most probably clusters, flip states, and shift states. The results are visualised for c) ChromDMM, d) ChIP-Partitioning and e) SPar-K. ChIP-Partitioning assigned 35 samples to cluster 1 and the rest to cluster 2, the heights of heatmaps in d) are not proportional to the cluster sizes. ChromDMM can clearly infer the correct clusters and register the profiles to reveal the original patterns, whereas ChIP-Partitioning and SPar-K have difficulties even to cluster the data correctly. In addition, ChromDMM retains the unimodal peaks and valleys between two-modal peaks in the middle of the cluster patterns, whereas ChIP-Partitioning and SPar-K drift the patterns left or right from the original profile centers.

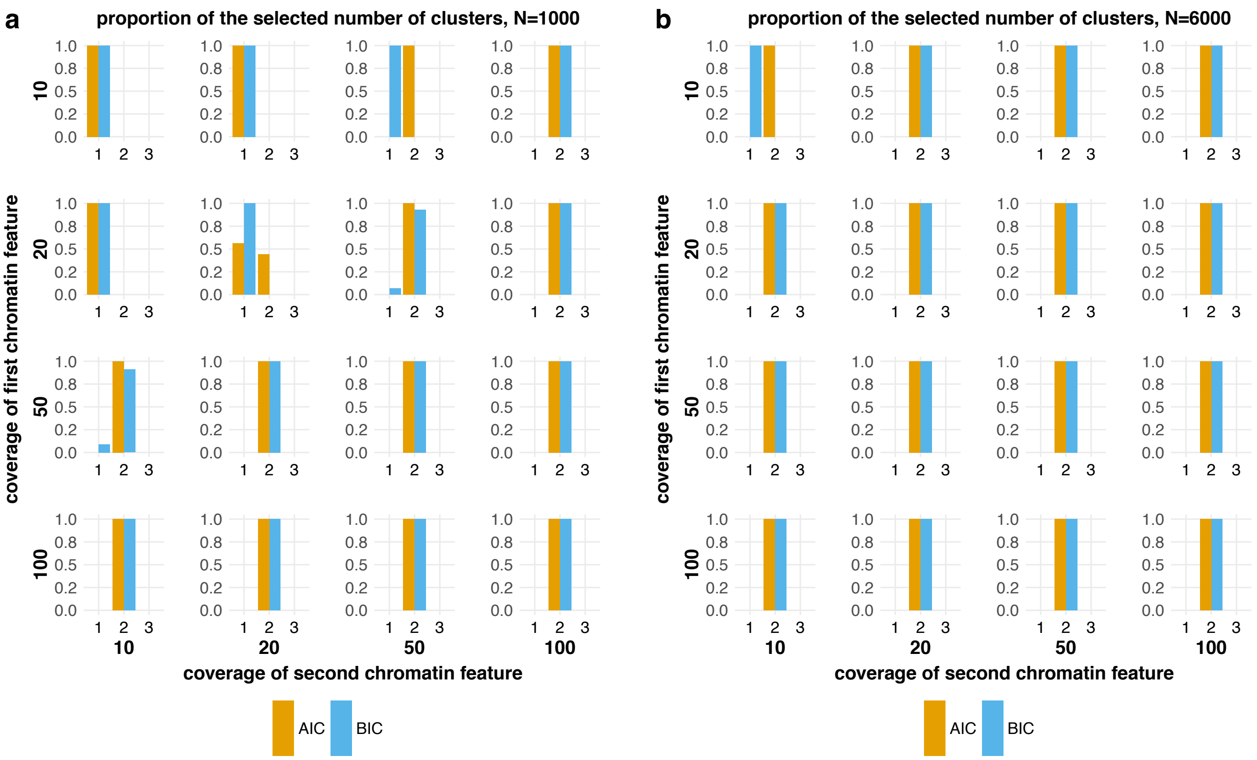

Figure S14: The proportion of number of clusters selected by AIC and BIC. The simulated data contained two clusters and two chromatin features (H3K4me1 and RNA POL II). The chromatin feature coverages were varied (10,20,50 and 100), and for each combination, 100 simulated data sets were generated and clustered with ChromDMM without the inference of the shift and flip states. Figure a) presents the results on data containing 1000 samples and Figure b) presents the results on data containing 6000 samples. Increasing the chromatin feature coverage as well as increasing the number of genomic loci being clustered improves the selection of correct number of clusters 2 by AIC and BIC. BIC has tendency to underestimate the number of clusters compared to AIC.

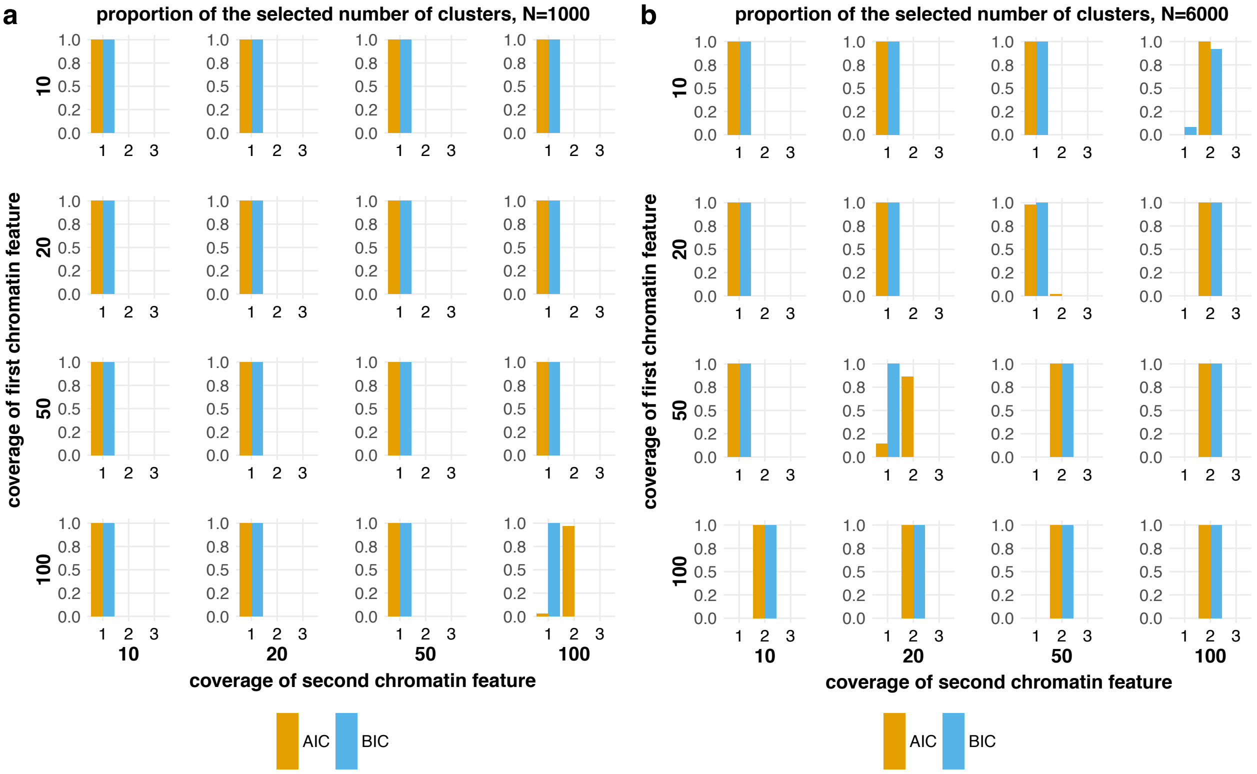

Figure S15: The proportion of number of clusters selected by AIC and BIC. The simulated data contained two clusters and two chromatin features (H3K4me1 and RNA POL II). The chromatin feature coverages were varied (10,20,50 and 100), and for each combination, 100 simulated shifted and flipped data sets were generated and clustered with ChromDMM with the inference of the shift and flip states. Figure a) presents the results on data containing 1000 samples and Figure b) presents the results on data containing 6000 samples. As in Figure S14, increasing the chromatin feature coverage as well as increasing the number of genomic loci being clustered improves the selection of correct number of clusters 2 by AIC and BIC. However, inferring the correct number of clusters by AIC and BIC by ChromDMM with shifting and flipping feature is a more challenging task. The more complex model with the number of parameters increasing as the function of the cluster number are likely penalised heavily.

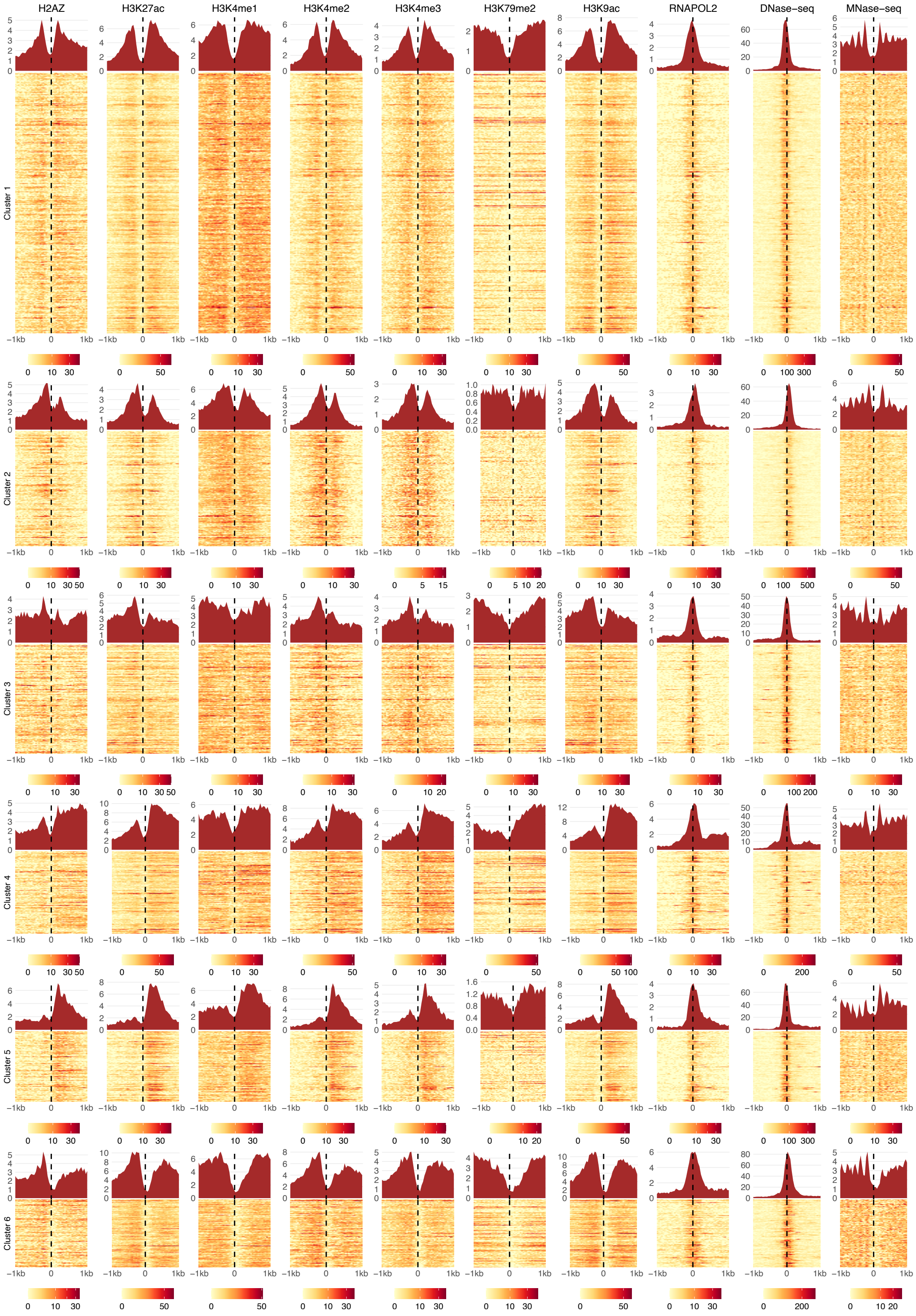

Figure S16: Six enhancer clusters identified by ChromDMM. The coverage signals of individual enhancers assigned to the clusters are visualised as heatmaps. The aggregate patterns are visualised on top of the heatmaps.

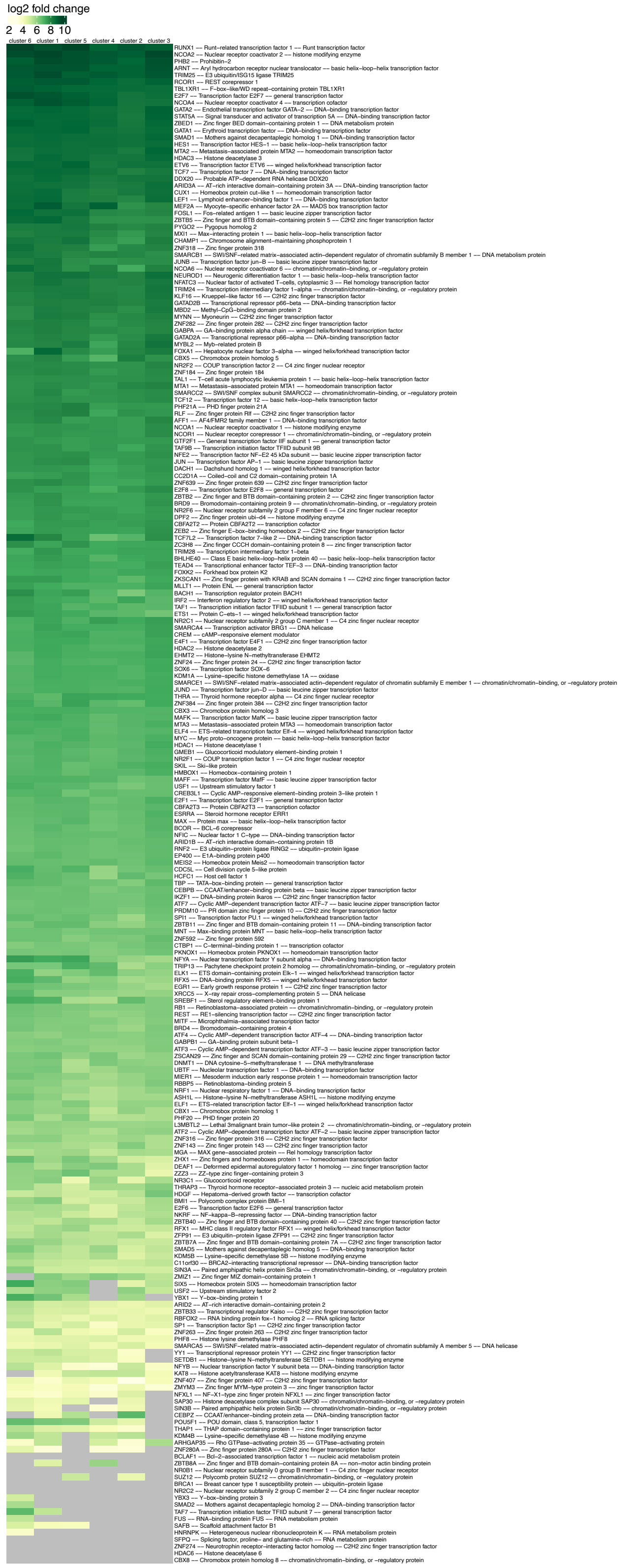

Figure S17: The fold enrichment of 220 TRFs at the enhancer clusters identified by ChromDMM.

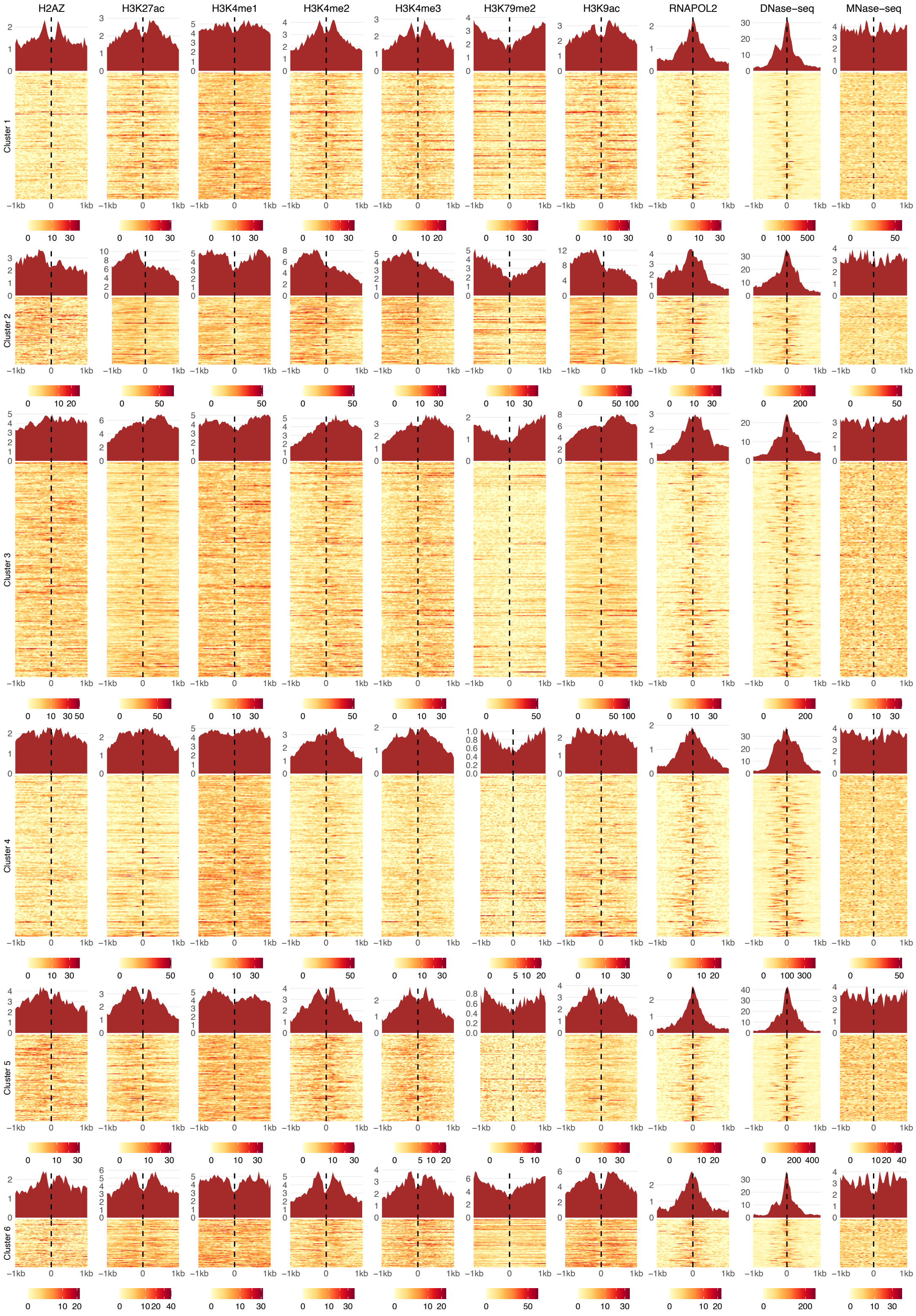

Figure S18: Six enhancer clusters identified by ChIP-Partitioning. The coverage signals of individual enhancers assigned to the clusters are visualised as heatmaps. The aggregate patterns are visualised on top of the heatmaps.

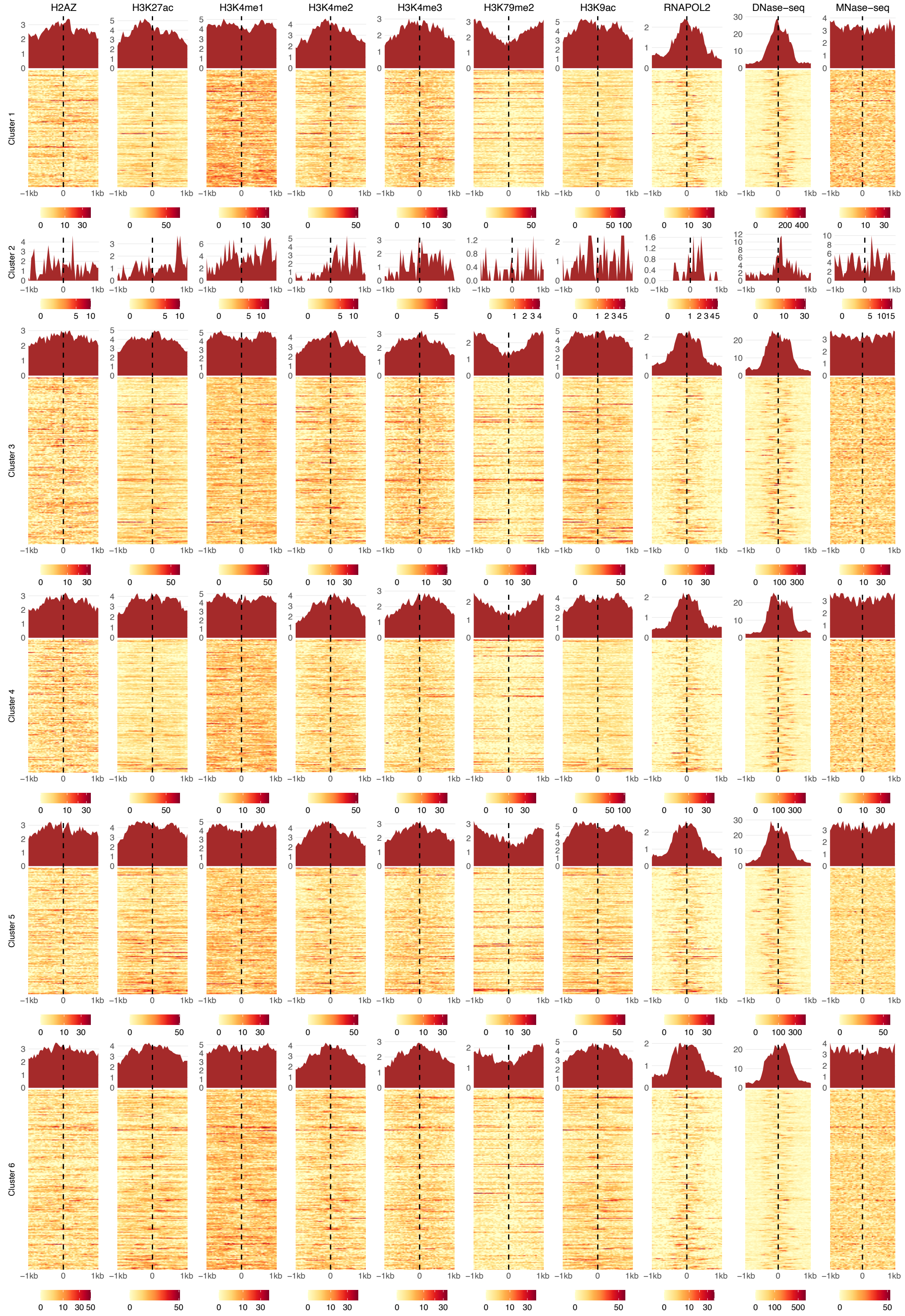

Figure S19: Six enhancer clusters identified by SPar-K. The coverage signals of individual enhancers assigned to the clusters are visualised as heatmaps. The aggregate patterns are visualised on top of the heatmaps. The second cluster includes only 3 genomic regions, hence the heatmap is not visible.

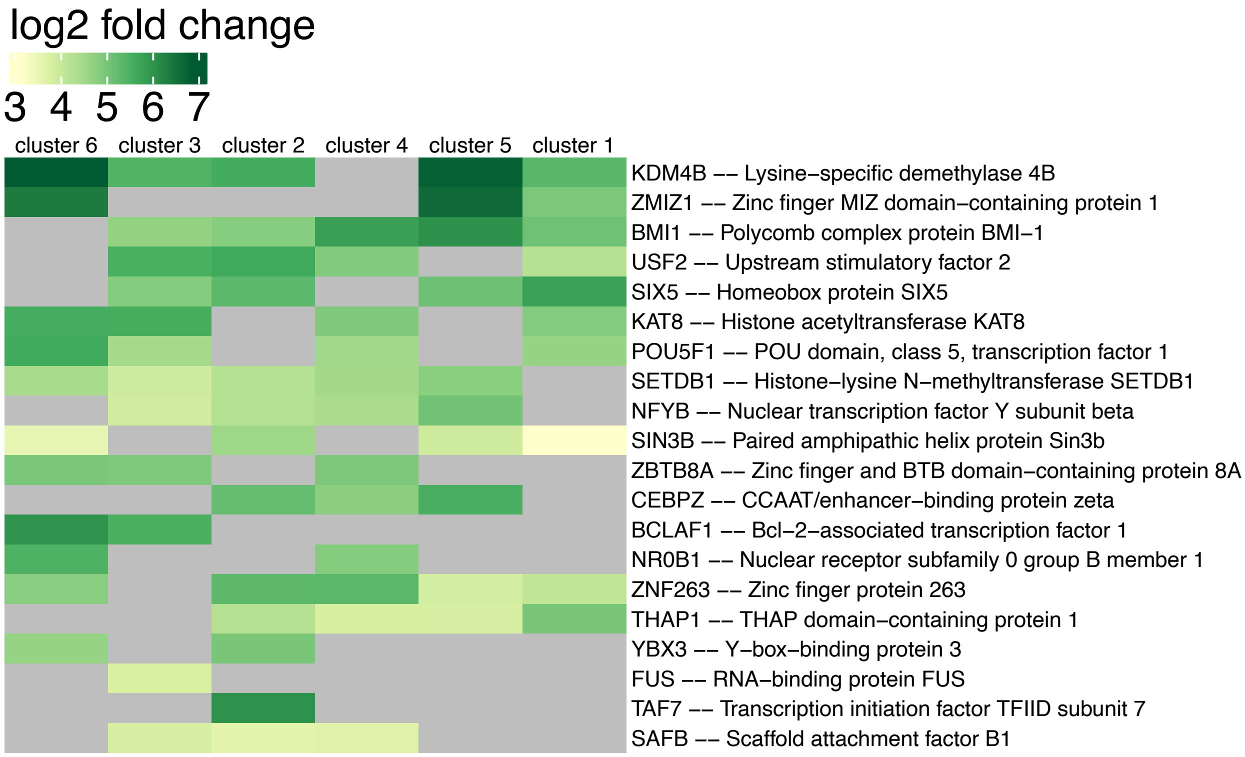

Figure S20: The fold enrichment of TRFs at the enhancer clusters identified by ChIP-Partitioning. The fold enrichments correponding to q-value larger than 0.01 were masked out from the heatmap.

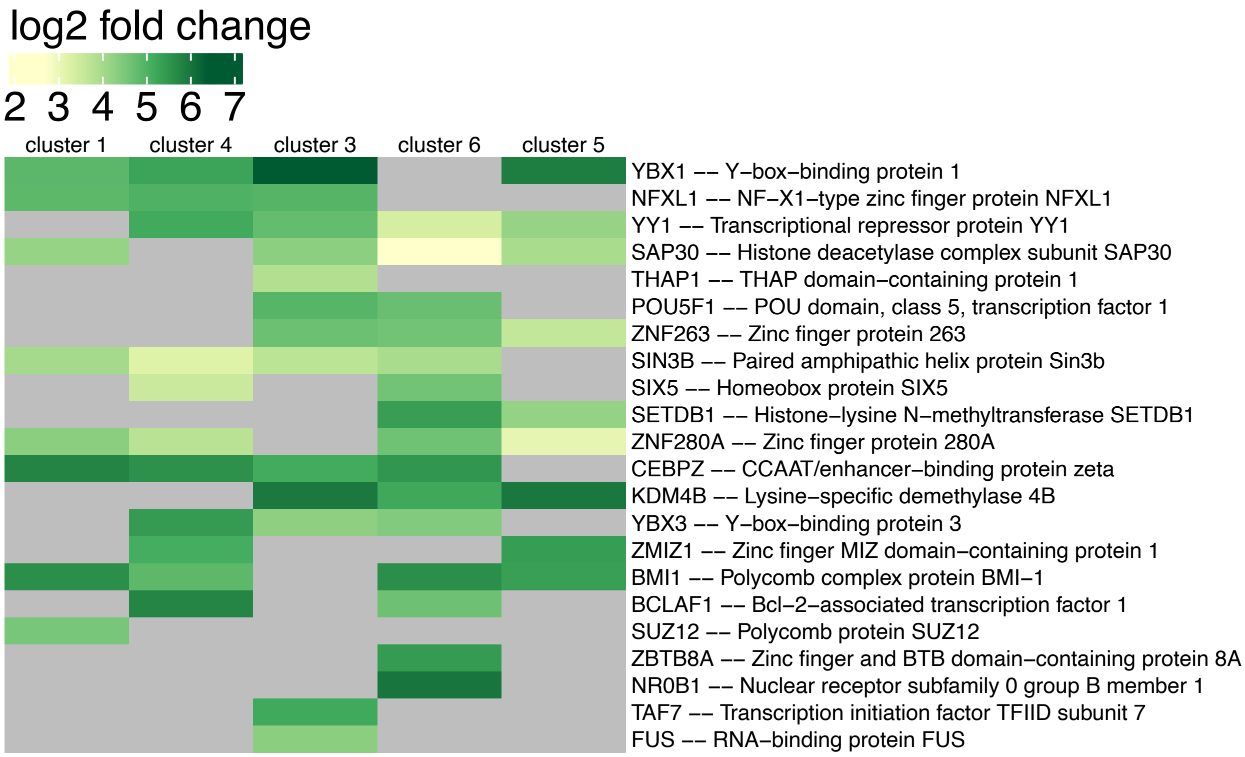

Figure S21: The fold enrichment of TRFs at the enhancer clusters identified by SPar-K. The fold enrichments correponding to q-value larger than 0.01 were masked out from the heatmap. Cluster 2 contained only three enhancers and the enrichment for most of the TRFs was not significant, hence the TRF enrichments for that cluster are not visualised.
